# Familial Parkinson’s Disease Mutations Converge on Alternative Splicing Programs Linked to Lysosomal, Autophagic and Vesicle Trafficking Pathways

**DOI:** 10.1101/2024.02.28.582420

**Authors:** Yeon J. Lee, Khaja Syed, Oriol Busquets, Hanqin Li, Jesse Dunnack, Helen S. Bateup, Frank Soldner, Dirk Hockemeyer, Donald C. Rio

## Abstract

Parkinson’s disease (PD) arises from genetic and environmental factors. Human genetics has identified mutations in ∼20 inherited familial genes linked to monogenic forms of PD. To investigate the how individual familial PD mutations might affect pre-mRNA splicing patterns, human pluripotent stem cells (hPSCs) carrying 12 distinct familial PD mutations were differentiated into midbrain lineage cells, including dopaminergic (mDA) neurons. Using available snRNA-seq data from human brain substantia nigra pars compacta (SNc), we curated a set of ∼ 2,250 genes expressed in DA neurons, where >90% of these genes were expressed in our differentiated hPSC cultures. We used the iSCORE-PD collection of edited hPSCs carrying pathogenic PD mutations in the *PRKN*, *SNCA*, *LRRK2*, *PINK1*, *DNAJC6*, *FBXO7*, *SYNJ1*, DJ1, *VPS13C*, *ATP13A2* and *GBA1* genes and carried out deep Illumina RNA-seq from these differentiated cultures to carry out pre-mRNA splicing analyses. Altered splicing patterns in familial PD mutant cells were categorized into five distinct functional classes: autosomal-dominant drivers (*SNCA, LRRK2*), mitochondrial quality control (*PRKN, PINK1, PARK7*), endolysosomal/vesicular trafficking (*ATP13A2, DNAJC6, FBXO7, SYNJ1*), lysosomal dysfunction (*GBA1*), and lipid transport/cellular stress pathways (*VPS13C*). Interestingly, we found common pre-mRNA splicing changes for the curated set of SN dopaminergic neuronal genes in the mutant cells, including for *SYNJ1*, itself a familial PD risk gene linked to dopamine release. These PD mutation-specific pre-mRNA isoform changes may serve as both diagnostic biomarkers for familial PD-associated genotypes and be promising therapeutic targets.

## Introduction

Parkinson’s disease (PD) affects 2-3% of individuals over 65 years of age and is the second most common neurodegenerative disease (Poewe et al. 2017; Aarsland et al. 2021). PD is characterized by severe deterioration of motor function caused by loss of dopaminergic neurons in the substantia nigra pars compacta (SNc) and is accompanied by deficient dopamine release in the caudate/putamen (Kouli et al. 2018). A hallmark of PD is the intracellular aggregation of the protein α-synuclein into histologically detectable inclusions called Lewy bodies (Poewe et al. 2017; Aarsland et al. 2021). Cells outside the brain, including cells throughout the central and peripheral autonomic nervous system are also affected as the disease progresses (Poewe et al. 2017; Mendoza-Velasquez et al. 2019; Chen et al. 2020).

Lewy body diseases include dementia with Lewy body disease (DLB) and Parkinson’s disease with dementia (PDD). The pathological hallmark of these disorders is the presence of Lewy bodies, intracellular inclusions composed largely of aggregated α-synuclein, in affected regions of the brain. Key insights into disease mechanisms came from human genetic studies that have identified more than 20 genes that cause familial forms of PD (Poewe et al. 2017; Aarsland et al. 2021). Many of these genes, including *SNCA*, which encodes α-synuclein itself, point to defects in the lysosomal-autophagy system (LAS) (Winslow et al. 2010; Navarro-Romero et al. 2020). α-synuclein oligomers inhibit macro-autophagy, contributing to protein accumulation in PD (Winslow and Rubinsztein 2011; Fellner et al. 2021). Similarly, *LRRK2* mutations linked to familial PD impair the function of lysosome-associated membrane protein (LAMP)-positive compartments and cilia (O’Hara et al. 2020; Khan et al. 2021; Alessi and Pfeffer 2024), potentially by disrupting autophagy within these structures or affecting autophagosome transport (Boecker et al. 2021). Other mutant genes linked to PD affect lysosome or mitochondrial function, as well as lipid and protein sorting (Plotegher and Duchen 2017; Navarro-Romero et al. 2020). The proposed underlying mechanisms include defects in proteostasis, calcium homeostasis, mitochondrial and autophagy dysfunction, oxidative stress, altered axonal transport and neuroinflammation (Lehtonen et al. 2019; Wilson et al. 2023).

Dysregulated alternative pre-mRNA splicing is emerging as a new and common mechanism implicated in the pathogenesis of neurodegenerative diseases such as Alzheimer’s disease (AD), PD and spinal muscular atrophy (SMA) (Li et al. 2021; Nikom and Zheng 2023; Rogalska et al. 2023). We have developed a platform to study the effects of familial, hereditary PD mutations on pre-mRNA splicing patterns and gene expression changes in *in vitro* cultured human pluripotent stem cell (hPSC)-derived dopaminergic (DA) neurons. Previously a genome-editing platform using Cas9 and prime editing was developed to engineer many of the most common familial PD mutations into hPSCs (Li et al. 2022b; Busquets et al. 2026). Using cells from this iSCORE-PD collection, we examined the effects on pre-mRNA splicing patterns in cells carrying disease-causing or high-risk variants in genes linked to PD including *PRKN, SNCA, LRRK2, PINK1, DNAJC6, FBXO7, SYNJ1, PARK7/DJ1, VPS13C, ATP13A2 and GBA1* (Polymeropoulos et al. 1997; Kruger et al. 1998; Sasaki et al. 2004; Isaacs et al. 2017; Busquets et al. 2026). Following differentiation of multiple independent cell clones carrying these mutations into midbrain lineage cells, including DA neurons (Syed et al. 2026), we carried out deep bulk short-read RNA sequencing (RNA-seq) using the Illumina platform to determine pre-mRNA patterns, using an annotation-free method called the <u>J</u>unction <u>U</u>sage <u>M</u>odel (JUM) for comparison of splicing patterns (Wang and Rio 2018) between the mutant and edited wild type (EWT) control cells (isogenic clones subjected to editing reagents in parallel without acquiring the target mutations). We compared our datasets to a curated set of ∼ 2,250 DA neuron-specific genes from snRNA-seq data expressed in the human substantia nigra (SN), which is the brain region where vulnerable neurons degenerate in PD (Wang et al. 2024) to detect differentially spliced (DS) transcripts. Recognizing that familial PD encompasses distinct molecular etiologies that may not operate through a single unified pathway, we separated the 11 mutant cell lines into five categories based on the genetic inheritance patterns, cellular pathways involved and clinical presentation. This sub-categorization allowed us to query pathway-specific alterations as well as broad disease signatures across our cellular models. We found common splicing changes across multiple familial PD mutant cells among the set of 2,250 DA neuron-specific genes (Wang et al. 2024), including in *SYNJ1*, itself a PD risk gene linked to DA release. Taken together, our data support the idea that the pre-mRNA splicing pattern defects we observe in hPSC-derived midbrain DA neurons (mDA) *in vitro* are relevant to the brain region affected in PD. These distinct transcript isoforms should serve as useful biomarkers and might be amenable to manipulation using RNA therapeutics (Anthony 2022; Li et al. 2022a; Rogalska et al. 2023).

## Results

### Rationale for this study

Many neurodegenerative diseases are linked to alterations in RNA processing pathways, e.g. amyotrophic lateral sclerosis (ALS; (van Blitterswijk and Landers 2010)), Alzheimer’s disease (AD; (Jakubauskiene and Kanopka 2021)) and multiple sclerosis (MS; (Reder et al. 2024)). In fact, RNA splicing patterns altered in *TDP-43*-linked ALS have been found to overlap splicing patterns found in PD *LRRK2* G2019S mutants generating novel peptides (Brenton et al. 2025). Because PD has been linked to multiple independent familial mutations, we sought to test whether DA neurons derived from hPSCs carrying these mutations exhibited any pre-mRNA splicing alterations. While single-cell RNA-seq data can define cell types, accurately determining altered pre-mRNA splicing patterns from these shallow sequencing data is difficult. Therefore, following differentiation of the iSCORE-PD hPSC collection with different familial PD mutations (Busquets et al. 2026; Syed et al. 2026), we carried out deep bulk RNA-sequencing to analyze pre-mRNA splicing patterns using JUM software (Wang and Rio 2018). The derivation and differentiation of the Isogenic Stem Cell Collection to Research Parkinson’s Disease (iSCORE-PD) lines into midbrain dopaminergic (mDA) cells have been detailed previously (Busquets et al. 2026). These cultures typically yield a variable percentage of dopaminergic (DA) neurons, ranging from 15% to 60% (Syed et al. 2026).. To find DA neuron-relevant transcripts to characterize at the splicing level, we curated a set of ∼2,250 genes that were found to be expressed in human SN DA neurons from single-nucleus RNA-seq datasets (Wang et al. 2024) (Supplemental Table 1). Greater than 90% of these genes were expressed in our bulk RNA-seq datasets.

Our findings were categorized into five groups based on the genetics and known functions of the mutant familial PD genes and their associated cellular pathways. The first category focuses on autosomal-dominant factors central to PD pathology, including *SNCA* and *LRRK2*. The second category encompasses autosomal-recessive genes involved primarily in mitochondrial quality control and mitophagy, including *PRKN, PINK1*, and *PARK7/DJ-1*. The third category comprises genes associated with atypical or complex parkinsonism and endolysosomal or vesicular-trafficking pathways, including *ATP13A2, DNAJC6, FBXO7,* and *SYNJ1*. Finally, *GBA1* and *VPS13C* represent distinct genetic contributors linked to lysosomal dysfunction, lipid/mitochondrial homeostasis and related cellular stress pathways (Lesage et al. 2016; Trevisan et al. 2024). Detailed descriptions of the pre-mRNA splicing (DS) patterns identified within the curated set of DA neuron–expressed genes for each mutant hPSC line are provided below. Notably, some of these splicing pattern alterations directly affected established PD genes, including *SYNJ1*. Additionally, we also carried out gene expression and splicing analyses on the bulk RNA-seq datasets (which are shown in the Supplemental Figures and Tables, see below). Importantly, we note that in both cases the transcripts whose splicing patterns are altered do not overlap to a large extent with those whose expression levels change in the mutant versus wild type cells.

#### Differentiation of dopaminergic (DA) neurons from human pluripotent stem cells (hPSCs) carrying familial PD mutations

To study the effects of PD-linked mutations on splicing and gene expression, we differentiated the entire iSCORE-PD collection (Busquets et al. 2026) into mDA cells. Multiple clonally independent isogenic stem cell lines were generated carrying common familial PD mutations including *PRKN* (X3DEL)*, SNCA* (A30P and A53T*), LRRK2* (G2019S), *PINK1* (Q129X)*, DNAJC6* (FS)*, FBXO7* (R498X)*, SYNJ1* (R258Q)*, DJ1* (X1_5DEL)*, VPS13C* (W395C)*, ATP13A2* (FS) *and GBA1* (IVS2) using CRISPR *Cas9* or prime editing techniques, as previously reported (Figure 1A; (Li et al. 2022b; Busquets et al. 2026). hPSCs were differentiated into mDA cells, which are the primary cell type that degenerates in PD using a modified version of a reported protocol (Kim et al. 2021b), which generated DA neurons from hPSCs by boosting WNT signaling during early midbrain patterning. After the initial patterning for 11 days, cells were differentiated over 3 weeks. The resulting differentiated mDA cells robustly expressed the key differentiation markers *FOXA2* and *MAP2*, across subclones of familial PD mutant-expressing cells (Figure 1B, (Busquets et al. 2026)). Gene expression analysis from bulk RNA seq data of the differentiated mDA cells at day 35 confirmed the expression of key neuronal markers (e.g. *FOXA2* and *MAP2*) compared to undifferentiated WIBR3 hPSC cells (EWT control cell lines), indicating robust differentiation within these hPSC-derived cell lines expressing familial PD mutations (Figure 1B). Other single-cell RNA-seq data indicate that these cultures generated ∼15-60% mature DA neurons (Syed et al. 2026). The deep sequencing that was performed allowed the simultaneous assessment of global gene expression and alternative pre-mRNA splicing patterns since splicing pattern detection requires Illumina reads that cross splice junctions (Supplemental Figures S1-S12; Supplemental Tables 2-5). Notably, we observed minimal overlap between genes with altered expression levels and those exhibiting differential splicing (Supplemental Figure S14; Supplemental Table 8). We focused our subsequent analyses of these data on a curated set of genes known to be expressed in both our hPSC-derived midbrain lineage cells and in normal patient human postmortem substantia nigra dopaminergic neurons (Wang et al. 2024).

**Figure 1.**
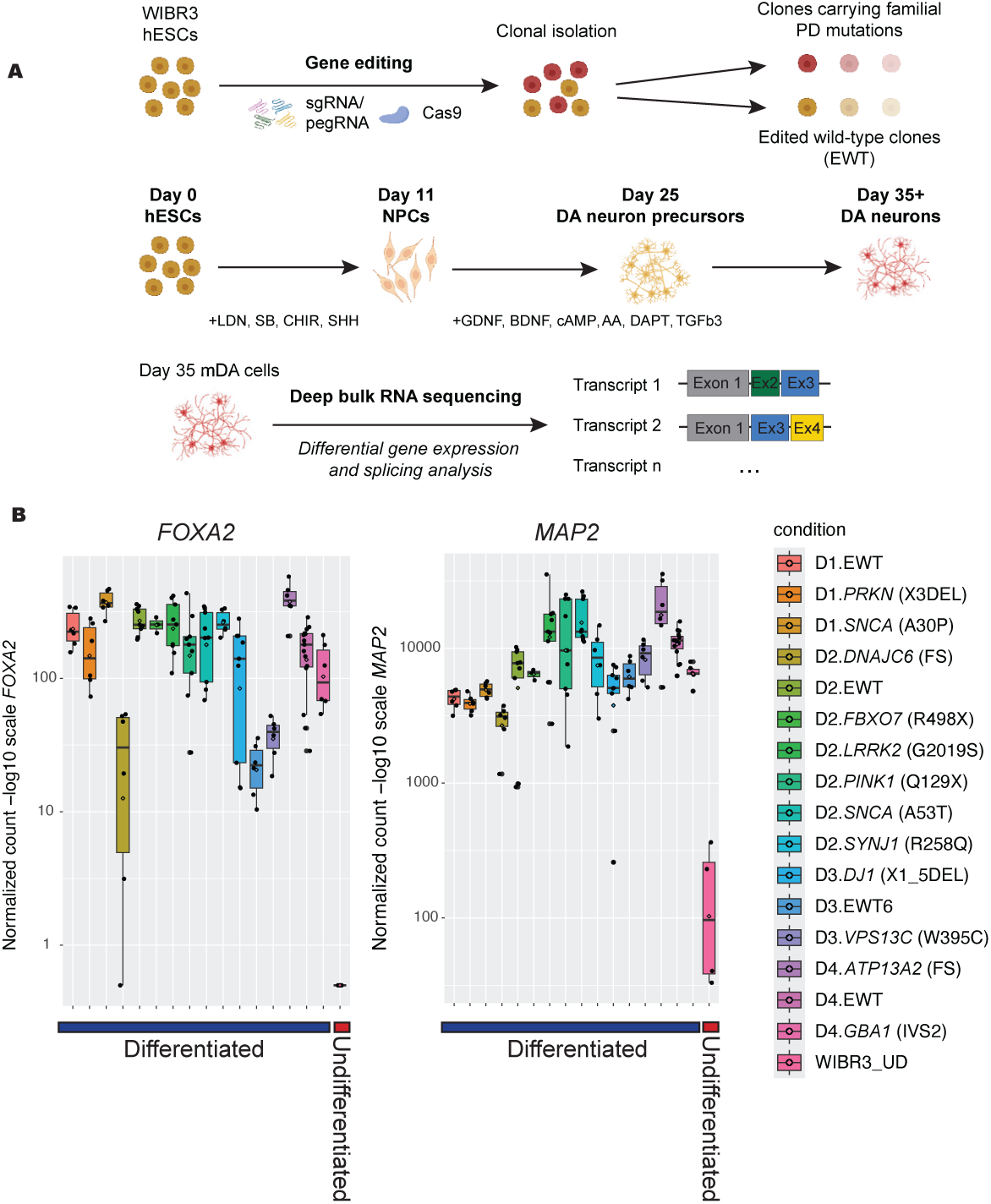
Differentiation of the genome-edited of human embryonic stem cells (hPSCs) carrying relevant familial Parkinson’s mutations into cells to midbrain lineage cells. (A) Schematic diagram of editing pipeline and expansion into clonal hPSC lines. (B) Normalized gene counts for neuronal markers including *FOXA2* (forkhead box A2) and *MAP2* (microtubule-associated protein 2), in 35d differentiated edited wild-type (EWT), familial *PRKN, SNCA, LRRK2, PINK1, DNAJC6, FBX7, SYNJ1, DJ1, VPS13C, ATP13A2* and *GBA1* mutant cells, in comparison to undifferentiated WIBR3 hPSCs. D1–D4 denote four independent sets of cell differentiations from bulk RNA sequencing datasets.

#### Identification of unique splicing signatures in hPSC-derived dopaminergic neurons carrying PD-associated mutations

During neuronal differentiation, alternative splicing can modulate diverse processes such as signaling activity, centriolar dynamics and metabolic pathways (Su et al. 2018). RNA-seq data can provide insights into the regulation of alternative splicing by revealing the usage of annotated or unknown (unannotated) splice sites as well as the expression level of exons. Using 35 day-differentiated mDA cells carrying common familial PD mutations or wild-type genotypes, we prepared total RNA, poly(A)-mRNA selected and then generated random-primed cDNA libraries from 1-3 independently derived subclonal cell lines in technical triplicates. These libraries were sequenced on the Illumina NovaSeq S1 or S4 platform with between 200-300M paired-end 150bp (PE150) reads. The large datasets coming from these experiments were analyzed using the JUM software (Wang and Rio 2018) to compare and quantitate pre-mRNA splicing pattern differences between the mutant and edited wild type (EWT) control cells. The ratio between reads including or excluding exons, also known as the percent-spliced-in index (PSI), indicates how efficiently sequences of interest are spliced into mature transcripts. We observed thousands of splicing events that differed between the EWT and mutant cell lines with the changes in PSI (ΔPSI) cut-off set to > 5%, in canonical splicing patterns and in a more complex category that JUM denotes as “composite”, which is a combination of splicing patterns beyond a simple splicing event within the same splicing structure (Wang and Rio 2018). RNA-seq data derived from 12 different familial mutant cell lines were analyzed for RNA splicing and gene expression changes compared to EWT controls (Supplemental Tables 2-4).

#### Common splicing pattern changes in DA-expressed genes linked to trafficking, neuronal structure and development and regulation of plasma membrane-bounded cell projection organization in the SNCA (A30P/A53T) and LRRK2 (G2019S) mutants

Alpha-synuclein (SNCA) is a prominent component in Lewy bodies and aggregation of this protein is commonly used as a marker for PD (Kruger et al. 1998). There are multiple mutations and genetic variations in the alpha-synuclein (*SNCA*) gene that drive PD pathology. While the A53T mutation is a well-known example, others include the previously mentioned A30P mutation and the E46K mutation, as well as gene duplications and triplications (Polymeropoulos et al. 1997; Kruger et al. 1998; Singleton et al. 2003; Chartier-Harlin et al. 2004; Zarranz et al. 2004). The mutations in *LRRK2* that are associated with aberrant kinase activity are common and important inherited mutations linked to PD, and a series of kinase inhibitor small molecules have been identified (Lesage and Brice 2009; Taymans and Greggio 2016). The G2019S mutation leads to a hyperactive form of the kinase (Jaleel et al. 2007; West et al. 2007; Greggio and Cookson 2009).

The *SNCA* and *LRRK2* variants analyzed in this study represent autosomal-dominant mutations. We analyzed and compared the JUM splicing pattern changes between the *SNCA* (A30P), *SNCA* (A53T) and *LRRK2* (G2019S) mutant cells and transcript isoform changes looking at the 2,250 mDA neuronally expressed genes (Wang et al. 2024); Supplemental Table 1). We evaluated the overlap of these DA neuron-expressed DS genes across the three mutant lines (Figure 2A; Supplemental Table 2). Genes showing differential splicing in at least two distinct dominant mutant backgrounds were defined as a shared, high-confidence DS signature and subjected to Gene Ontology (GO) enrichment analysis (Figure 2B). This analysis revealed significant enrichment in pathways critical for neuronal structure and development, including regulation of plasma membrane-bounded cell projection organization, neuron projection morphogenesis, and cytoskeleton organization (Figure 2B). Additional enriched terms highlighted broader neurodevelopmental dysregulation, such as regulation of neuron differentiation and regulation of neurogenesis (Figure 2B). An example of the JUM output showing quantitation of differential splicing patterns is shown (Figure 2C).

**Figure 2.**
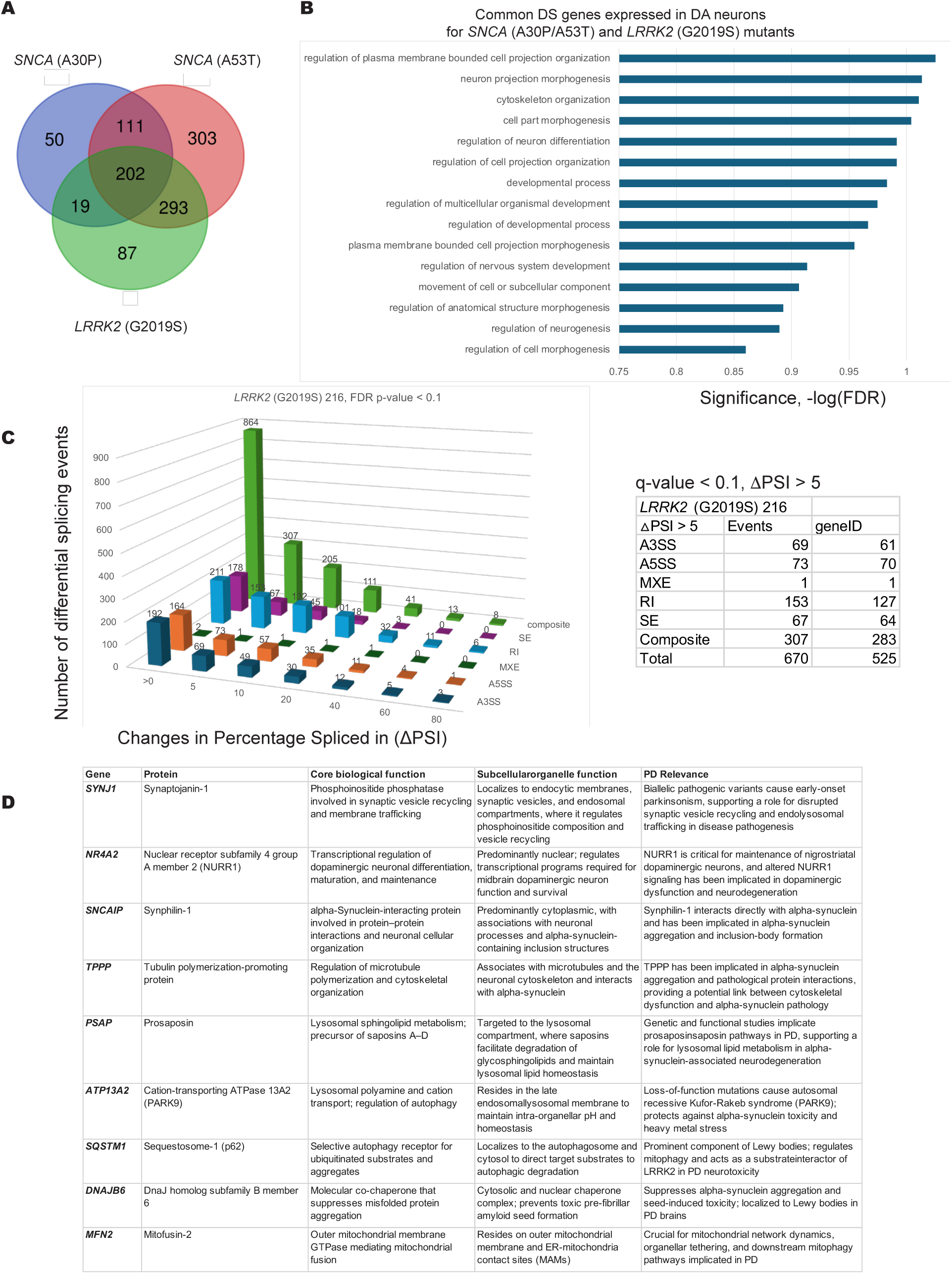
Identification and functional profiling of common differentially spliced (DS) genes in *SNCA*- and *LRRK2*-mutant dopaminergic (DA) neurons expressed genes. **(A)** Venn diagram of DS genes: Overlap of differentially spliced, DA neuron–expressed genes identified across *SNCA* (A30P), *SNCA* (A53T), and *LRRK2* (G2019S) iPSC-derived mutant models. The central intersections highlight 202 genes commonly altered across all three lines and 625 genes shared by at least two mutant lines. (B) Gene Ontology (GO) enrichment: Top biological process terms enriched among the shared DS genes expressed in DA neurons, ranked by significance [-log(FDR)]. Enriched pathways prominently feature neuron projection morphogenesis, plasma membrane organization, and cytoskeletal remodeling. (C) Distribution and types of splicing events: Representative distribution of splicing events in the *LRRK2* (G2019S) clonal line 216 across changes in percent spliced in (Delta PSI > 5). The accompanying table breaks down total events (n = 670, Delta PSI > 5) and underlying gene IDs (n = 525) by specific splicing type: alternative 3’ splice site (A3SS), alternative 5’ splice site (A5SS), mutually exclusive exons (MXE), retained introns (RI), skipped exons (SE), and composite events. (D) Prioritized PD-causing and risk genes: Table highlighting key shared DS genes with established roles in Parkinson’s disease pathology, detailing their official protein names, core biological functions, subcellular/organellar localizations, and disease relevance.

Notably, splicing of transcripts from the *(SYNJ-1)* gene are altered in the *SNCA* and *LRRK2* mutant cells, along with other genes linked to PD among the curated DA neuron-expressed genes (Figure 2D). *SYNJ-1* encodes synaptojanin-1 which is a lipid phosphatase involved in synaptic vesicle endocytosis (De Camilli 2026) and dopamine transporter regulation and release (Saenz et al. 2024). The *SYNJ-1* gene is itself a familial PD risk gene, and we find that its splicing pattern is altered in most of the other familial mutants iSCORE-PD cell lines we analyzed (Supplemental Table 2; see Figures 2-7).

While filtering for genes expressed in primary human SN neurons highlights cell-type-specific targets, examining the total pool of differentially spliced (DS) genes across all expressed transcripts from the bulk RNA-sequencing data provides a more comprehensive view of widespread pathway dysregulation of broadly expressed genes in these mutant lines. We also observed overlap of splicing changes from the bulk *SNCA* (A30P), *SNCA* (A53T) and *LRRK2* (G2019S) RNA-sequencing data (Supplemental Figures S1-S3) with a complete detailed analysis of these data (Supplemental Table 4). Interestingly, many of the genes in this GO term cluster whose splicing patterns are changed are linked to trafficking, mitochondria and lysosomes and include genes linked to PD (Bonet-Ponce and Cookson 2019; Bellucci et al. 2022; Komori and Kuwahara 2023; Pfeffer 2023; Tian et al. 2024);Supplemental Figures S1-S3). Finally, there is overlap in splicing patterns from the mutant hPSCs and transcript isoforms altered in patient brain samples from three Lewy body diseases (Feleke et al. 2021) (Supplemental Figures S1D-S3D). Alterations in the splicing isoforms produced from these genes in the PD mutant cells could affect membrane trafficking, organelle transport or mitochondrial/lysosomal function by changing protein levels or protein-protein interactions (Gonatopoulos-Pournatzis and Blencowe 2020).

#### Shared Splicing Alterations in DA Genes are Linked to Axon Guidance and Clathrin-Mediated Vesicle Trafficking in PRKN, PINK1, and DJ1 Mutants

Moving beyond dominant risk factors, a second distinct group of familial PD risk genes is linked to mitochondrial quality control and mitophagy via an autosomal-recessive inheritance pattern. PRKN is an E3 ubiquitin ligase that, together with the protein kinase PINK1, coordinates mitochondrial quality control through selective autophagy (mitophagy) and the degradation of damaged mitochondria (Narendra et al. 2008; Winklhofer 2014). Specifically, PINK1 acts as a mitochondrial damage sensor that recruits and activates PRKN to flag compromised organelles for clearance. The *PINK1* Q129X1 mutation is genetically linked to PD (Valente et al. 2004; Beilina et al. 2005; Ibanez et al. 2006; Ishihara-Paul et al. 2008). DJ1 is critical for mitochondrial function and mitophagy (Canet-Aviles et al. 2004; Dodson and Guo 2007; Thomas et al. 2011). Mutations in PARK7 (encoding the oxidative stress-protective protein DJ-1) are linked to early-onset Parkinson’s disease(Bonifati et al. 2003a; Bonifati et al. 2003b; Pankratz et al. 2006). Loss of DJ-1 function exacerbates oxidative stress and impairs mitochondrial ATP production via the accumulation of dysfunctional organelles, contributing to neurodegeneration.

We analyzed and compared the JUM splicing pattern changes among the curated DA neuron-expressed genes in the *PRKN* gene exon 3 deletion (X3DEL), *PINK1* (Q129X) and *DJ1* (X1-5DEL) mutant cells (Figure 3A and Supplemental Table 2). There is overlap in splicing isoform changes between the *PRKN* gene exon 3 deletion (X3DEL), *PINK1* (Q129X) and *DJ1* (X1-5DEL) mutant cells (Figure 3A). This analysis identified genes with GO terms for axon guidance, clathrin coated vesicle assembly, and receptor localization to synapse (Figure 3B). These genes are expressed in human postmortem SN DA neurons (Supplemental Figure S1; (Wang et al. 2024) and are in common between the three mutant cell lines (Figure 3B). An example of the JUM output data is displayed showing the type and quantitation of altered splicing events in the *DJ1 (X1-5DEL)* clonal mutant line 2860 (Figure 3C). Many of the genes whose splicing patterns are changed are linked to vesicle trafficking, lysosomes and the cytoskeleton and are genes linked to PD (Figure 3D). Notably, again we identified splicing pattern changes in the *SYNJ-1* pre-mRNA (Figure 3D), making a previously unidentified link between several of the familial PD mutants. We also observed overlap in splicing pattern changes from the bulk RNA-sequencing data from the *PRKN* gene exon 3 deletion (X3DEL), *PINK1* (Q129X) and *DJ1* (X1-5DEL) mutant cells (Supplemental Figures S4-S6; Supplemental Table 4). There is also overlap in splicing patterns from the hPSC-derived cells and transcript isoforms altered in patient brain samples from three Lewy body diseases (Feleke et al. 2021); Supplemental Figures S4D-S6D). Ultimately, these splicing alterations suggest a mechanistic link connecting distinct familial PD mutations to downstream disruptions in membrane dynamics, vesicular transport and endolysosomal function. These changes in splicing isoforms were likely missed by analyzing gene expression patterns alone.

**Figure 3.**
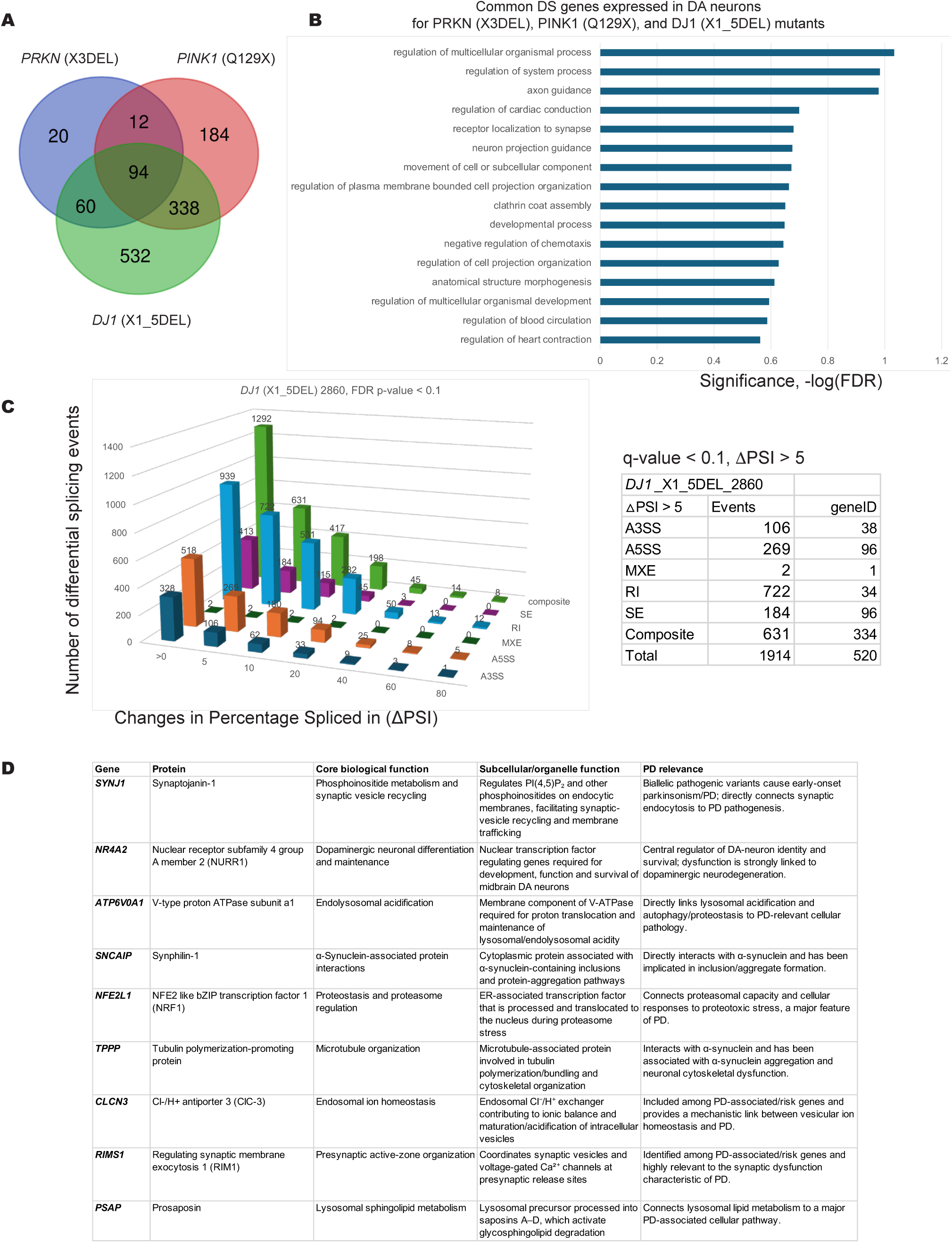
Mutations in *PRKN, PINK1,* and *DJ1* differentially affect gene splicing in autosomal recessive Parkinson’s disease, highlighting aberrations in axon guidance and synaptic vesicle dynamics. **(A)** Intersection and distribution of 504 high-confidence differentially spliced (DS) genes detected via JUM across *PRKN* (X3DEL), *PINK1* (Q129X), and *DJ1* (X1_5DEL) autosomal recessive mutant lines. Selection criteria required gene expression in DA neurons and differentially spliced in at least two recessive mutant conditions. (B) Pathway enrichment profiling of the 504 shared high-confidence DS targets, revealing preferential clustering in processes governing axon guidance, neuron projection navigation, clathrin coat assembly, and cell projection organization. (C) Quantification of absolute splicing changes (delta PSI > 5%, q-value < 0. 1) and event categories (A3SS, A5SS, MXE, RI, composite, and SE) are detailed in the bar chart and integrated table. The *DJ1* (X1_5DEL) clonal line 2860 is shown as a representative example to illustrate event compositions (including RI, SE, and composite events). (D) Biological overview of key Parkinson’s disease-associated genes identified within the 504-gene target pool, summarizing protein function, cellular localization, and relevance to recessive PD mechanisms.

#### Shared Alternative Splicing Changes in ATP13A2, DNAJC6, FBXO7, and SYNJ1 Mutants Converge on Lysosomal and Vesicle Trafficking Pathways

Expanding beyond core mitochondrial pathways, the third category of familial PD risk mutations encompasses genes linked to atypical or complex parkinsonism that operate primarily within endolysosomal dynamics and vesicular trafficking pathways. The *ATP13A2* (*PARK9*) gene encodes a lysosomal ion transport protein that functions in acidification of lysosomes. Mutations in the *ATP13A2* gene are linked to Kufor-Rakeb Syndrome, a form of early onset PD (Ramirez et al. 2006; Park et al. 2015; Fujii et al. 2023). Studies have reported decreased ATP13A2 protein levels in SN DA neurons and frontal cortex of PD patients compared to controls (Kitada et al. 1998; Dehay et al. 2012; Fujii et al. 2023). DNAJC6 is a co-chaperone protein implicated in vesicle transport and lipid homeostasis. It is found mutated in early onset PD patients (Edvardson et al. 2012; Olgiati et al. 2016). FBXO7 is a ubiquitin ligase that regulates the cell cycle and mediates mitochondrial clearance and mutations in this gene have been linked to PD (Di Fonzo et al. 2009; Zhou et al. 2018). SYNJ1 is a lipid phosphatase involved in endosomal trafficking and vesicle recycling and mutations in *SYNJ1* have been linked to PD, several neurological disorders (Drouet and Lesage 2014; Soukup et al. 2018; Choudhry et al. 2021); (Saenz et al. 2024; De Camilli 2026) and dysregulation of the dopamine transporter (Saenz et al. 2024).

The JUM splicing pattern analysis revealed overlap in splicing changes between the curated set of 2,250 DA neuron gene transcripts in the *ATP13A2* Frameshift (FS), *DNAJC6* (FS), *FBXO7* (R498X), *SYNJ1* (R258Q) mutant cells (Figure 4A, Supplemental Table 2). and in the bulk RNA-sequencing analysis (Supplemental Figures S7-S10). This analysis identified genes with GO terms for lysosomal and vesicle trafficking and GTPase signaling organization (Figure 4B) for the DA neuron genes that were expressed in human postmortem SN (Wang et al., 2024) (Supplemental Table 1) and that were in common between the four mutant cell lines (Figure 4A; Supplemental Table 2). Representative JUM-identified splicing events in the *SYNJ1* (R258Q) clonal mutant line A5, which affected 772 genes, are shown in Figure 4C. Many of the genes in this GO term cluster whose splicing patterns are changed are linked to vesicle trafficking and lysosomes (Figure 4D). Notably, splicing alterations were again observed in the transcripts from the *SYNJ1* gene (Figure 4D). The complete detailed analysis of these data from the bulk RNA sequencing experiments is shown in Supplemental Figures S7-S10 and Supplemental Table 3). Again, there is overlap in splicing patterns from the mutant hPSCs and transcript isoforms altered in patient brain samples from three Lewy body diseases (Feleke et al. 2021); Supplemental Figures S7-S10). Alterations in the splicing isoforms produced from these genes provide a genetic link between monogenic PD mutations and global dysregulation of endolysosomal and vesicular trafficking networks.

**Figure 4.**
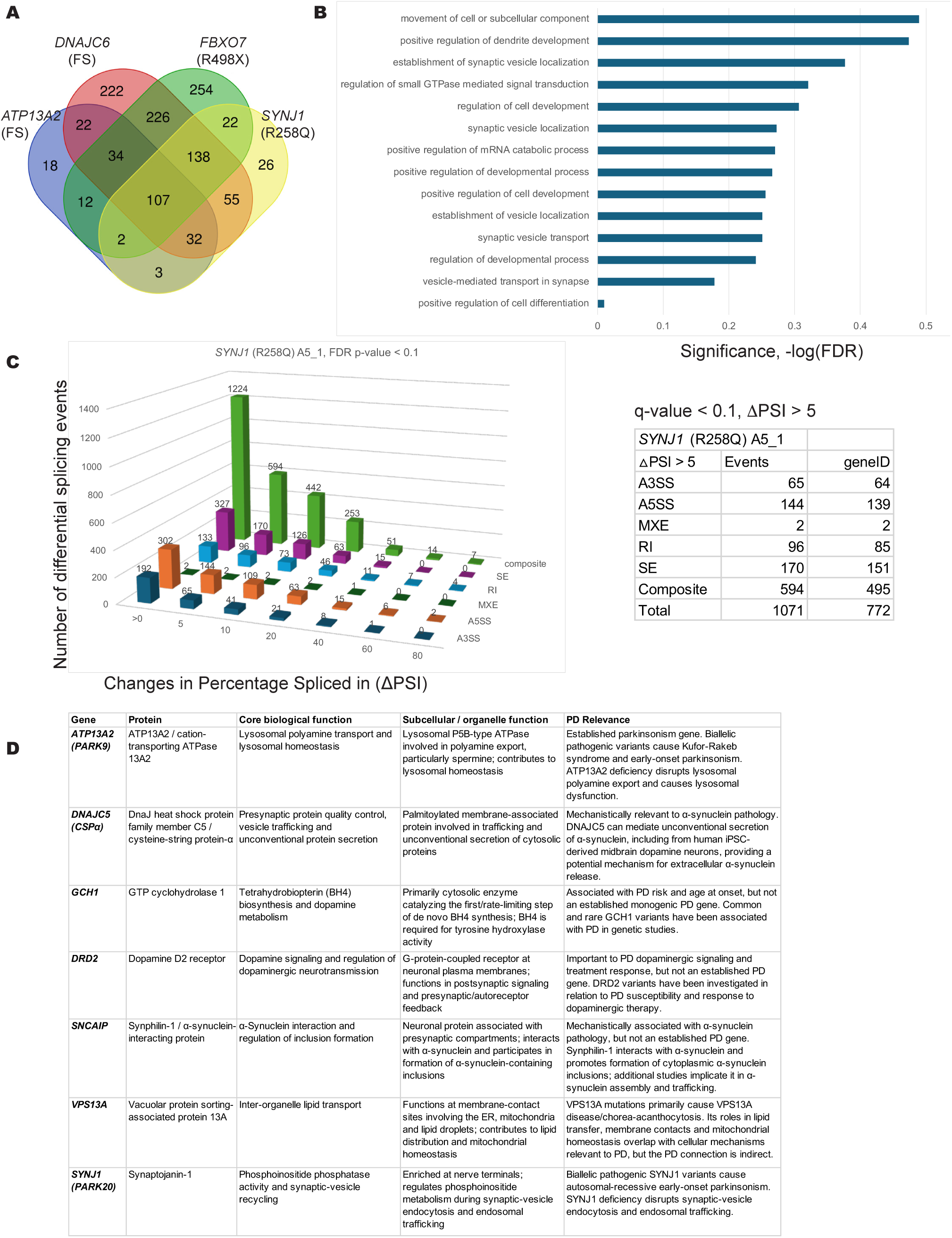
Differential pre-mRNA splicing profiling across atypical early-onset Parkinson’s disease mutant models highlight disruption of synaptic transport and vesicle localization. (A) Overlap of high-confidence differentially spliced (DS) genes expressed in DA neurons identified across four atypical early-onset parkinsonism models: *ATP13A2* (FS), *<u>FBXO7</u>* (R498X), *SYNJ1* (R258Q), and *DNAJC6* (FS). Filtering prioritized DA neuron-expressed transcripts shared between at least two mutant conditions. (B) Functional enrichment analysis of shared high-confidence DS genes, demonstrating significant enrichment for GO terms related to synaptic vesicle transport, vesicle localization, and positive regulation of cell differentiation. (C) Three-dimensional breakdown of alternative splicing event categories (A3SS, A5SS, MXE, RI, SE, and composite) plotted against magnitude of inclusion change (delta PSI). Data from the representative *SYNJ1* (R258Q) clonal line A5_1 illustrates event distributions under stringent significance filtering (q-value < 0.1, delta PSI > 5%). (D) Tabular summary of representative candidate genes impacted by alternative splicing within the dataset, highlighting protein identity, primary biological processes, subcellular localization, and relevance to Parkinson’s disease and related neurodegenerative disorders.

#### Aberrant Pre-mRNA Splicing in VPS13C (W395C) Mutants Impacts Lipid Metabolism and MAPK Signaling Pathways

The *VPS13C* gene encodes a massive lipid-transfer protein localized at organelle contact sites, playing a critical role in non-vesicular lipid transport between the endoplasmic reticulum (ER) and endolysosomal compartments (Kumar et al. 2018). Because of its unique structural function in maintaining organelle crosstalk and membrane lipid homeostasis, VPS13C is a distinct membrane-contact-site lipid-transfer factor, separate from classical endolysosomal factors such as GBA1 and from primary mitochondrial or synaptic machinery. Some studies suggest VPS13C plays a role in mitophagy (Lesage et al. 2016; Schrӧder et al. 2024; Wang et al. 2025). Recessive mutations in *VPS13C* gene are linked to early onset PD (Lesage et al. 2016; Schormair et al. 2018; Smolders et al. 2021). Loss-of-function mutations at splice sites in VPS13C cause pathogenic mis-splicing, leading to frame shifts or premature stop codons associated with early-onset Parkinson’s disease (Lesage et al. 2016). However, VPS13C is not typically recognized as a primary cryptic splicing target driven by TDP-43 pathology in neurodegenerative disease (Ma et al. 2022). Focusing on the 2,250 dopaminergic neuron– expressed genes (Supplemental Table 1) our splicing analysis identified 654 altered splicing events (Supplemental Table 2), with affected transcripts significantly enriched in the mitophagy and dopamine pathways (Figure 5A). Overall, using Gene Ontology, these genes seem linked to steroid metabolism and the mitogen-activated protein kinase (MAPK) cascade (Figure 5B). An example of the JUM output for the *VPS13C* (W395C) C3 1 cell clone is shown in Figure 5C. Strikingly, a substantial proportion of these splicing events involve intron retention, which could trigger nonsense mediated mRNA decay and lead to defects in proteostasis (Figure 5C). Cross-referencing these data with PD genetic databases revealed several PD-associated genes among the spliced transcripts expressed in DA neurons (Figure 5D). Specifically, alterations in splicing patterns were once again detected in transcripts derived from the *SYNJ1* gene itself (Figure 5D).

**Figure 5.**
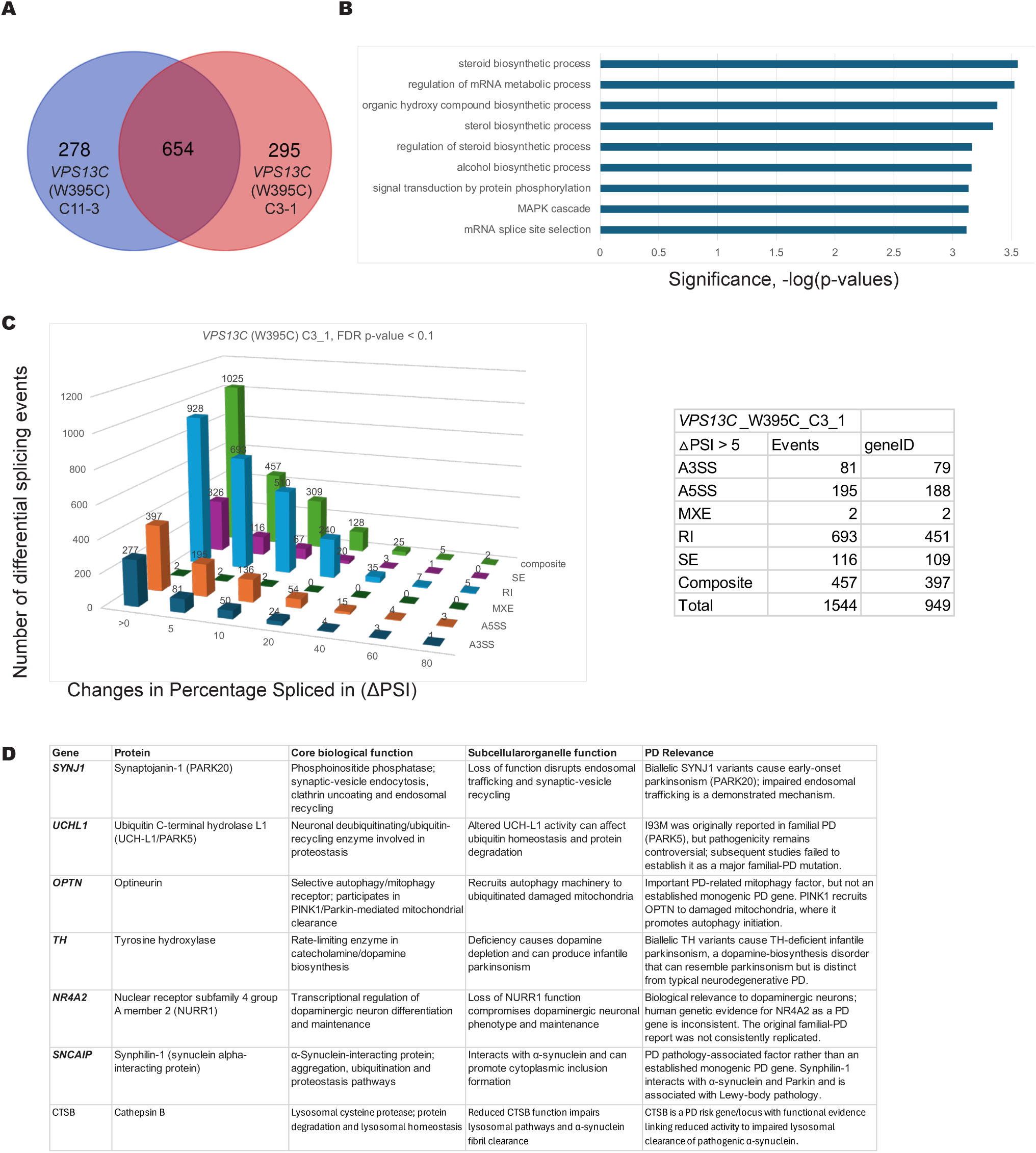
Pre-mRNA alternative splicing landscape in *VPS13C* mutant models reveals altered transcript processing and lipid/metabolic pathway enrichment. **(A)** Overlap of high-confidence differentially spliced (DS) genes identified among DA neuron-expressed genes between two independent clonal lines (C11-3 and C3-1) carrying the *VPS13C* (W395C) mutation, resulting in 654 high-confidence DS targets. (B) Gene Ontology (GO) pathway enrichment analysis performed on the 654 shared high-confidence DS genes. Significance scores (-log p-values) highlight primary enrichment in mRNA splice site selection, MAPK signal transduction cascades, and metabolic pathways, including sterol and steroid biosynthetic processes. (C) Three-dimensional profile of alternative splicing event categories (A3SS, A5SS, MXE, RI, SE, and composite) relative to changes in percentage spliced in (delta PSI) for the *VPS13C* (W395C) C3_1 line under stringent criteria (FDR p-value < 0.1, delta PSI > 5%). The accompanying table details total event counts and unique gene IDs per splicing class. (D) Functional matrix of key candidate genes identified within the *VPS13C* alternative splicing dataset, detailing primary protein functions, organellar/subcellular localizations, and specific pathological relevance to Parkinson’s disease and associated neurodegenerative traits.

Our bulk analysis of RNA from cells carrying the *VPS13C* W395C mutation identified 4,175 pre-mRNA splicing changes from two differentiated mutant cell clones compared to the EWT controls (Supplemental Figures S11A and S11B). The pre-mRNA isoforms that were altered in the *VPS13C* mutant cells are from genes linked to metabolic processes, chromatin modification and RNA splicing (Supplemental Figure S11C). Comparative analysis between splicing events affected in *VPS13C* mutant differentiated hPSCs and the publicly available RNA-seq data from Lewy body patient brains demonstrated that there was overlap between the transcripts whose splicing is affected, with Dementia with Lewy Bodies (DLB) having the most overlap (Feleke et al. 2021); Supplemental Figure S11D). The genes whose transcripts are affected at the level of pre-mRNA splicing in this data set are indicated in Supplemental Tables S2 and S3. Overall, these pre-mRNA splicing alterations highlight a crucial link between VPS13C dysfunction, impaired lipid homeostasis, and aberrant splicing changes prominently targeting genes in biosynthetic pathways (Figure 5B).

#### Disruption in GBA1 (IVS2) Mutant Alters Neurodevelopmental and Trafficking Networks

Due to its central role as the leading genetic risk factor for PD and its distinct enzymatic function in lipid catabolism, *GBA1* was analyzed as a standalone category (Gegg and Schapira 2018; Vieira and Schapira 2022). The *GBA1* gene encodes a lysosomal enzyme that maintains glycosphingolipid homeostasis (Sidransky et al. 2009; Huh et al. 2023). Mutations in the *GBA1* gene, such as the c.115+1G>A variant, are among the most common risk variants found in PD patients, as they can disrupt the gene’s pre-mRNA splicing pattern, leading to a non-functional glucocerebrosidase protein (Neumann et al. 2009; Onal et al. 2024). We compared the splicing pattern changes in two *GBA* mutant iSCORE-PD cell clones for the curated mDA neuron-expressed genes (Figure 6A and 6B). The 445 genes with altered splicing patterns include genes involved in cell projection organization, vesicle-mediated transport in synapses, and regulation of GTPase activity (Figure 6B; Supplemental Table 2). An example of the JUM data for one of the *GBA* mutant cell clones is shown in Figure 6C. Representative genes within this splicing dataset encode core proteins directly linked to PD pathogenesis and synaptic/lysosomal pathways, including *SNCA*, *SYNJ1*, *VPS13C*, *SQSTM1*, *ATP6V1A*, and *RIMS1* (Figure 6D). Importantly, splicing alterations in *SYNJ1* re-emerged within the *GBA1* mutant cells (Figure 6D), further corroborating pre-mRNA splicing dysregulation of this locus as a convergent molecular feature across familial PD models.

**Figure 6.**
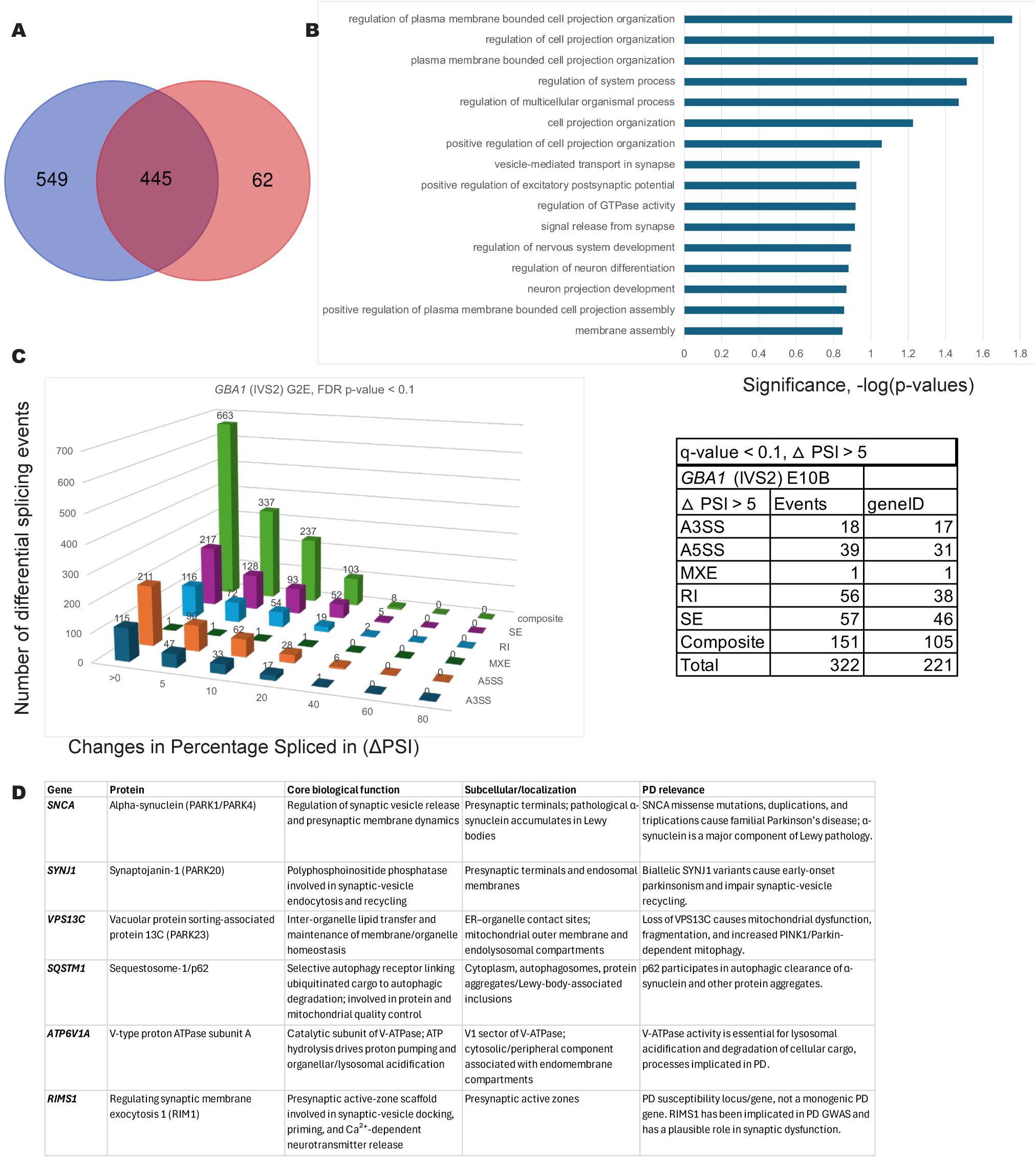
Alterations in pre-mRNA splicing profiles in *GBA1* (IVS2) mutant models implicate aberrant synaptic projection organization and endolysosomal transport pathways. **(A)** Distribution and overlap of 445 high-confidence differentially spliced (DS) genes identified among DA neuron-expressed transcripts across duplicate clonal lines of *GBA1* (IVS2) mutants E10B and G2E. (B) Gene Ontology (GO) biological process analysis of the shared 445 high-confidence DS genes. Significance scores (-log(FDR q-value)) highlight top enriched pathways governing cell projection organization, synaptic vesicle transport, excitatory postsynaptic potential regulation, and plasma membrane assembly. (C) Three-dimensional profile of alternative splicing event categories (A3SS, A5SS, MXE, RI, SE, and composite) plotted against inclusion level changes (delta PSI) for the GBA1 (IVS2) G2E line, shown as a representative example. The inset table quantifies total event counts and unique gene IDs under stringent filtering criteria (delta PSI > 5%, q-value < 0.1) for the G2B clonal line. (D) Functional profiling table of key Parkinson’s disease-relevant target genes identified within the *GBA1* dataset (including *SNCA, SYNJ1, VPS13C, SQSTM1, ATP6V1A*, and *RIMS1*), detailing protein identity, core biological function, organellar localization, and specific pathophysiological relevance to PD mechanisms.

Using mDA cells homozygous for the *GBA1* intron 2 mutation, a variant associated with Gaucher disease that also confers a significant risk for PD (Sidransky et al. 2009), analysis of the whole bulk RNA-seq data set identified 1,857 high-confidence pre-mRNA splicing alterations across two independent mutant clones (Supplemental Figures S12A and S12B). Genes whose splicing was affected include cell projection, cytoskeletal organization and regulation of GTPase activity (Supplemental Figure S12C). As observed with other mutants, there is overlap between the splicing event changes detected in the *GBA1* mutant cell line and the altered transcript isoforms from Lewy body disease brain biopsy RNA-sequencing data ((Feleke et al. 2021); Supplemental Figure S12D). The genes whose transcripts are affected at the level of pre-mRNA splicing are indicated in Supplemental Tables S2 and S3.

Differential alternative splicing (AS) events between EWT and *GBA1* (FS) mutants, initially identified by JUM analysis, were validated through RT–PCR assays. These experiments confirmed the predicted splicing shifts across multiple targets (Supplemental Figure S13). Expression profiling analysis indicated that genes involved in mitotic cell cycle checkpoint and microtubule-based processes are up-regulated, while genes involved in regulation of trans-synaptic signaling and regulation of transport are down-regulated in the *GBA1* IVS2 mutant cells compared to the EWT controls (Supplemental Figures S13E and S13F). The differentially expressed genes identified through GO enrichment analysis are detailed in Supplemental Table 5.

#### Splicing Pattern Changes in the SYNJ-1 Lipid Phosphatase and the Ubiquitin Ligase Ubr4 Are Founds in Multiple PD Familial Mutant hPSC Cell Lines and in PD Patients

Having observed that *SYNJ1* pre-mRNA is repeatedly alternatively spliced across multiple categories of familial PD mutations, we next sought to determine whether additional transcripts are broadly and recurrently targeted by alternative splicing across the diverse genetic backgrounds of our platform. We screened the differential splicing profiles of the twelve PD mutant lines, comprising mutations in *PRKN, SNCA, LRRK2, PINK1, DNAJC6, FBXO7, SYNJ1, DJ1, VPS13C, ATP13A2* and *GBA1*, to identify commonalities. Interestingly, we found two genes relevant to PD whose splicing patterns were changed in most or all of the familial PD mutant cell lines. In the curated set of DA neuron expressed genes, changes in the splicing patterns of *SYNJ-1* gene transcripts were detected (see Figures 2D-6D and Supplemental Table 6 and 7). *SYNJ1* is a complex gene comprising 32 exons and multiple pre-mRNA isoforms (Figure 7A), encoding a 145 kDa brain-enriched species and a 170 kDa ubiquitously expressed protein (De Camilli 2026). The prevalent splicing pattern changes we detected in the PD familial mutant cell lines are shown, which involve complex composite transcript patterns or skipped exons (Figure 7A; Supplemental Table 7). Two examples of changes in 3’ splice site use among the composite class of splicing events are shown that should alter the coding potential of the transcripts made beyond the two known isoforms encoding the 190 and 145kD proteins (Figure 7B and 7C). We note that common splicing events, including skipped exons and more complex composite patterns, use the exact same splicing junctions in the *SYNJ-1* gene across multiple mutants (Supplemental Table 7). The genetics of *SYNJ-1* show that even heterozygous mutants show defects in vesicle transport (Saenz et al. 2024; De Camilli 2026) and dopamine release (Saenz et al. 2024).

**Figure 7.**
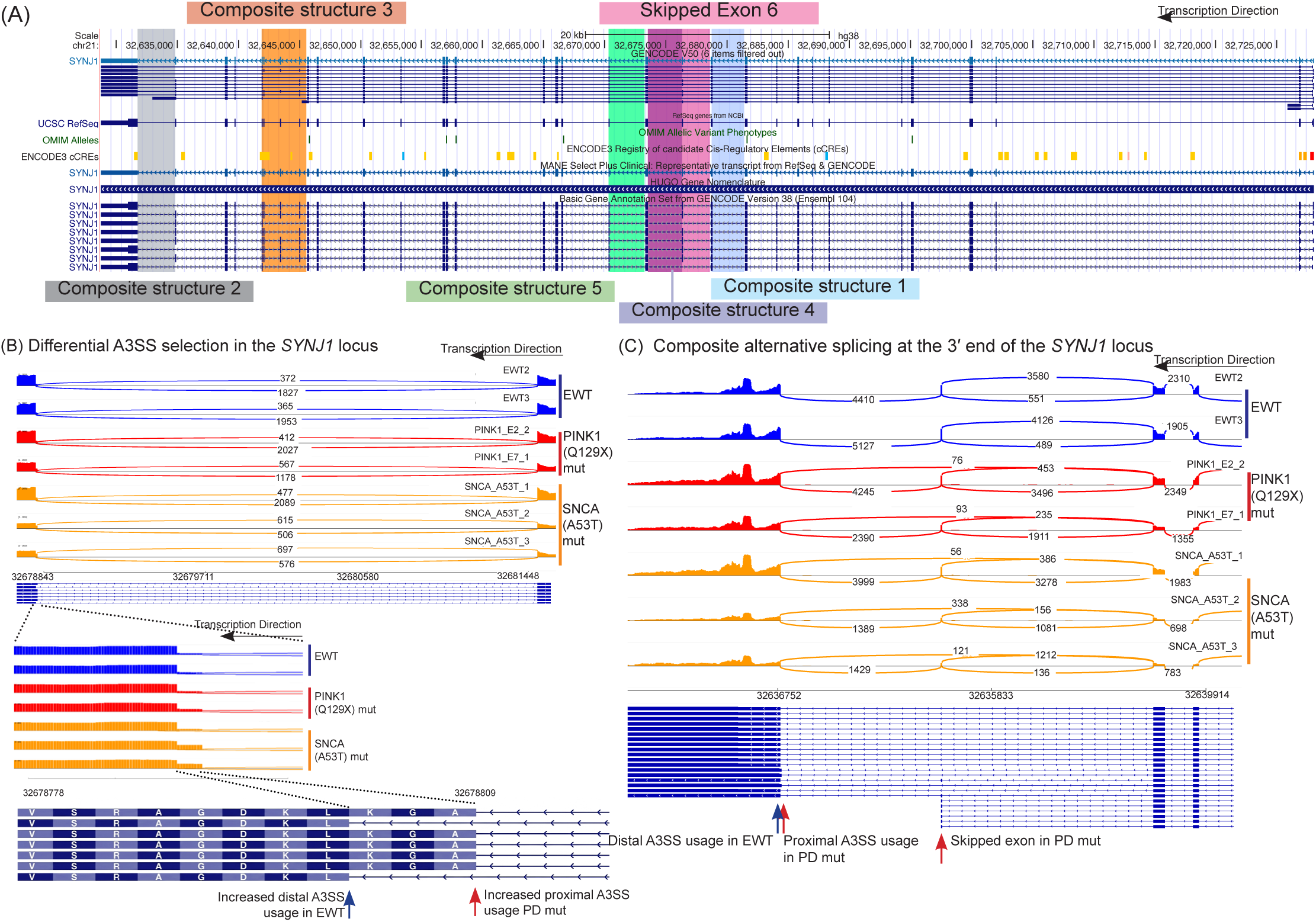
Alternative splicing and isoform diversity at the *SYNJ1* Locus in Parkinson’s disease models. (A) Genomic track view of the *SYNJ1* locus displaying composite transcript structures (1–5) alongside annotated annotations on hg38 UCSC browser. Green box highlights a skipped exon (6) region. Arrow indicates the direction of transcription. (B) Sashimi plots illustrating differential alternative splicing of Composite Structure 1 within the *SYNJ1* locus across wild-type (EWT) and Parkinson’s disease mutant lines [*PINK1* (Q129X) and *SNCA* (A53T)]. Splice junction reads quantify the preference for distal alternative 3’ splice site (A3SS) usage in EWT controls compared to increased proximal A3SS usage in the mutant lines. (C) Composite alternative splicing patterns near the 3’ end of the *SYNJ1* transcript across EWT and PD mutant replicates. Splice junction read densities highlight shifts between distal A3SS, proximal A3SS, and skipped exon events associated with the mutant genotypes.

Expanding our scope beyond DA neuron-enriched factors revealed that broad splicing dysregulation also targets ubiquitous cellular machinery; notably, *UBR4* (Ubiquitin Protein Ligase E3 Component N-Recognin 4) was differentially spliced in nine of the twelve mutant lines investigated (Supplemental Table 2 and 6). *UBR4* loss has been linked to mitochondrial dysfunction, excessive reactive oxygen species and impaired mitophagy (Jeong et al. 2025), which are defects linked to PD. UBR4 is a large ubiquitin ligase that functions as part of a protein complex linked to protein quality control and stress response signaling (Parsons et al. 2015; Kim et al. 2021a; Grabarczyk et al. 2025). The *UBR4* gene is large, having over 100 exons and whose transcripts are subject to alternative splicing. Our JUM analysis reveals that the wild-type transcript includes a microexon (exon 50 of 108) containing a premature termination codon. This microexon is predominantly included in control EWT cells, whereas its skipping (skipped exon event, SE) is significantly elevated in various PD mutant lines—such as the *LRRK2* (G2019S) mutant (Supplemental Figure S15A) and the complete differential splicing profile for the UBR4 gene (Supplemental Figure S15B). This would lead to more production of a protein-encoding transcript. We find that this particular exon (exon 50) is skipped more often in many of the familial mutant cell lines, as well as in some PD patient biopsies (Supplemental Figure S15; chr1: 19148626-19150575). These results indicate that splicing control would modulate the levels of active UBR4 protein and likely perturb the balance of UBR4 available to interact with its obligatory cofactors (Kim et al. 2018; Jeong et al. 2023; Grabarczyk et al. 2025) and thereby affect its ability to maintain proteostasis in PD patient neurons.

#### Differential Gene Expression Profiles of Day 35 hPSC-derived mDA Cells Carrying Familial PD Mutations

We used the RNA-sequencing data from the *PRKN, SNCA, LRRK2, PINK1, DNAJC6, FBXO7, SYNJ1,* DJ1*, VPS13C, ATP13A2 and GBA1* mutant and edited wild type control cell lines to analyze and compare overall gene expression (RNA transcript levels) using DESeq2 (Love et al. 2014). This analysis of the bulk RNA-seq data showed that there was differential gene expression of several hundred to several thousand genes in the *LRRK2*, *SNCA* (A53T), *PINK1*, DJ1*, VPS13C* and *SYNJ1* mutant cells (Supplemental Figures S1-S12D and F and Supplemental Table 5). Mutations in *SNCA (*A30P), *PRKN*, *FBXO7*, *ATP13A2*, *PINK1*, *DNAJC6*, DJ1, *VPS13C* and *SYNJ1* altered expression of genes linked to cell communication, synaptic transmission, membrane and ion transport, neuronal projections and protein trafficking to the membrane (Supplemental Figures S1-S12E and F; Supplemental Table 5). We note that there is some variability in the extent of differentiation of each mutant or wild type cell clone in a given experiment, but these measurements were made in technical triplicate with 1-3 independent cell clones per mutation with deep sequencing depth. We found that there was often variable overlap between gene expression versus pre-mRNA splicing changes in different mutant cells compared to wild type but that many of the splicing changes are distinct from the transcript level gene expression changes (Supplemental Figure S14; Supplemental Table 8). These observations may not be surprising, as alternative splicing in the nervous system frequently occurs without changing overall gene expression levels; instead, it generates diverse mRNA isoforms with altered junction structures, stability, or protein-coding sequences that modify downstream protein-protein interactions (Ellis et al. 2012; Gonatopoulos-Pournatzis and Blencowe 2020). The lack of changes in RNA levels of transcripts that undergo alternative splicing has been observed before (Ellis et al. 2012; Gonatopoulos-Pournatzis and Blencowe 2020; Feleke et al. 2021; Lee and Rio 2024). Alterations in transcript levels or isoform changes could lead to different levels or composition of protein complexes, altering proteostasis that would alter cellular phenotypes relevant to PD, including the cytoskeleton, GTPase regulation, organelle and vesicle trafficking, cilia, neuronal projections, as well as altered splicing factor networks (Supplemental Figures S1-S12; Supplemental Table 5).

Taken together, these experiments provide a comprehensive list of genes whose expression levels change in the iSCORE-PD cell collection (Busquets et al. 2026) carrying 12 distinct familial PD mutations when the hPSCs are differentiated to mDA neurons *in vitro*. Many of the genes whose expression levels are affected, like Rab family members (Bonet-Ponce and Cookson 2019; Bellucci et al. 2022; Alessi and Pfeffer 2024) and *SNCA* itself (Polymeropoulos et al. 1997; Dauer and Przedborski 2003; Singleton et al. 2003) are linked to PD. In other neurodegenerative diseases, RNA binding proteins, such as TARDBP/TDP-43 and FUS (Neumann et al. 2006; Hewitt et al. 2010; Mackenzie et al. 2010; Moens et al. 2025) are altered and which have been linked to ALS and frontotemporal lobar degeneration. In line with the canonical role of dysregulated RNA-binding proteins in ALS, our data suggest an emerging paradigm in Parkinson’s disease wherein broad pre-mRNA splicing dysregulation affecting critical synaptic and endocytic regulators, such as *SYNJ1,* acts as a unifying downstream effector across diverse familial PD etiologies (Supplemental Table 2 and 3).

## Discussion

Parkinson’s disease (PD) lacks a cure and L-DOPA, its main treatment, loses efficacy and causes side effects over time (Kwon et al. 2022; Dias et al. 2023). Early detection is challenging due to limited biomarkers (Concha-Marambio et al. 2023; Fernandes Gomes et al. 2023). Intensive fundamental research is vital to uncover underlying mechanisms for developing early diagnostics and new therapies (Schekman and Riley 2019). Defects in protein homeostasis, aggregation, RNA binding proteins and RNA processing lead to pathophysiological changes in PD, as is the case for other neurodegenerative diseases, like ALS and spinal muscular atrophy (SMA) (Brinegar and Cooper 2016; Conlon and Manley 2017; Mann and Donnelly 2021). Identifying RNA processing events and pre-mRNA splicing isoforms linked to these disease states has led to identification of disease-relevant transcripts for therapeutic interventions using antisense oligonucleotides (ASOs) (Bennett et al. 2019; Meyer et al. 2020), small molecules (Meyer et al. 2020), and gene therapy (Li et al. 2023). Given the advances in the delivery of nucleic acid therapeutics (Damase et al. 2021) and messenger RNA (Huang et al. 2022), it is likely that once key RNA level targets are identified for PD, therapeutic interventions can be developed. In fact, ASOs have been shown to downregulate mutant *LRRK2* G2019S mRNA in human macrophages to decrease immune responses (Wallings et al. 2023) and reduce LRRK2 function to mitigate mitophagy defects in mutant PD patient-derived cells and human LRRK2-expressing mice (Korecka et al. 2020). The platform developed and outlined here utilizing hPSCs with PD relevant mutations has the potential to aid in the development of PD therapeutics.

It is clear that for other diseases, such as a variety of cancers, including blood cancers like myelodysplastic syndrome (MDS), and neurodegenerative diseases like SMA, Huntington’s disease (HD) and ALS, that mutations in RNA binding proteins, known pre-mRNA splicing factors and/or specific splicing events can play causative roles in the disease states and are now successful therapeutic targets (Kapeli et al. 2017; Gebauer et al. 2021; Rogalska et al. 2023). Many neurodegenerative disease states involve changes in proteostasis, mitochondrial, lysosomal and other metabolic and cell biological pathways (Jagtap et al. 2023; Wilson et al. 2023). In many cases, protein aggregation, for instance the TDP-43 protein in ALS and α-synuclein in PD aggregates are key indicators of disease. In the case of ALS, two key RNA processing events involving the *stathmin 2* (*STMN2*) (Baughn et al. 2023) and *UNC13A* genes (Brown et al. 2022; Lipstein 2022; Ma et al. 2022) are targets for therapeutic interventions. Splicing alterations in *STMN-2* and *UNC13A* have recently been found in Alzheimer’s disease patients (Agra Almeida Quadros et al. 2024). In addition, there is widespread cryptic RNA splicing events leading to the production of novel peptides that might trigger immune responses (Akiyama et al. 2022; Stern et al. 2024) or neuronal dysfunction (Mehta et al. 2023; Guo et al. 2026). Recently, RNA-seq and proteomics data from the iSCORE-PD *LRRK2* (G2019S) mutant cell lines characterized here were used to identify cryptic RNA splicing patterns and expression of the encoded novel peptides were shown to overlap with those found in *TDP-43* mutant ALS samples (Brenton et al. 2025) highlighting the utility of this platform.

Here, we have developed a system to investigate RNA-level changes that are caused by mutations in 11 of the 20 most common familial PD hereditary genes, including *PRKN, SNCA, LRRK2, PINK1, DNAJC6, FBXO7, SYNJ1,* DJ1*, VPS13C, ATP13A2 and GBA1* (Poewe et al. 2017; Nalls et al. 2019; Blauwendraat et al. 2020). Our results linking splicing pattern alterations to defective PD relevant pathways expressed in DA neurons are consistent with defects in global pre-mRNA splicing patterns discovered in other neurodegenerative diseases such as ALS (Wang et al. 2020; Li et al. 2021), SMA (Singh and Singh 2018) and various cancers (Zhang and Manley 2013; El Marabti and Abdel-Wahab 2021; Stanley and Abdel-Wahab 2022; Bradley and Anczukow 2023; Choi et al. 2023). It is also clear that mutations in RNA binding protein genes or altered expression levels of RNA binding proteins, such as TDP-43, FUS and hnRNPA1 (Kapeli et al. 2017; Nussbacher et al. 2019; Lee and Rio 2024), splicing factors and spliceosome components, such as U2AF1 and SF3B1 (El Marabti and Abdel-Wahab 2021; Stanley and Abdel-Wahab 2022; Bradley and Anczukow 2023; Choi et al. 2023) play roles in neurodegeneration and cancer, respectively.

One important finding from this work is that there are many RNA splicing changes in the PD mutant hPSC-derived mDA cells for genes expressed human SN (Wang et al. 2024) and overlap with RNA splicing changes in genes identified in Lewy body disease biopsies (Feleke et al. 2021). The PD relevance of these data is supported by the splicing pattern changes in genes linked to defective organelle trafficking and function. In our studies with the familial *PRKN, SNCA, LRRK2, PINK1, DNAJC6, FBX7, SYNJ1,* DJ1*, VPS13C, ATP13A2 and GBA1* mutants, we have identified splicing pattern changes in most of the PD mutant cells for the *SYNJ-1* gene, itself a familial PD risk gene encoding a phospholipid phosphatase linked to vesicle recycling (De Camilli 2026) and dopamine release (Saenz et al. 2024). We also detected splicing changes for the well-known ubiquitin ligase *UBR4*, which is part of a ubiquitylation complex, linked to stress signaling pathways (Parsons et al. 2015; Kim et al. 2021a; Grabarczyk et al. 2025). Alterations in *UBR4* splicing can lead to different mRNA isoforms that could affect the function of UBR4 in proteostasis in mDA and PD patient neurons. Thus, the platform we have developed enabled the identification of common defects in RNA processing patterns in genes expressed in mDA cells derived from hPSCs engineered with a panel of PD familial mutations (Busquets et al. 2026). Moreover, this platform has been used to detect similar cryptic splicing patterns in *LRRK2* (G2019S) mutant cells with those of *TDP-43* splicing patterns in ALS (Brenton et al. 2025).

The complexity of splicing patterns in PD has also been noted for the *SNCA* gene where novel mRNA isoforms were detected that were previously unknown and may contribute to SNCA aggregation in PD (Evans et al. 2024). Thus, there is mounting evidence from both prior single-gene studies and genome-wide results that changes in pre-mRNA splicing patterns in PD patients may be critical for the pathogenies of PD (Bruno et al. 2026). Historically, RNA processing defects have been viewed primarily through the lens of ALS and frontotemporal dementia. However, our observations that common genes, such as *SYNJ*1, are differentially spliced across multiple familial PD mutations expressed in mDA neurons indicate that pre-mRNA splicing dysregulation may be widespread in PD. Mechanistically, this convergence across distinct genetic drivers (*LRRK2*, *PRKN*, etc.) may stem downstream from common cellular stressors in PD pathogenesis, such as chronic mitochondrial dysfunction, oxidative stress, or energy deficits, which are known to compromise the functional integrity of nuclear splicing factors and RNA-binding proteins. Alternatively, this may reflect a maladaptive transcriptomic signature unique to vulnerable midbrain dopaminergic neurons under stress. This highlights shared RNA-centric pathogenic mechanisms across distinct neurodegenerative disorders. Notably, beyond its canonical role in synaptic vesicle endocytosis, *SYNJ1* harbors an RNA Recognition Motif (RRM) domain (Saenz et al. 2024; De Camilli 2026) which raises the intriguing possibility that *SYNJ1* may moonlight as an RNA-interacting factor or directly interface with RNA processing machinery in dopaminergic neurons. Ultimately, while the functional consequences of these splicing alterations, and whether they directly drive pathophysiology, remain to be fully determined, this study uncovers a new aspect of RNA dysregulation in PD that can be targeted in future investigations to define its precise contribution to disease mechanisms.

## STAR Method

### KEY RESOURCES TABLE

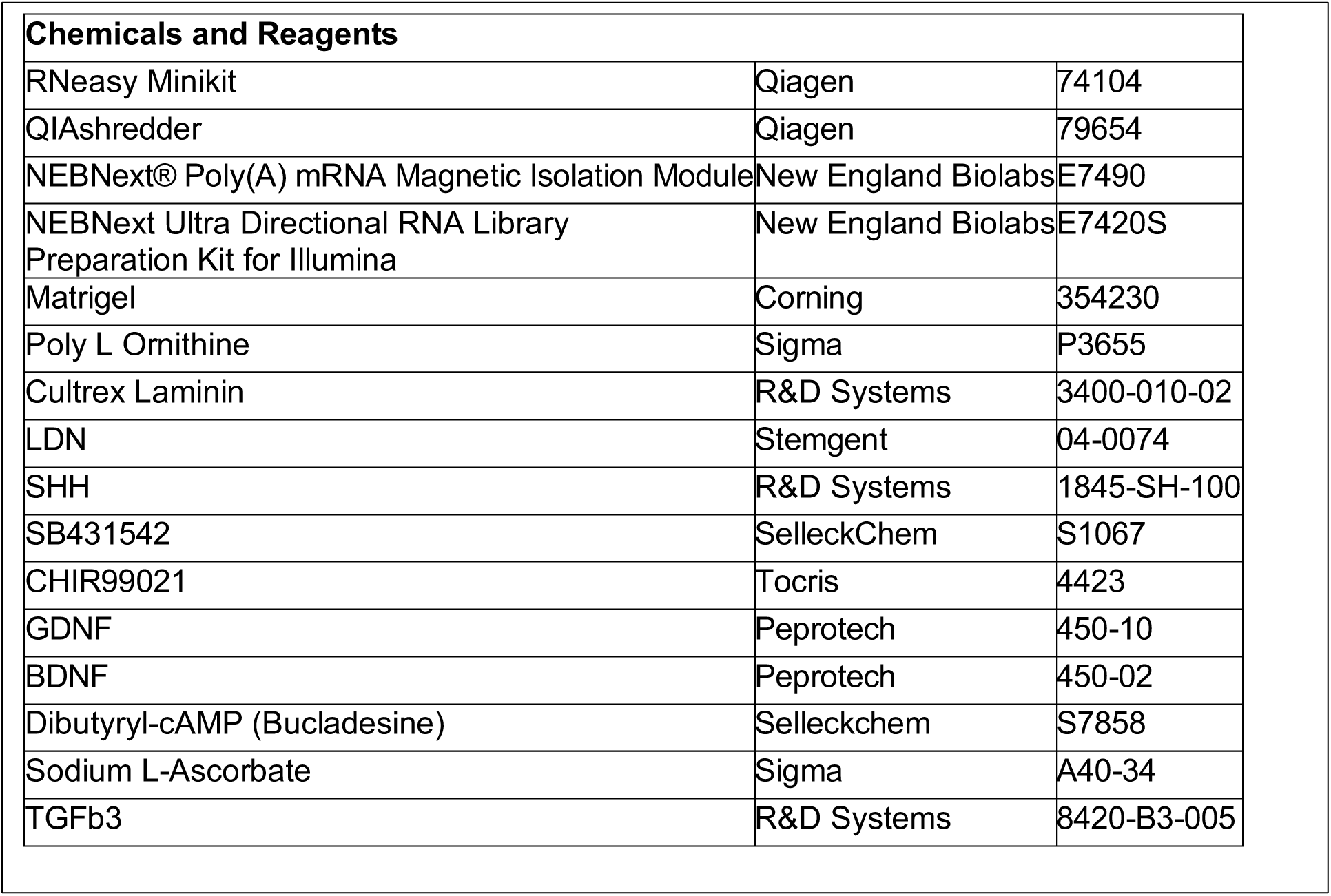

## Methods

### WIBR3-hPSCs derived DA neuron differentiation

WIBR3 hPSCs (passage 25-30) were maintained and grown on iMatrix-511 (Takara, T303) coated 6-well plates in mTESR plus media (Stem Cell Technologies, 100-0276). For DA neuron differentiation, the hPSCs were dissociated into single cells and seeded on matrigel (Corning, 356231) coated plates with Rock inhibitor (Chemdea CD0141) at 10 μM to initiate the differentiation. We followed the recently published protocol (Kim et al. 2021b) with slight modifications. Briefly, single cell suspension of hPSCs were maintained with Neurobasal N2/B27 (ThermoFisher, 21103049) media containing 2 mM glutamine (Sigma, G8450) with 200 ng/ml SHH C25II (R&D Systems, 1845-SH-100) 250 nM LDN (Stemgent, 04-0074),10 μM SB431542 (SelleckChem, S1067), 0.7 μM CHIR99021 (Tocris, 4423) on day 1 of differentiation. On day 4, CHIR concentration was increased to 7.5 μM and on day 7 LDN, SB, SHH were removed, and media was supplemented with only CHIR. On day 10, media was changed to Neurobasal B27 with 2 mM glutamine supplemented with 20 ng/ml BDNF (Peprotech, 450-02), 20 ng/ml GDNF (Peprotech, 450-10), 200 μM ascorbic acid (AA) (Sigma, A40-34) 500 μM dbcAMP (SelleckChem, S7858) and 1 ng/ml TGFb3 (R&D Systems, 8420-B3-005). On Day 11, cells were dissociated with Accutase (ThermoFisher, 00-4555-56) and split into a 1:2 or 1:4 ratio and replated on Poly L-Ornithine (PLO; 30 μg/ml; Sigma, P3655), Cultrex laminin (1 μg/ml; R&D Systems, 3400-010-02) coated plates and maintained in maturation media containing BDNF, GDNF, TGFb3, AA, DAPT from day 12. On day 16, cells were dissociated with Accutase (ThermoFisher, 00-4555-56) and replated at desired high density (1-2x10^6^ cells per well of 12 well plate) depending on cell growth and maintained in maturation medium (at this stage cells were cryopreserved in serum free cryopreservation media cell banker 2 (Amsbio, 11914) and thawed as per the experiment). On day 25, cells were dissociated with Accutase (ThermoFisher, 00-4555-56) and replated at higher density (>1x10^6^ cells per of 12 well plate until the experiment. On day 35, cells were lysed in the RLT lysis buffer using Qiagen kit (RNeasy, 74104) and RNA was isolated for bulk RNA sequencing.

### RT-PCR validation for splicing analysis

To validate several changes in alternative pre-mRNA splicing patterns found in the RNA-seq data we used reverse-transcription polymerase chain reaction (RT-PCR). In these experiments, we used analysis of *GAPDH* as a loading control. We found alterations in the levels of spliced mRNAs (Supplemental Figure S13). 0.5 microgram of total RNA was reverse-transcribed according to the manufacturer’s instructions (Bio-Rad, 1708891) and subjected to RT–PCR with the following conditions: 30 sec at 98°C (one cycle); 10 sec at 98°C, 30 sec at 60°C, and 30 sec at 72°C (35 cycles); and 5 min at 72°C (one cycle). The primer sets used in the RT-PCR reaction is listed in Supplement Figure S10B. RT-PCR products were resolved, visualized, and quantitated by use of an 2% agarose gels. Differential splicing events for *UBR4*, *DOCK3*, and *CAMPTA2* were shown as examples in *GBA1* (FS) mut compared to EWT control cells (Supplemental Figure S13).

### RNA-seq library preparation and sequencing

RNA isolation was performed using the RNeasy Minikit (Qiagen, 74104), followed by 30 min. of DNase treatment (Ambion, AM2238) at 37°C, poly(A)^+^ RNA transcripts were isolated [NEBNext poly(A) mRNA magnetic isolation module; New England Biolabs, E7490] from 0.5 µg of total RNA for RNA library preparation and sequencing using NEBNext Ultra Directional RNA Library Preparation Kit for Illumina (New England Biolabs, E7420S) according to the manufacturer’s instruction. The samples were sequenced on an Illumina S1 or S4 with 150-bp paired-end reads at the Vincent J. Coates Genomics Sequencing Laboratory at the University of California at Berkeley. Typical samples gave ∼200-300M PE150 reads.

### Analysis of pre-mRNA splicing patterns using JUM and differential expression analysis using DESeq2

Pre-mRNA splicing analysis using JUM.2.0.2 software to detect pre-mRNA splicing patterns was performed as described before (Lee et al. 2018; Wang and Rio 2018). An adjusted *p*-value (FDR) of 0.1 was used as the statistical cutoff for differentially spliced AS events with at least 10 junction reads.

For Differential expression analysis, uniquely mapped reads were mapped to hg38 using STAR_2.5.3a (Dobin et al. 2013) modified version using --outFilterMultimapNmax 1. Read counts were calculated using HTSeq script htseq-count (Anders et al. 2015) with the following specifications: -s yes -r pos -f. Differential gene expression analysis was performed using DESeq2 (Love et al. 2014), only genes with at least 10 reads across all samples were considered for further analysis.

## Resource Availability

### Lead contact

Further information and requests for resources and reagents should be directed to and will be fulfilled by the lead contact, Dr. Donald Rio.

### Data deposition

The data reported will be deposited in the Gene Expression Omnibus (GEO) database upon the acceptance of the manuscript.

Publicly available RNA-seq data for WIBR3 hPSCs were downloaded (GEO accession numbers: GSM3069316, GSM3069317, GSM3069324, and GSM3069325) for comparison with differentiated EWT and familial PD mutants.

## Acknowledgments

This work was funded by Aligning Science Across Parkinson’s (ASAP-024409, ASAP-000486) through the Michael J. Fox Foundation for Parkinson’s Research (MJFF) and the NIH (R35GM118121; DR). For the purpose of open access, the authors have applied a CC-BY public copyright license to the Author Accepted Manuscript version arising from this submission. Illumina RNA sequencing was carried out at the U.C., Berkeley QB3 Coates Genome Sequencing Laboratory (GSL), supported by National Institutes of Health (NIH) S10 Instrumentation grants S10RR025622, S10RR029668, and S10RR027303.

## Author Contributions

D.H., F.S., H.S.B and D.C.R. supervised the study; Y.J.L. performed RNA isolation and RNA-seq library preparation, quality controls, RT-PCR validation, the bioinformatic and statistical analyses of alternative splicing and gene expression profiling, generated all the figures and tables. K.M.S performed dopaminergic differentiations from genome-edited cell lines and contributed Figure 1A. H.L, O.B, J.D established the PD familial mutant cell lines. Y.J.L. and D.C.R. wrote the initial draft of the manuscript. All authors contributed to and edited the final manuscript.

