## Supplemental_Figures for "Familial Parkinson’s Disease Mutations Converge on Alternative Splicing Programs Linked to Lysosomal, Autophagic and Vesicle Trafficking Pathways"

#### Supplemental Figure S1

**A**

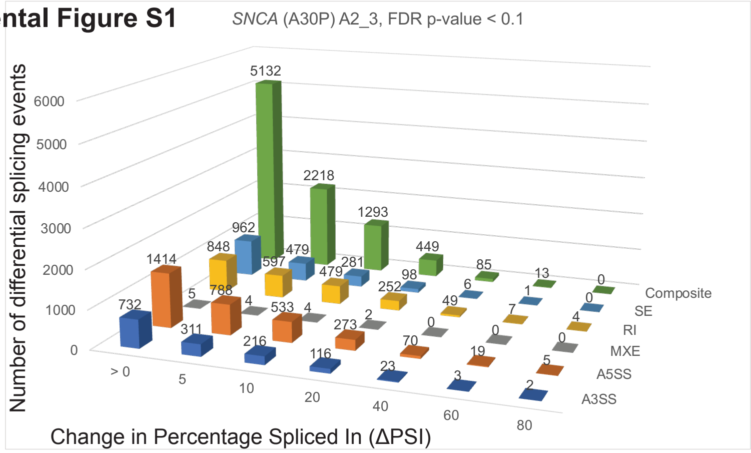

| FDR p-value < 0.1, ΔPSI ≥ 5 | | |
|---|---|---|
| *SNCA* (A30P) A2_3 | | |
| PSI >5 | Events | geneID |
| A3SS | 311 | 235 |
| A5SS | 788 | 651 |
| MXE | 4 | 4 |
| RI | 597 | 363 |
| SE | 479 | 428 |
| Composite | 2218 | 1826 |
| Total | 4397 | 3103 |

**B**

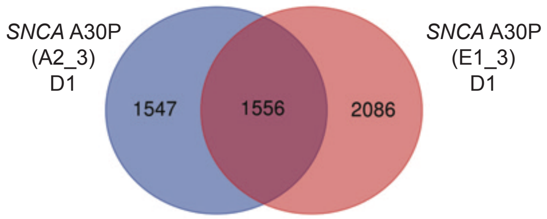

**C** GO-Enrichment analysis of high-confidence 1,556 DS genes in *SNCA* (A30P) mutant

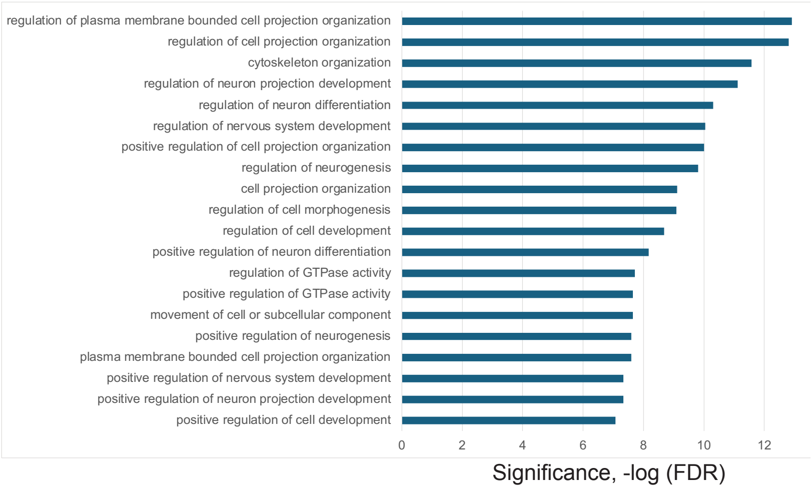

**D**

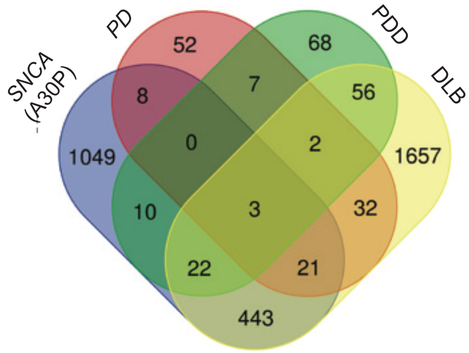

| | total | Genes |
|---|---|---|
| DLB, PD, PDD, *SNCA* (A30P) | 3 | *SEC14L1, SRRM2, EPB41L2* |
| DLB, PD, *SNCA* (A30P) | 21 | *FYN, MAST4, KIDINS220, PTPRF, EML4, WDR27, PPP1R21, CALD1, UBR4, EVL, NRXN2, L3MBTL1, MACF1, COL16A1, SHTN1, PRUNE2, AK2, ABLIM2, EML2, DOCK10, MICAL2* |
| DLB, PDD, *SNCA* (A30P) | 22 | *DLG1, NADSYN1, TARSL2, CLIP2, PPP1R32, LINC-PINT, TARBP1, MYO9A, SYTL2, ERBIN, COL16A1, SHTN1, ARHGAP17, STX2, ANK2, UNK, TRIM9, DYNC2H1, INO80C, ATP5SL, ROM1, ST18* |
| PD, *SNCA* (A30P) | 8 | *TENM4, FHL1, SYT7, TXNL4A, FMNL1, PSD3, ZNF708, RASGRF2* |
| PDD, *SNCA* (A30P) | 10 | *TMEM161B-AS1, LAMA3, SDHAF2, TMEM91, POLR2J3, INPP1, MAP7D2, ESD, ANK1, KIAA1191* |

**E**

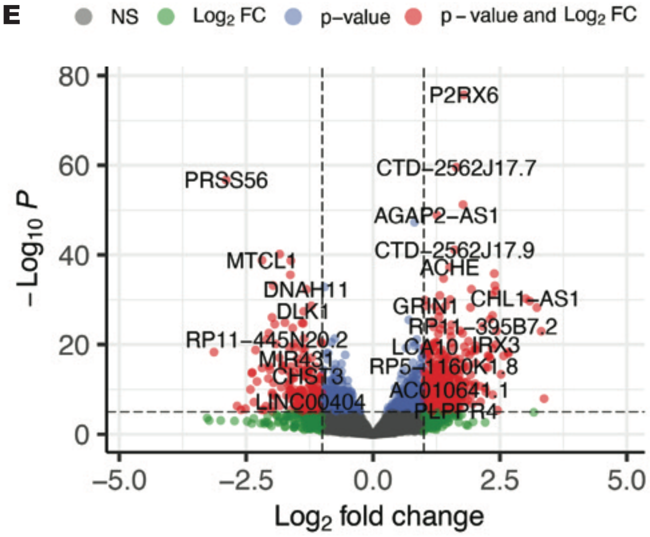

**F** GO-Enrichment analysis of 1,393 up-regulated DE genes in *SNCA* (A30P) mutant

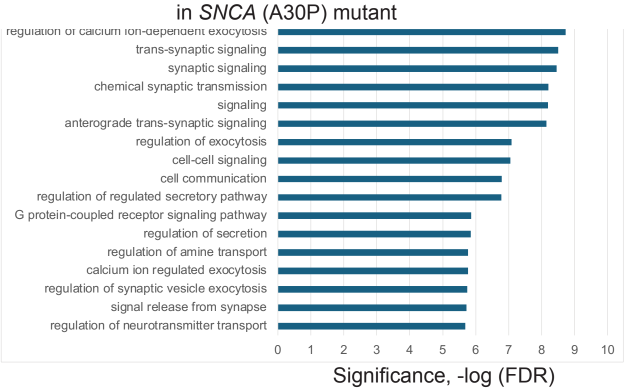

GO-Enrichment analysis of 944 down-regulated DE genes in *SNCA* (A30P) mutant

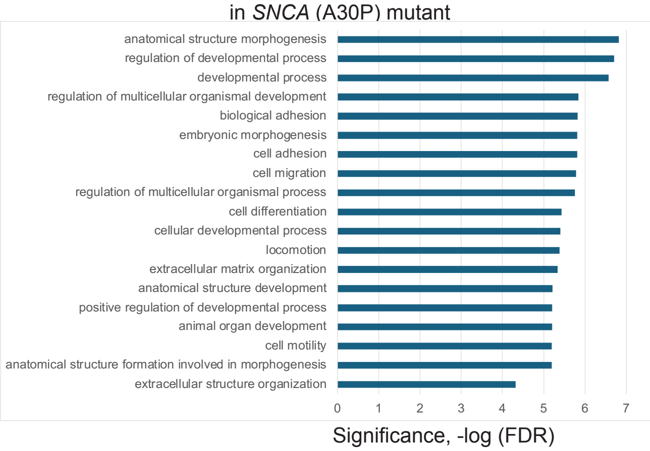

**Supplemental Figure S1. Differentially spliced and differentially expressed transcripts in mDA neuronal cells carrying the familial *SNCA* (A30P) mutation.** A. Differential alternative splicing events detected upon *SNCA* (A30P) mutation in differentiated DA neurons and progenitor cells (35d) using JUM (Wang and Rio, 2018). 1,566 high confidence splicing events (found in at least two mutant cell clones) were significantly altered in the *SNCA* (A30P) mutant samples in DA neurons and progenitor cells versus the edited wild type controls (EWT; with adjusted p-value < 0.1, ΔPSI ≥5). Data for one of three clonal mutant cell lines is shown as an example. Shown on the y-axis is the number of differential splicing events and differences in percentage-spliced-in (or ΔPSI) is shown on the x-axis. The splicing events were filtered based on the magnitude of delta PSI (ΔPSI). B. Comparison of genes that have differentially spliced transcripts in different clonal lines of the *SNCA* (A30P) mutant, this results in 1,556 high confidence differentially spliced transcripts. D1 indicates neuronal differentiation set number. C. A graph of Gene Ontology (GO) term enrichment of 1,556 differentially spliced transcripts in *SNCA* (A30P) mutant protein-expressing DA neuronal cells compared to the edited wild type control (EWT). These genes are involved in regulation of cell projection organization, regulation of neuron differentiation and regulation of GTPase activity. D. Comparison of differentially spliced transcripts in *SNCA* (A30P) mutant cells to differentially spliced transcripts from genes found in Lewy body disease patient brain biopsy RNA-seq data (Feleke et al., 2021): (Parkinson disease (PD), Parkinson disease with dementia (PDD), dementia with Lewy bodies (DLB)). E. A volcano plot of D1 (differentiation set 1) is shown as an example with some up- and down-regulated gene names shown. DESeq2 analysis of the *SNCA* (A30P) mutant compared to the edited wild type (EWT) controls reveals 2,338 differentially expressed genes (adjusted p-values < 0.05, FC > 1.5). F. A graph of Gene Ontology (GO) term enrichment of either 1,394 up-regulated genes or 944 down-regulated genes in the *SNCA* (A30P) mutant protein-expressing mDA neuronal cells compared to the edited wild type (EWT) controls.

**Supplemental Figure S2**

**A**

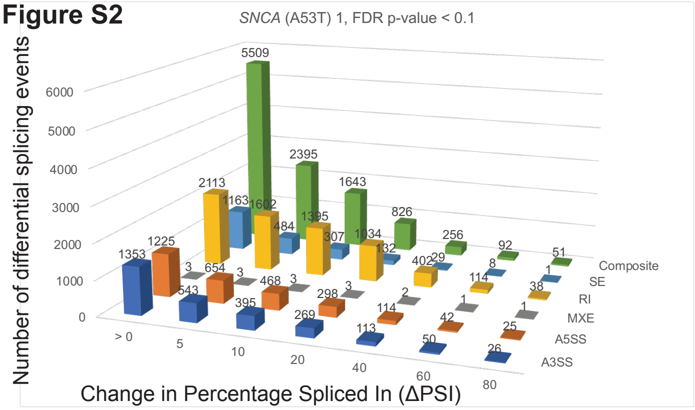

| FDR p-value < 0.1, △PSI ≥ 5 | | |
|---|---|---|
| *SNCA* (A53T) 1 | | |
| PSI >5 | Events | geneID |
| A3SS | 543 | 404 |
| A5SS | 654 | 539 |
| MXE | 3 | 1 |
| RI | 1602 | 826 |
| SE | 484 | 444 |
| Composite | 2395 | 2045 |
| Total | 5681 | 3714 |

**B**

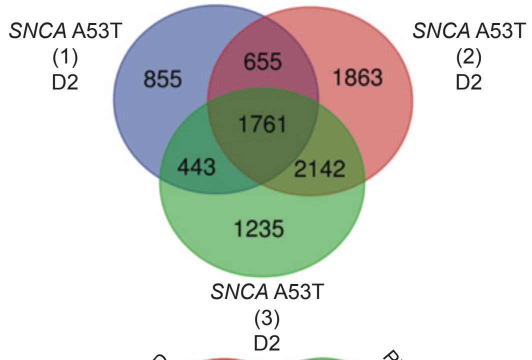

**C**

GO-Enrichment analysis of high-confidence DS genes in *SNCA* (A53T) mutant

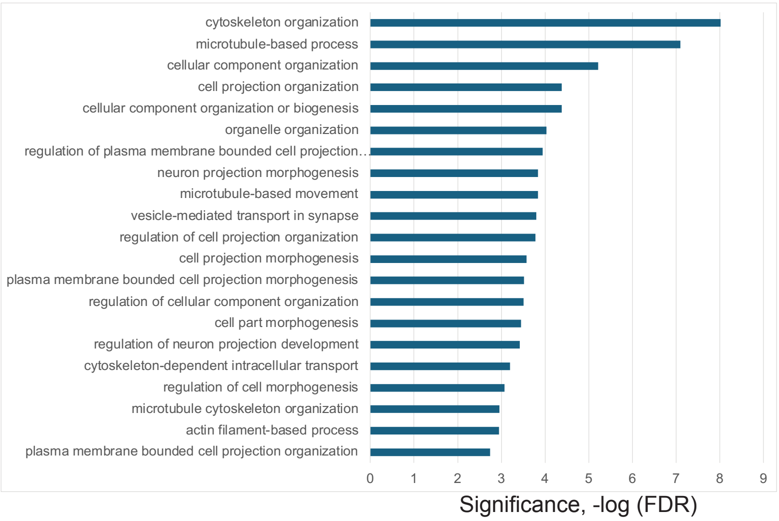

**D**

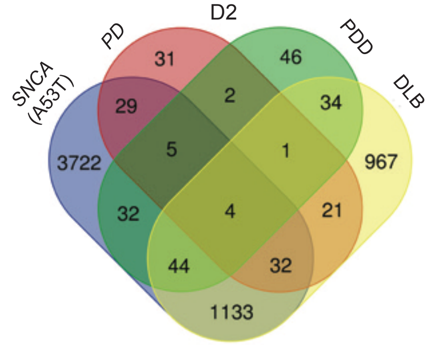

| | total | Genes |
|---|---|---|
| DLB, PD, PDD, *SNCA (A53T)* | 4 | *SEC14L1, SEC24C, SRRM2, EPB41L2* |
| PD, PDD, *SNCA* (A53T) | 5 | *PLXNB2, MEF2BNB-MEF2B, SULT1A1, DDX46, ZGPAT* |
| DLB, PD, *SNCA* (A53T) | 32 | *UPF3A, ABHD12, MAST4, EML4, WDR27, GRK4, KIF3A, ZNF789, CALD1, UBR4, NRXN2, MACF1, PRUNE2, ABLIM2, MRPL43, EML2, TCTN2, AP3B1, FYN, SPIDR, KIDINS220, PTPRF, EVL, L3MBTL1, LPGAT1, CADPS, CD46, AK2, APIP, DENND3, DOCK10, MICAL2* |
| DLB, PDD, *SNCA* (A53T) | 44 | *ZFHX4-AS1, ACAP3, CLIP2, LINC-PINT, STK39, MYO9A, USP33, ERBIN, SHTN1, ARHGAP17, RBMS2, STRADA, AP4B1, ANK2, UNK, DYNC2H1, ZDHHC11, ING4, MSI2, DLG1, FBXO42, BRD8, WIPI2, VEZT, PPP1R32, FAM228B, TARBP1, CTTNBP2, TNRC6A, IKBKB, NPIPA8, SYTL2, COL16A1, LUC7L3, MPRIP, STX2, PDE8A, TRIM9, INO80C, ODF2, OPTN, NT5C2, FAM49B, ST18* |
| PD, *SNCA* (A53T) | 29 | *LOC101927359, FHL1, LRRC16B, TCFL5, FMNL1, SLIT2, PSD3, ANKZF1, SYNC, VARS2, SLC12A5, ARHGAP6, RASGRF2, TENM4, BBS7, RBFOX1, WBSCR22, RYR1, SYT7, TXNL4A, TYK2, ZNF91, LINC00403, ETV1, CCZWPW1, MYO5A, RRP8, ZNF708, CYB561D1* |
| PDD, *SNCA* (A53T) | 32 | *ZNF83, GFPT2, TMEM161B-AS1, NOXA1, LAMA3, CHD2, PLEKHH1, PANK2, ST6GAL1, UBR5, INPP1, MAP7D2, ARHGAP18, PREB, OGFOD3, XPO1, ANK1, KIAA1191, UGP2, EOGT, DCAF8, UBE2F-SCLY, DRAM2, KIAA0586, CCDC113, DPY19L1P1, SDHAF2, TMEM91, ELP2, ESD, DROSHA, DOT1L* |

**E**

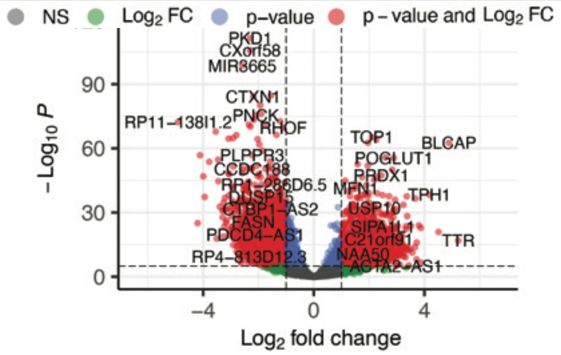

**F**

GO-Enrichment analysis of 4,708 up-regulated DE genes in *SNCA* (A53T) mutant

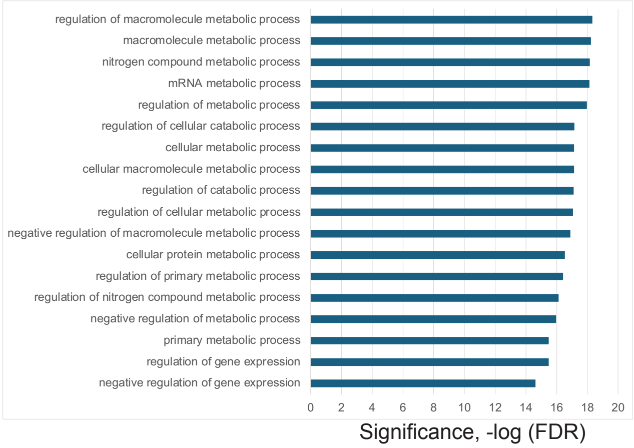

GO-Enrichment analysis of 4,759down-regulated DE genes in *SNCA* (A53T) mutant

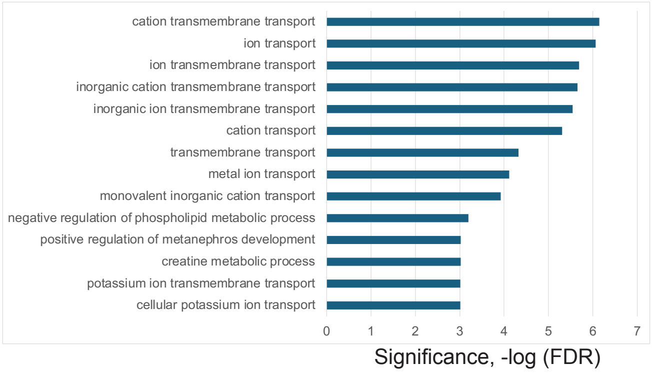

**Supplemental Figure S2. Differentially spliced and differentially expressed transcripts in mDA neuronal cells expressing the *SNCA* (A53T) mutation.** A. Differential AS events detected upon *SNCA* (A53T) mutation in differentiated DA neurons and progenitor cells (35d) using JUM (Wang and Rio, 2018). 5,001 splicing events were significantly altered in *SNCA* (A53T) mutant samples in DA neurons and progenitor cell line versus the edited wildtype control (with adjusted p-value < 0.1, ΔPSI ≥5). Data for one of three clonal mutant cell lines is shown as an example. Shown on the y-axis is the number of differential splicing events and difference in percentage-spliced-in (or ΔPSI) is shown on the x-axis. The splicing events were filtered based on the magnitude of delta PSI (ΔPSI). B. Comparison of genes that are differentially spliced in 3 different clonal lines of *SNCA* (A53T) mutant, this results in 5,001 high confidence differentially spliced genes. D2 indicates neuronal differentiation sets. C. A graph of Gene Ontology (GO) term enrichment of 5,001 differentially spliced transcripts in *SNCA* (A53T) mutant protein-expressing DA neuronal cells compared to the edited wild type control (EWT). D. Comparison of differentially spliced transcripts in *SNCA* (A53T) mutant cells to differentially spliced transcripts from genes found in Lewy body disease patient brain biopsy RNA-seq data (Feleke et al., 2021): (Parkinson disease (PD), Parkinson disease with dementia (PDD), dementia with Lewy bodies (DLB)). E. A volcano plot of D2 (differentiation set 2) is shown as an example with some up- and down-regulated gene names shown. DESeq2 analysis of the *SNCA* (A53T) mutant compared to the edited wild type (EWT) controls reveals 9,467 differentially expressed genes (adjusted p-values < 0.05, FC > 1.5). F. A graph of Gene Ontology (GO) term enrichment of either 4,708 up-regulated genes or 4,759 down-regulated genes in the *SNCA* (A53T) mutant protein-expressing mDA neuronal cells compared to the edited wild type (EWT) controls.

#### Supplemental Figure S3

**A**

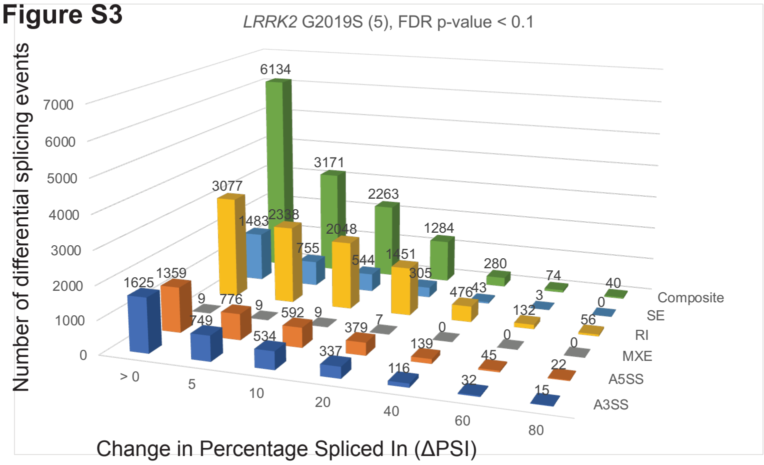

| FDR p-value < 0.1, △PSI ≥ 5 | | |
|---|---|---|
| *LRRK2* G2019S (5) | | |
| PSI >5 | Events | geneID |
| A3SS | 749 | 571 |
| A5SS | 776 | 639 |
| MXE | 9 | 5 |
| RI | 2338 | 1106 |
| SE | 755 | 681 |
| Composite | 3171 | 2623 |
| Total | 7798 | 4648 |

**B**

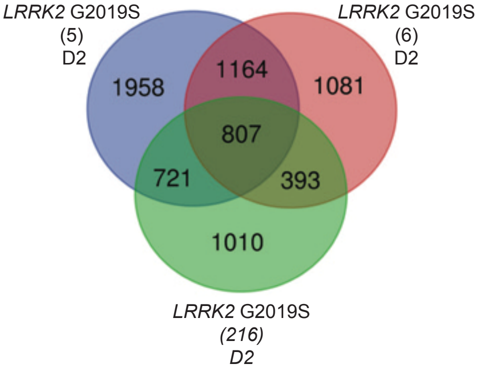

**C**

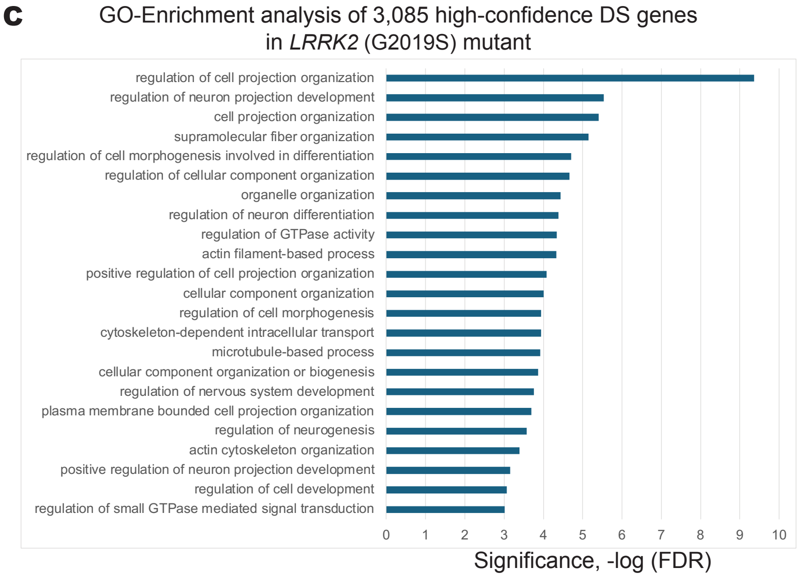

**D**

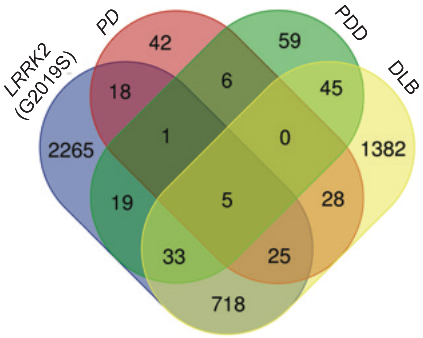

| | total | Genes |
|---|---|---|
| DLB, *LRRK2* (G2019S), PD, PDD | 5 | *SEC14L1, ADHFE1, SEC24C, SRRM2, EPB41L2* |
| *LRRK2* (G2019S), PD, PDD | 1 | *ZGPAT* |
| DLB, LRRK2 (G2019S), PD | 25 | *SNX19, UPF3A, MAST4, WDR27, GRK4, ZNF789, CALD1, UBR4, NRXN2, MACF1, PRUNE2, ABLIM2, MRPL43, EML2, CPNE5, TMEM99, AP3B1, FYN, UBE2Q2, KIDINS220, PTPRF, EVL, L3MBTL1, CADPS, CD46* |
| DLB, *LRRK2* (G2019S), PDD | 33 | *NADSYN1, ACAP3, CLIP2, ERBIN, SHTN1, ARHGAP17, RBMS2, STRADA, AP4B1, ANK2, UNK, DYNC2H1, ZDHHC11, MSI2, DLG1, FBXO42, WIPI2, VEZT, PPP1R32, FAM228B, TARBP1, CTTNBP2, TNRC6A, IKBKB, NPIPA8, COL16A1, LUC7L3, MPRIP, INO80C, ATP5SL, NT5C2, PEX5L, ST18* |
| *LRRK2* (G2019S), PD | 18 | *LRRC16B, TCFL5, SLIT2, PSD3, SLC12A5, ARHGAP6, RASGRF2, RYR1, AGAP4, SYT7, TXNL4A, TYK2, PLA2G7, ZNF91, LINC00403, ETV1, MYO5A, ZNF708* |
| *LRRK2* (G2019S), PDD | 19 | *ZNF83, GFPT2, LAMA3, CHD2, PLEKHH1, MKL2, POLG2, INPP1, ARHGAP18, XPO1, UGP2, LOC100130950, DCAF8, UBE2F-SCLY, MITD1, TMEM91, DROSHA, DOT1L, TCEA3* |

**E**

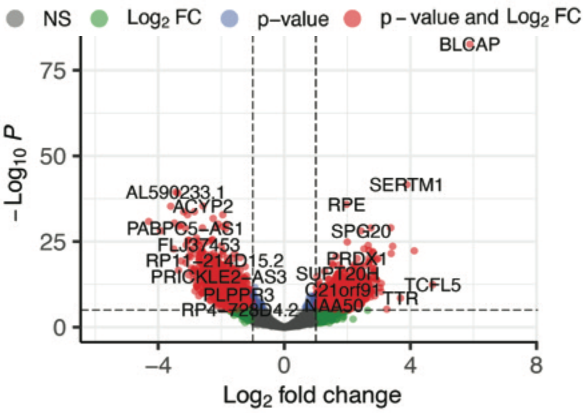

**F**

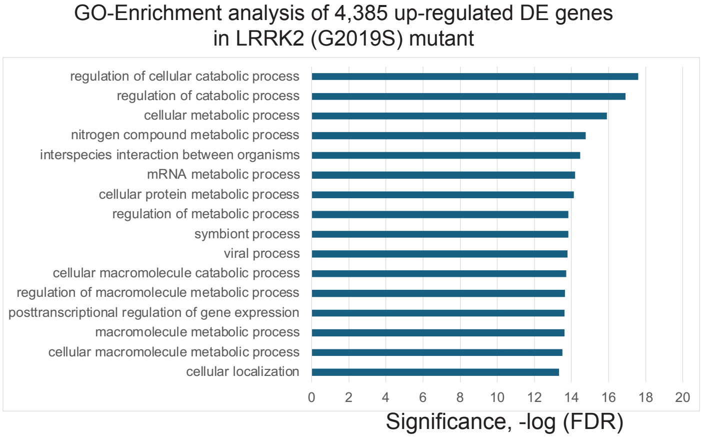

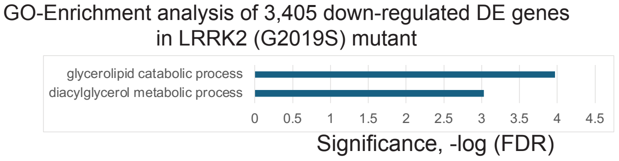

**Supplemental Figure S3. Differentially spliced and differentially expressed transcripts in mDA neuronal cells carrying the familial *LRRK2 (G2019*S) mutation.** A. Differential alternative splicing events detected upon *LRRK2* (G2019S) in differentiated DA neurons and progenitor cells (35d) using JUM (Wang and Rio, 2018). 3,085 splicing events were significantly altered in *LRRK2* (G2019S) mutant samples in DA neurons and progenitor cell line versus the edited wild type control (with adjusted p-value < 0.1, ΔPSI ≥5). Data for one of three clonal mutant cell lines is shown as an example. Shown on the y-axis is the number of differential splicing events and difference in percentage-spliced-in (or ΔPSI) shown on the x-axis. The splicing events were filtered based on the magnitude of delta PSI (ΔPSI). B. Comparison of gene transcripts that are differentially spliced in three independent clonal lines of the *LRRK2* (G2019S) mutant resulted in 3,085 high confidence differentially spliced genes. D2 indicates neuronal differentiation set. C. A graph of Gene Ontology (GO) term enrichment of 3,085 differentially spliced transcripts in *LRRK2* (G2019S) mutant protein-expressing DA neuronal cells compared to the edited wild type control (EWT). These genes are involved in regulation of cytoskeleton organization, regulation of neuron differentiation, and cilium organization. D. Comparison of differentially spliced transcripts in the *LRRK2* (G2019S) mutant cells to differentially spliced transcripts from genes found in Lewy body disease patient brain biopsy RNA-seq data (Feleke et al., 2021): (Parkinson disease (PD), Parkinson disease with dementia (PDD), dementia with Lewy bodies (DLB)). E. A volcano plot of D1 (differentiation set 2) is shown as an example with some up- and down-regulated gene names shown. DESeq2 analysis of the *LRRK2* (G2019S) mutant compared to the edited wild type (EWT) controls reveals 7,790 differentially expressed genes (adjusted p-values < 0.05, FC > 1.5). F. A graph of Gene Ontology (GO) term enrichment of either 4685 up-regulated genes or 3405 down-regulated genes in the *LRRK2* (G2019S) mutant protein-expressing mDA neuronal cells compared to the edited wild type (EWT) controls.

#### Supplemental Figure S4

**A**

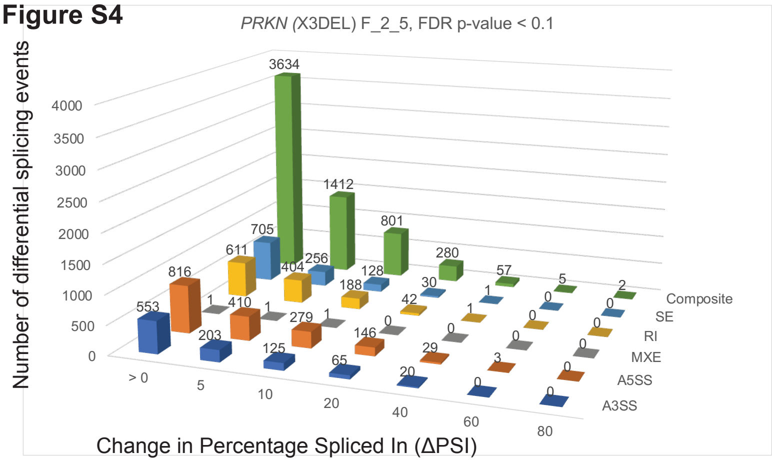

| qvalue < 0.1, PSI > 5 | | |
|---|---|---|
| *PRKN* (X3DEL) | | |
| PSI >5 | Events | geneID |
| A3SS | 203 | 149 |
| A5SS | 410 | 347 |
| MXE | 1 | 1 |
| RI | 404 | 235 |
| SE | 256 | 238 |
| Composite | 1412 | 1232 |
| Total | 2686 | 2052 |

**B**

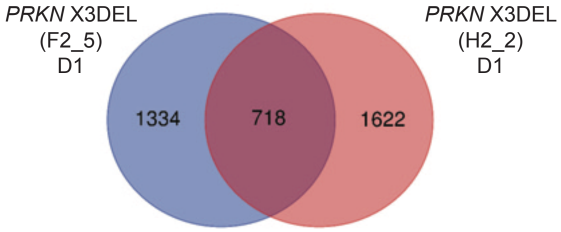

**C** GO-Enrichment analysis of high-confidence DS genes in *PRKN* (X3DEL) mutant

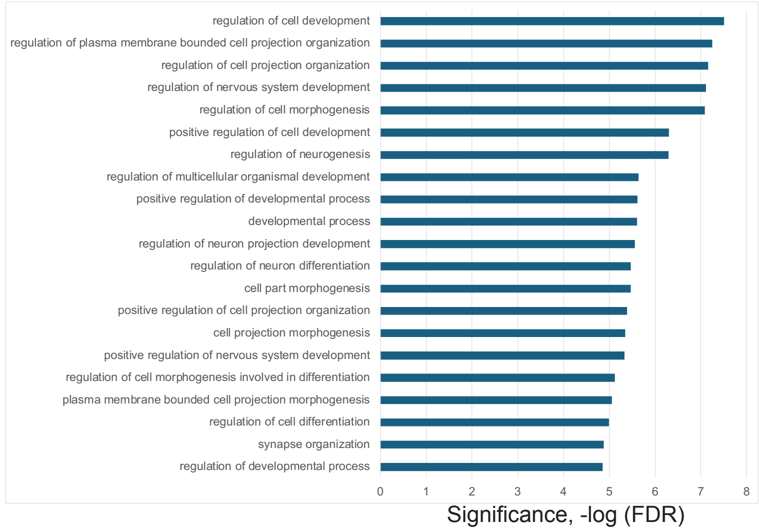

**D**

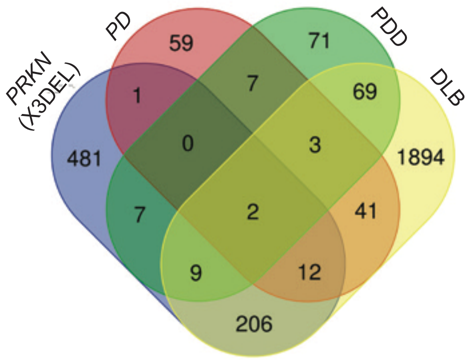

| | total | Genes |
|---|---|---|
| DLB, PD, PDD, *PRKN* (X3DEL) | 2 | *SRRM2, EPB41L2* |
| DLB, PD, *PRKN* (X3DEL) | 12 | *FYN, MAST4, KIDINS220, EML4, CALD1, EVL, NRXN2, L3MBTL1, MACF1, CADPS, DOCK10, MICAL2* |
| DLB, PDD, *PRKN* (X3DEL) | 9 | *DLG1, BRD8, PPP1R32, LINC-PINT, TNRC6A, SHTN1, RBMS2, ANK2, ATP5SL* |
| PD, *PRKN* (X3DEL) | 1 | *PSD3* |
| PDD, *PRKN* (X3DEL) | 7 | *TMEM161B-AS1, SDHAF2, TMEM91, INPP1, MAP7D2, ARHGAP18, ANK1* |

**E**

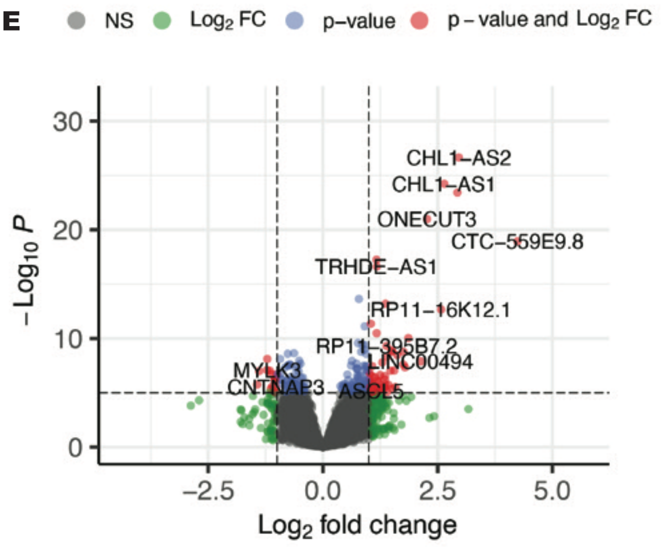

**F**

GO-Enrichment analysis of 474 up-regulated DE genes in *PRKN* (X3DEL) mutant

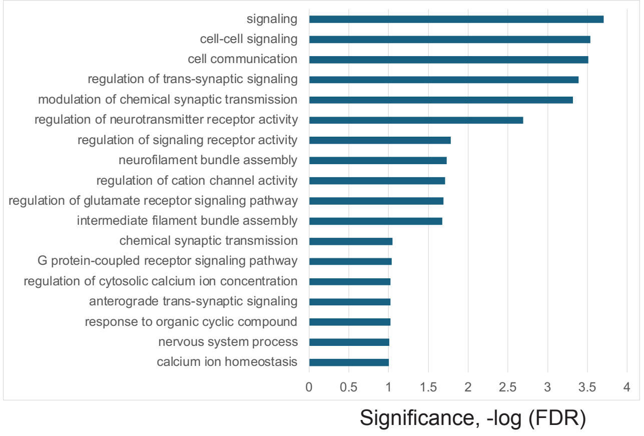

GO-Enrichment analysis of 249 down-regulated DE genes in *PRKN* (X3DEL) mutant

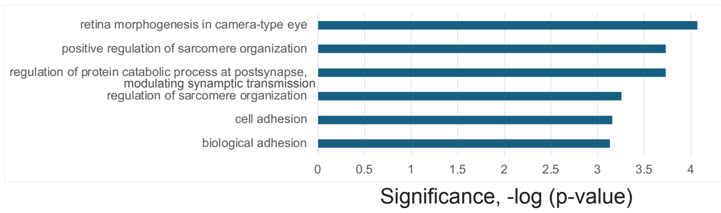

**Supplemental Figure S4. Differentially spliced and differentially expressed transcripts in mDA neuronal cells carrying the familial *PRKN* Exon 3 deletion (X3DEL) mutation.** A. Differential alternative splicing events detected upon *PRKN* exon 3 deletion in differentiated DA neurons and progenitor cells (35d) using JUM (Wang and Rio, 2018). 718 splicing events were significantly altered in the *PRKN* exon 3 deletion mutant samples in DA neurons and progenitor cells versus the edited wild type control cells (EWT, with adjusted p-value < 0.1, ΔPSI >5). Data for one of three clonal mutant cell lines is shown as an example. Shown on the y-axis is the number of differential splicing events and difference in percentage-spliced-in (or ΔPSI) is shown on the x-axis. The splicing events were filtered based on the magnitude of delta PSI (ΔPSI). B. Comparison of genes that have differentially spliced transcripts in two different clonal lines carrying the *PRKN* (X3DEL) mutation, which results in 718 high confidence differentially spliced gene transcripts. (D1 neuronal differentiation set number). C. A graph of Gene Ontology (GO) term enrichment of 718 differentially spliced transcripts in *PRKN* (X3DEL) mutant DA neuronal cells compared to the edited wild type (EWT) controls. D. Comparison and overlap of differentially spliced transcripts from genes in the *PRKN* (X3DEL) mutant cells to differentially spliced transcripts from genes found in Lewy body disease patient brain biopsy RNA-seq data (Feleke et al., 2021): (Parkinson disease (PD), Parkinson disease with dementia (PDD), dementia with Lewy bodies (DLB)). E. A volcano plot of D1 (differentiation set 1) is shown as an example with some up- and down-regulated gene names shown. DESeq2 expression analysis of the *PRKN* (X3DEL) mutant compared to the edited wild type (EWT) controls reveals 723 differentially expressed genes (adjusted p-values < 0.05, FC > 1.5). F. A graph of Gene Ontology (GO) term enrichment of either 474 up-regulated genes or 249 down-regulated genes in the *PRKN* (X3DEL) mutant protein-expressing mDA neuronal cells compared to the edited wild type (EWT) controls.

### Supplemental Figure S5

**A**

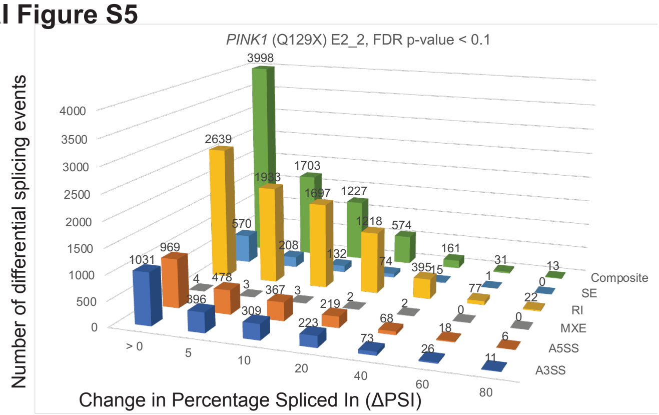

| FDR p-value < 0.1, ∆PSI ≥ 5 | | |
|---|---|---|
| *PINK1* (Q129X) E2_2 | | |
| PSI ≥ 5 | Events | geneID |
| A3SS | 396 | 278 |
| A5SS | 478 | 458 |
| MXE | 3 | 1 |
| RI | 1933 | 809 |
| SE | 208 | 186 |
| Composite | 1703 | 1455 |
| Total | 4721 | 2837 |

**B**

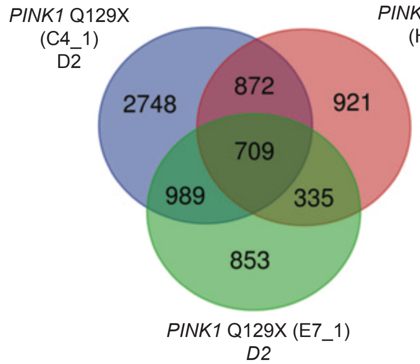

**C**

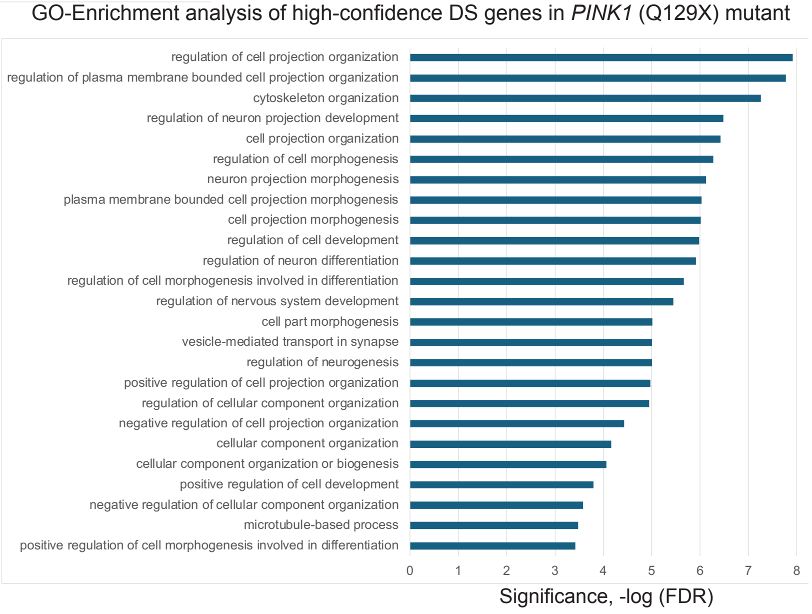

**D**

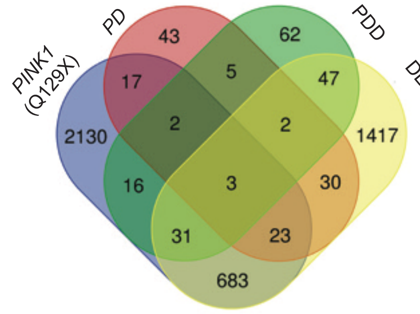

| | total | Genes |
|---|---|---|
| DLB, PD, PDD, *PINK1* (Q129X) | 3 | *SEC14L1, SRRM2, EPB41L2* |
| PD, PDD, *PINK1* (Q129X) | 2 | *PLXNB2, MEF2BNB-MEF2B* |
| DLB, PD, *PINK1* (Q129X) | 23 | *UPF3A, MAST4, WDR27, GRK4, ZNF789, CALD1, UBR4, NRXN2, MACF1, PRUNE2, ABLIM2, MRPL43, EML2, CPNE5, AP3B1, UBE2Q2, KIDINS220, PTPRF, EVL, SLC25A16, CADPS, DOCK10, MICAL2* |
| DLB, PDD, *PINK1* (Q129X) | 31 | *NADSYN1, ZFHX4-AS1, TARSL2, CLIP2, LINC-PINT, ERBIN, SGIP1, RBMS2, STRADA, ANK2, UNK, ZDHHC11, ING4, MSI2, DLG1, ZNF718, WIPI2, VEZT, PPP1R32, FAM228B, TARBP1, CTTNBP2, TNRC6A, NPIPA8, COL16A1, ACACB, PDE8A, HSF4, ODF2, NT5C2, FLJ36777* |
| PD, *PINK1* (Q129X) | 17 | *FHL1, LRRC16B, PPP6R2, SLIT2, SYNC, SLC12A5, RASGRF2, TENM4, RBFOX1, RYR1, SYT7, TYK2, LINC00403, ZCWPW1, MYO5A, RRP8, CTNNA2* |
| PDD, *PINK1* (Q129X) | 16 | *ZNF83, TMEM161B-AS1, MAPK7, LAMA3, CHD2, PLEKHH1, UBAP2, INPP1, ANK1, UGP2, LARS2, EOGT, DPY19L1P1, TMEM91, ELP2, C3orf14* |

**E**

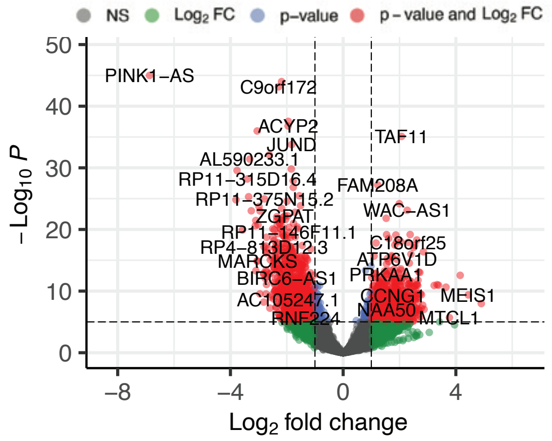

**F**

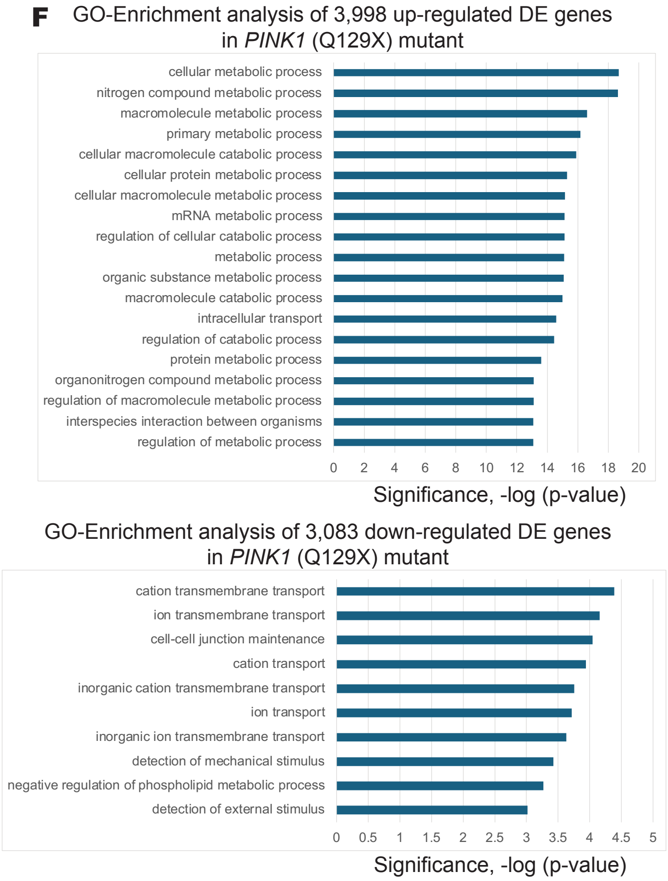

**Supplemental Figure S5. Differentially spliced and differentially expressed transcripts in mDA neuronal cells expressing the *PINK1* (Q129X) mutation.** A. Differential alternative splicing events detected upon *PINK1* (Q129X) in differentiated DA neurons and progenitor cells (35d) using JUM (Wang and Rio, 2018). 2,905 splicing events were significantly altered in *PINK1* (Q129X) mutant samples in DA neurons and progenitor cell line versus the edited wildtype control (with adjusted p-value < 0.1, ΔPSI ≥5). Data for one of three clonal mutant cell lines is shown as an example. Shown on the y-axis is the number of differential splicing events and difference in percentage-spliced-in (or ΔPSI) shown on the x-axis. The splicing events were filtered based on the magnitude of delta PSI (ΔPSI). B. Comparison of genes that are differentially spliced in 3 different clonal lines of *PINK1* (Q129X) mutant, this results in 2,905 high confidence differentially spliced genes. D2 indicates different neuronal differentiation set number. C. A graph of Gene Ontology (GO) term enrichment of 2,905 differentially spliced transcripts in *PINK1* (Q129X) mutant protein-expressing DA neuronal cells compared to the edited wild type control (EWT). D. Comparison of differentially spliced genes in *PINK1* (Q129X) mutant cells to differentially spliced transcripts from genes found in Lewy body disease patient brain biopsy RNA-seq data (Feleke et al., 2021): (Parkinson disease (PD), Parkinson disease with dementia (PDD), dementia with Lewy bodies (DLB)). E. A volcano plot of D2 (differentiation set 2) is shown as an example with some up- and down-regulated gene names shown. DESeq2 analysis of the *PINK1* (Q129X) mutant compared to the edited wild type (EWT) controls reveals 7,081 differentially expressed genes (adjusted p-values < 0.05, FC > 1.5). F. A graph of Gene Ontology (GO) term enrichment of either 3,998 up-regulated genes or 3,083 down-regulated genes in the *PINK1* (Q129X) mutant protein-expressing mDA neuronal cells compared to the edited wild type (EWT) controls.

#### Supplemental Figure S6

**A**

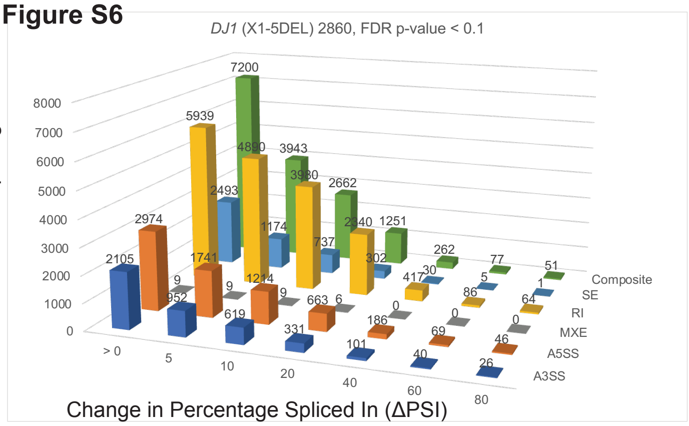

| FDR p-value < 0.1, ΔPSI ≥ 5 | | |
|---|---|---|
| *DJ1* (X1-5DEL) 2860 | | |
| PSI >5 | Events | geneID |
| A3SS | 952 | 784 |
| A5SS | 1741 | 1530 |
| MXE | 9 | 8 |
| RI | 4890 | 3047 |
| SE | 1174 | 1044 |
| Composite | 3943 | 3229 |
| Total | 12709 | 7201 |

**B**

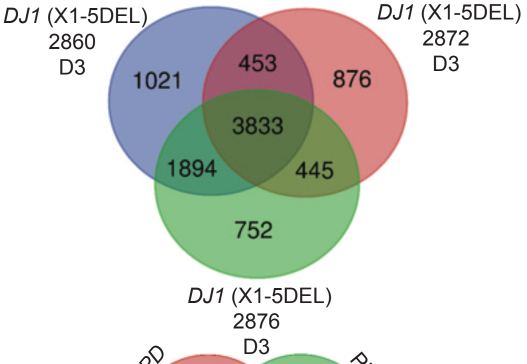

**C**

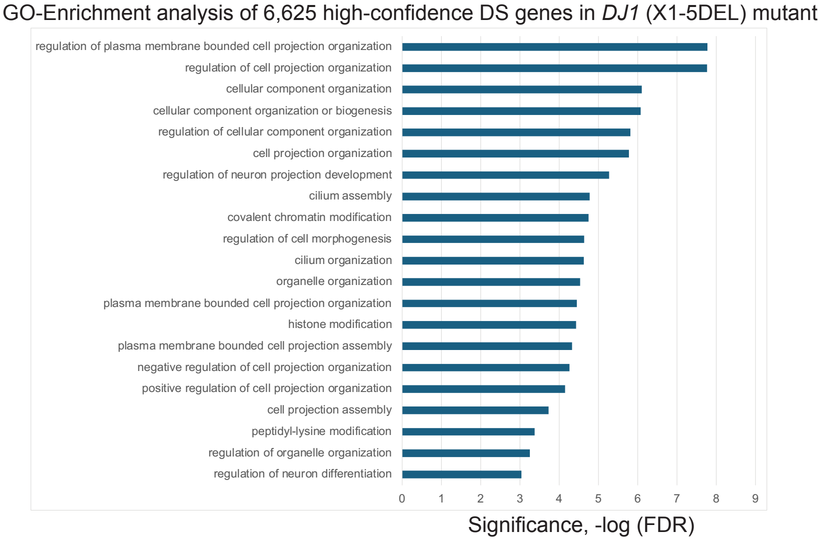

**D**

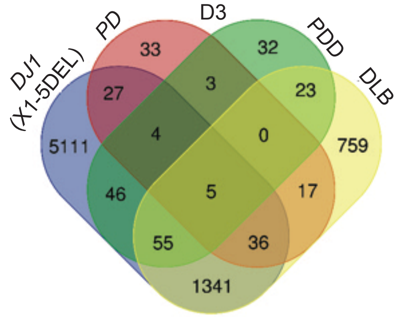

| | total | Genes |
|---|---|---|
| *DJ1* (X1-5DEL), DLB, PD, PDD | 5 | *SEC14L1, ADHFE1, SEC24C, SRRM2, EPB41L2* |
| *DJ1* (X1-5DEL), PD, PDD | 4 | *PLXNB2, MEF2BNB-MEF2B, DDX46, CCDC93* |
| *DJ1* (X1-5DEL), DLB, PD | 36 | *SNX19, UPF3A, ABHD12, MAST4, EML4, WDR27, GRK4, KIF3A, ZNF789, CALD1, UBR4, NRXN2, MACF1, PRUNE2, ABLIM2, MRPL43, EML2, B3GALNT1, TCTN2, FYN, SPIDR, UBE2Q2, KIDINS220, PTPRF, CAPS2, EVL, L3MBTL1, SCN3B, SLC25A16, CADPS, CD46, AK2, DOCK10, MICAL2, VCL, LRP2BP* |
| *DJ1* (X1-5DEL), DLB, PDD | 55 | *NADSYN1, ZFHX4-AS1, ACAP3, TARSL2, CLIP2, LINC-PINT, STK39, SLC16A4, USP33, SHTN1, SLTM, ARHGAP17, SGIP1, FAM171A1, STRADA, AP4B1, ANK2, UNK, TPRA1, ZDHHC11, DCUN1D4, ING4, MSI2, ROM1, DLG1, FBXO42, BRD8, VEZT, PPP1R32, FAM228B, TARBP1, TNRC6A, IKBKB, SYTL2, COL16A1, TMEM14B, LUC7L3, ACACB, MPRIP, MICU3, STX2, PDE8A, HSF4, TRIM9, INO80C, ODF2, ATP5SL, OPTN, PRPF18, NT5C2, FLJ36777, PEX5L, NPL, FAM49B, ST18* |
| *DJ1* (X1-5DEL), PD | 27 | *FHL1, LRRC16B, TCFL5, PPP6R2, IKBKAP, FMNL1, C19orf18, DUS1L, SLIT2, PSD3, ANKZF1, SYNC, SLC12A5, RASGRF2, WDR59, RBFOX1, WBSCR22, ZNF781, SYT7, TYK2, DCHS2, ZCWPW1, TAMM41, MYO5A, NSUN2, RRP8, CYB561D1* |
| *DJ1* (X1-5DEL), PDD | 46 | *HAUS2, ZNF83, GFPT2, TMEM161B-AS1, NOXA1, MAPK7, LAMA3, CHD2, PLEKHH1, MKL2, TMEM218, POLR2J3, PANK2, UBAP2, POLG2, CYP4V2, UBR5, INPP1, TIGD7, ARHGAP18, PREB, OGFOD3, XPO1, ANK1, GLYR1, KIAA1191, EOGT, NPC1, DCAF8, UBE2F-SCLY, ALDH3A2, DRAM2, SPG11, KIAA0586, SDHAF2, TMEM91, TIRAP, ELP2, C3orf14, CACNG7, ESD, DROSHA, DOT1L, TCEA3, GPR89A, SNX17* |

**E**

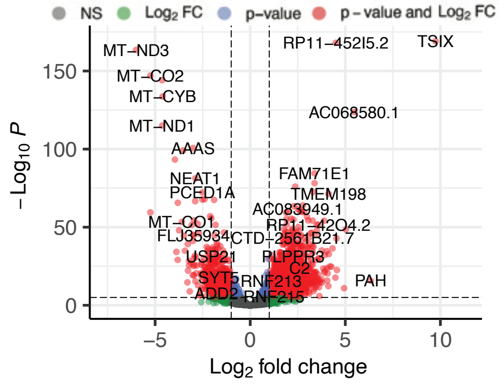

**F**

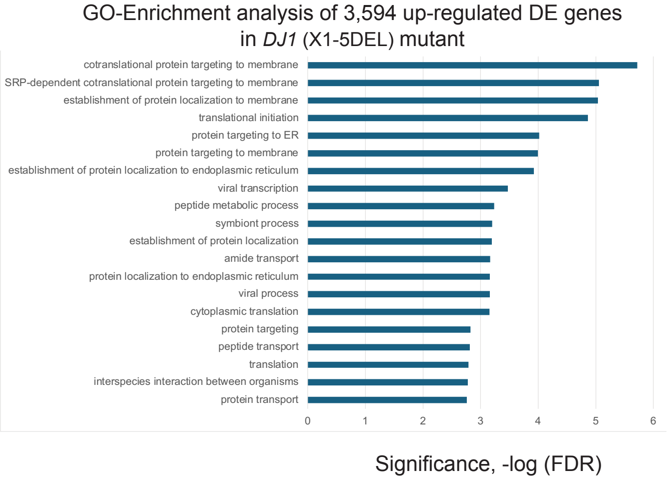

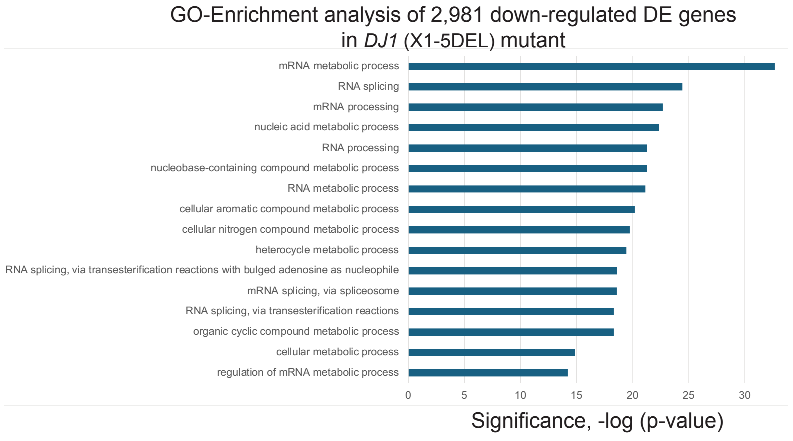

**Supplemental Figure S6. Differentially spliced and differentially expressed transcripts in mDA neuronal cells expressing the *DJ1* (X1-5DEL) mutation.** A. Differential alternative splicing (AS) events detected in *DJ1* (X1-5DEL) mutant differentiated mDA neurons and progenitor cells (35d). Over 6,500 splicing events were significantly altered in *DJ1* (X1-5DEL) mutant samples of mDA neurons and progenitor cells versus the edited wild type control cells (with adjusted p-value < 0.1, ΔPSI ≥5). Data for one clonal line is shown as an example. Shown on the y-axis is the number of differential splicing events and difference in percentage spliced in (or ΔPSI) shown on the x-axis. The splicing events were filtered based on the magnitude of delta PSI (ΔPSI). B. Comparison of transcripts from genes that are differentially spliced in 2 different clonal lines of the *DJ1* (X1-5DEL) mutant, which results in 6,625 high confidence differentially spliced gene transcripts. D3 indicates neuronal differentiation set number. C. A graph of Gene Ontology (GO) term enrichment of 6,625 differentially spliced transcripts in *DJ1* (X1-5DEL) mutant protein-expressing mDA neuronal cells compared to the edited wild type control cells. D. Comparison of differentially spliced transcripts in the *DJ1* (X1-5DEL) mutant cells to differentially spliced transcripts found in diseased patient brains (Parkinson disease (PD), Parkinson disease with dementia (PDD), dementia with Lewy bodies (DLB)). E. A volcano plot of D3 (differentiation set 3) is shown as an example with some up- and down-regulated gene names shown. DESeq2 analysis of the *DJ1* (X1-5DEL) mutant compared to the edited wild type (EWT) controls reveals 6,575 differentially expressed genes (adjusted p-values < 0.05, FC > 1.5). F. A graph of Gene Ontology (GO) term enrichment of either 3,594 up-regulated genes or 2,981 down-regulated genes in the *DJ1* (X1-5DEL) mutant protein-expressing mDA neuronal cells compared to the edited wild type (EWT) controls.

#### Supplemental Figure S7

**A**

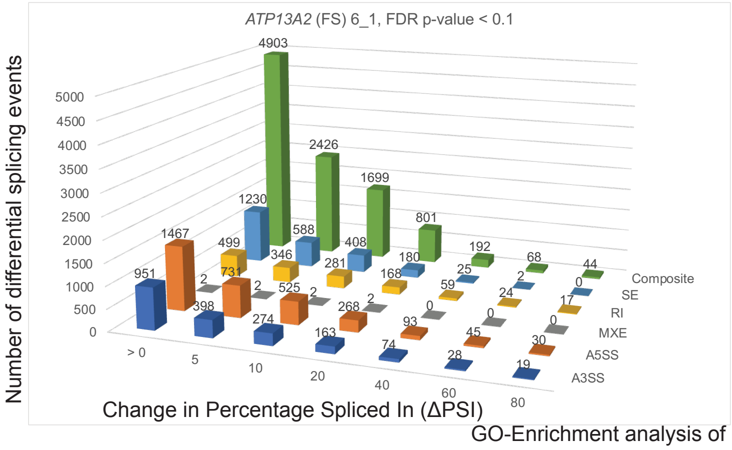

| FDR p-value < 0.1, ΔPSI ≥ 5 | | |
|---|---|---|
| *ATP13A2* (FS) 6_1 | | |
| PSI >5 | Events | geneID |
| A3SS | 398 | 320 |
| A5SS | 731 | 634 |
| MXE | 2 | 2 |
| RI | 346 | 282 |
| SE | 588 | 536 |
| Composite | 2426 | 2056 |
| Total | 4491 | 3380 |

**B**

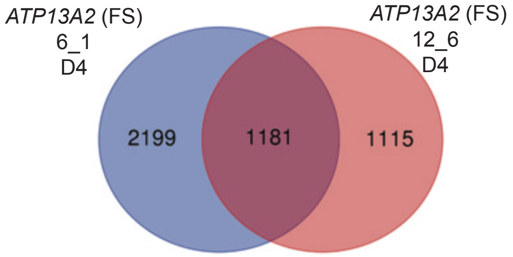

GO-Enrichment analysis of 1,181 high-confidence DS genes in *ATP13A2* (FS) mutant

**C**

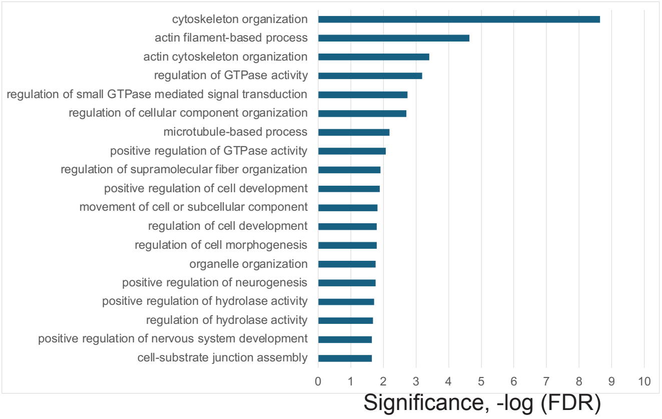

**D**

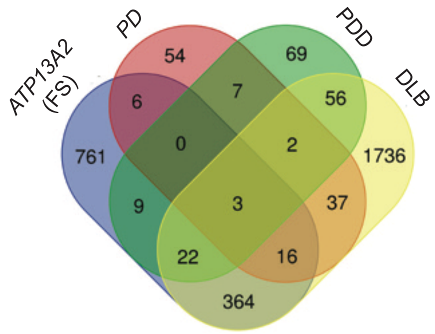

| | total | Genes |
|---|---|---|
| *ATP13A2* (FS), DLB, PD, PDD | 3 | *SEC14L1, SRRM2, EPB41L2* |
| *ATP13A2 (FS)* , DLB, PD | 16 | *UPF3A, MAST4, KIDINS220, PTPRF, EML4, WDR27, KIF3A, CALD1, L3MBTL1, MACF1, CD46, AK2, ABLIM2, EML2, IPO11, LRP2BP* |
| *ATP13A2 (FS)* , DLB, PDD | 22 | *DLG1, ZFHX4-AS1, VEZT, CLIP2, PPP1R32, TARBP1, CTTNBP2, TNRC6A, ERBIN, COL16A1, SHTN1, MPRIP, ARHGAP17, STX2, STRADA, PDE8A, ANK2, TRIM9, DYNC2H1, INO80C, ATP5SL, NT5C2* |
| *ATP13A2 (FS)* , PD | 6 | *TENM4, RBFOX1, FHL1, SYNC, RRP8, RASGRF2* |
| *ATP13A2 (FS)* , PDD | 9 | *DPY19L1P1, SDHAF2, XRN1, TMEM218, MAP7D2, ESD, C4orf33, SNX17, DCAF8* |

**E**

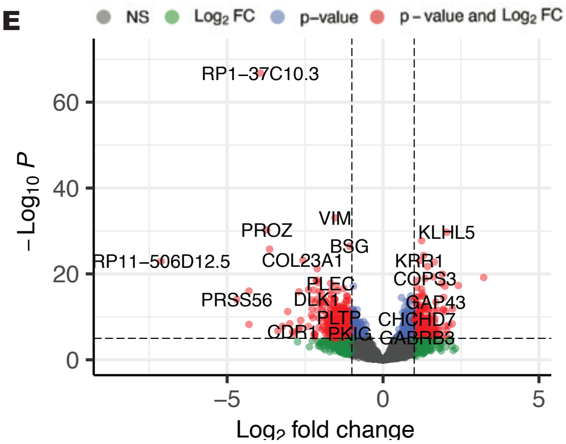

**F**

GO-Enrichment analysis of 2,266 up-regulated DE genes in *ATP13A2* (FS) mutant

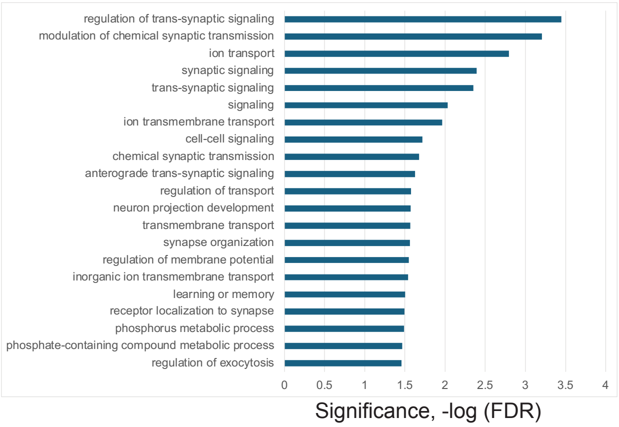

GO-Enrichment analysis of 1,943 down-regulated DE genes in *ATP13A2* (FS) mutant

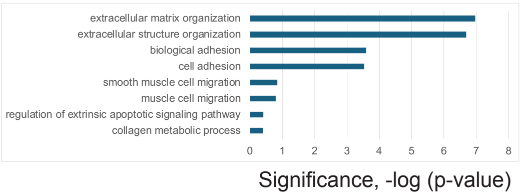

**Supplemental Figure S7. Differentially spliced and differentially expressed transcripts in mDA neuronal cells expressing the *ATP13A2* (FS) mutation.** A. Differential alternative splicing (AS) events detected upon *ATP13A2* (FS) mutant differentiated mDA neurons and progenitor cells (35d). 1,111 splicing events were significantly altered in *ATP13A2* (FS) mutant samples of mDA neurons and progenitor cell line versus the edited wildtype control (with adjusted p-value < 0.1, ΔPSI ≥5). Data for one clonal line is shown as an example. Shown on the y-axis is the number of differential splicing events and difference in percentage spliced in (or ΔPSI) shown on the x-axis. The splicing events were filtered based on the magnitude of delta PSI (ΔPSI). B. Comparison of transcripts from genes that are differentially spliced in 2 different clonal lines of the *ATP13A2* (FS) mutant, which results in 1,181 high confidence differentially spliced gene transcripts. D4 indicates neuronal differentiation set number. C. A graph of Gene Ontology (GO) term enrichment of 1,181 differentially spliced transcripts in *ATP13A2* (FS) mutant protein-expressing mDA neuronal cells compared to the edited wild type control cells. D. Comparison of differentially spliced transcripts in *ATP13A2* (FS) mutant cells to differentially spliced transcripts found in diseased patient brains (Parkinson disease (PD), Parkinson disease with dementia (PDD), dementia with Lewy bodies (DLB)). E. A volcano plot of D4 (differentiation set 4) is shown as an example with some up- and down-regulated gene names shown. DESeq2 analysis of the *ATP13A2* (FS) mutant compared to the edited wild type (EWT) controls reveals 4,209 differentially expressed genes (adjusted p-values < 0.05, FC > 1.5). F. A graph of Gene Ontology (GO) term enrichment of either 2,266 up-regulated genes or 1,943 down-regulated genes in the *ATP13A2* (FS) mutant protein-expressing mDA neuronal cells compared to the edited wild type (EWT) controls.

#### Supplemental Figure S8

**A**

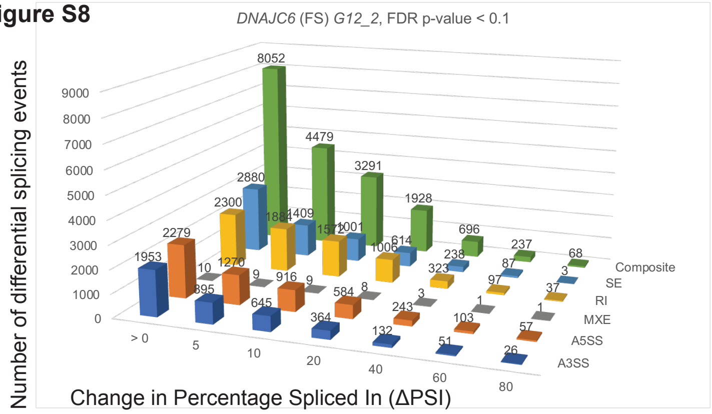

| *FDR p-value < 0.1, ΔPSI ≥ 5* | | |
|---|---|---|
| *DNAJC6* (FS) G12_2 | | |
| PSI >5 | Events | geneID |
| A3SS | 895 | 782 |
| A5SS | 1270 | 1087 |
| MXE | 9 | 9 |
| RI | 1884 | 1256 |
| SE | 1409 | 1202 |
| Composite | 4479 | 3610 |
| Total | 9946 | 6084 |

**B**

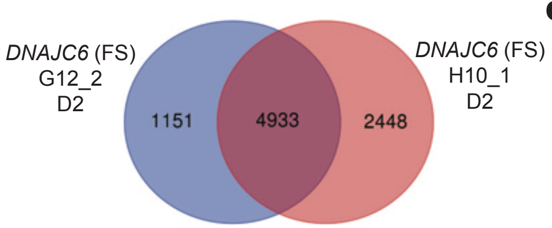

**C** GO-Enrichment analysis of 4,933 high-confidence DS genes in *DNAJC6* (FS) mutant

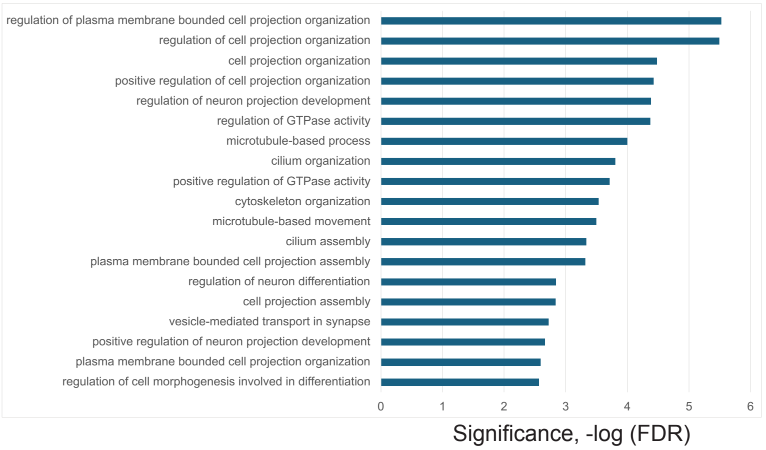

**D**

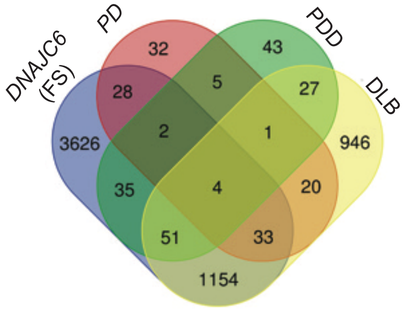

| | total | Genes |
|---|---|---|
| DLB, *DNAJC6* (FS), PD, PD | 4 | *SEC14L1, SEC24C, SRRM2, EPB41L2* |
| *DNAJC6* (FS), PD, PDD | 2 | *PLXNB2, MEF2BNB-MEF2B* |
| DLB, *DNAJC6* (FS), PD | 33 | *SNX19, UPF3A, MAST4, EML4, WDR27, GRK4, KIF3A, ZNF789, CALD1, UBR4, MACF1, PRUNE2, ABLIM2, MRPL43, CPNE5, TMEM99, AP3B1, FYN, UBE2Q2, KIDINS220, PTPRF, GABRG2, EVL, L3MBTL1, LPGAT1, CADPS, CD46, AK2, APIP, DENND3, DOCK10, MICAL2, LRP2BP* |
| DLB, *DNAJC6* (FS), PDD | 51 | *NADSYN1, ZFHX4-AS1, TARSL2, CLIP2, LINC-PINT, STK39, BBS9, MYO9A, USP33, ERBIN, SHTN1, SLTM, ARHGAP17, SGIP1, RBMS2, STRADA, ANK2, UNK, DYNC2H1, TPRA1, ZDHHC11, DCUN1D4, MSI2, ROM1, DLG1, FBXO42, ZNF718, BRD8, WIPI2, VEZT, PPP1R32, FAM228B, TARBP1, TNRC6A, IKBKB, SYTL2, LUC7L3, MPRIP, MICU3, STX2, PDE8A, HSF4, TRIM9, INO80C, ODF2, ATP5SL, NT5C2, PEX5L, FAM49B, ST18, PIK3IP1* |
| *DNAJC6* (FS), PD | 28 | *FHL1, TCFL5, FMNL1, ZCCHC2, DUS1L, SLIT2, PSD3, ANKZF1, SYNC, VARS2, SLC12A5, ARHGAP6, RASGRF2, TENM4, RBFOX1, WBSCR22, RYR1, SYT7, TYK2, ZNF91, LINC00403, ETV1, ZCWPW1, MYO5A, RRP8, SCN2A, CYB561D1, CTNNA2* |
| *DNAJC6* (FS), PDD | 35 | *ZNF83, GFPT2, TMEM161B-AS1, MAPK7, CHD2, PLEKHH1, MKL2, TMEM218, UBAP2, POLG2, ZFHX4, MAP7D2, C4orf33, TIGD7, ARHGAP18, PREB, OGFOD3, XPO1, ANK1, KIAA1191, UGP2, STARD9, EOGT, NPC1, DCAF8, UBE2F-SCLY, ALDH3A2, DRAM2, SPG11, CCDC113, DPY19L1P1, TMEM91, ELP2, ESD, DOT1L* |

**E**

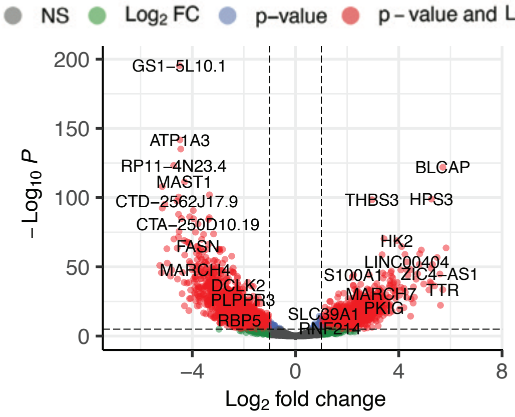

**F**

GO-Enrichment analysis of 3,763 up-regulated DE genes in *DNAJC6* (FS) mutant

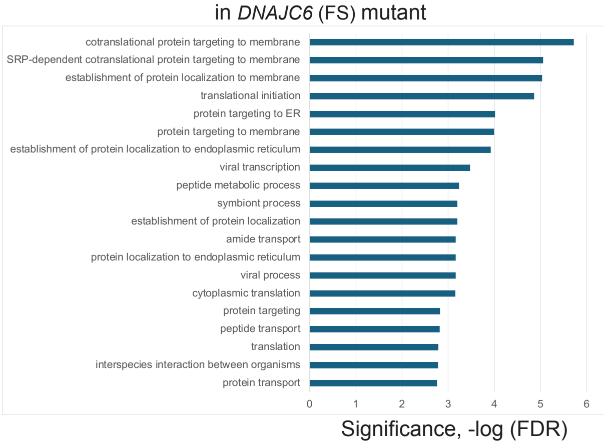

GO-Enrichment analysis of 3,620 down-regulated DE genes in *DNAJC6* (FS) mutant

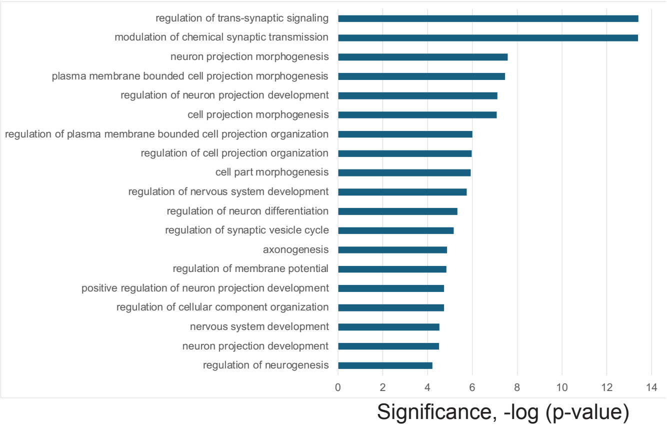

**Supplemental Figure S8. Differentially spliced and differentially expressed transcripts in mDA neuronal cells expressing the *DNAJC6* (FS) mutation.** A. Differential alternative splicing (AS) events detected in *DNAJC6* (FS) mutant differentiated mDA neurons and progenitor cells (35d). ~5,000 splicing events were significantly altered in *DNAJC6* (FS) mutant samples of mDA neurons and progenitor cells versus the edited wild type control cells (with adjusted p-value < 0.1, ΔPSI ≥5). Data for one clonal line is shown as an example. Shown on the y-axis is the number of differential splicing events and difference in percentage spliced in (or ΔPSI) shown on the x-axis. The splicing events were filtered based on the magnitude of delta PSI (ΔPSI). B. Comparison of transcripts from genes that are differentially spliced in 2 different clonal lines of the *DNAJC6* (FS) mutant, which results in 4,933 high confidence differentially spliced gene transcripts. D2 indicates neuronal differentiation set number. C. A graph of Gene Ontology (GO) term enrichment of 4,933 differentially spliced transcripts in *DNAJC6* (FS) mutant protein-expressing mDA neuronal cells compared to the edited wild type control cells. D. Comparison of differentially spliced transcripts in the *DNAJC6* (FS) mutant cells to differentially spliced transcripts found in diseased patient brains (Parkinson disease (PD), Parkinson disease with dementia (PDD), dementia with Lewy bodies (DLB)). E. A volcano plot of D1 (differentiation set 1) is shown as an example with some up- and down-regulated gene names shown. DESeq2 analysis of the *DNAJC6* (FS) mutant compared to the edited wild type (EWT) controls reveals 7,383 differentially expressed genes (adjusted p-values < 0.05, FC > 1.5). F. A graph of Gene Ontology (GO) term enrichment of either 3,763 up-regulated genes or 3,620 down-regulated genes in the *DNAJC6* (FS) mutant protein-expressing mDA neuronal cells compared to the edited wild type (EWT) controls.

#### Supplemental Figure S9

**A**

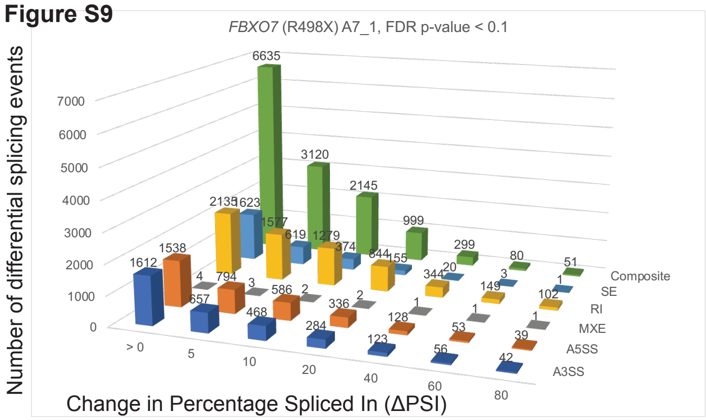

| FDR p-value < 0.1, ΔPSI ≥ 5 | | |
|---|---|---|
| FBXO7 (R498X) A7_1 | | |
| PSI >5 | Events | geneID |
| A3SS | 657 | 532 |
| A5SS | 794 | 701 |
| MXE | 3 | 2 |
| RI | 1577 | 1198 |
| SE | 619 | 558 |
| Composite | 3120 | 2648 |
| Total | 6770 | 4752 |

**B**

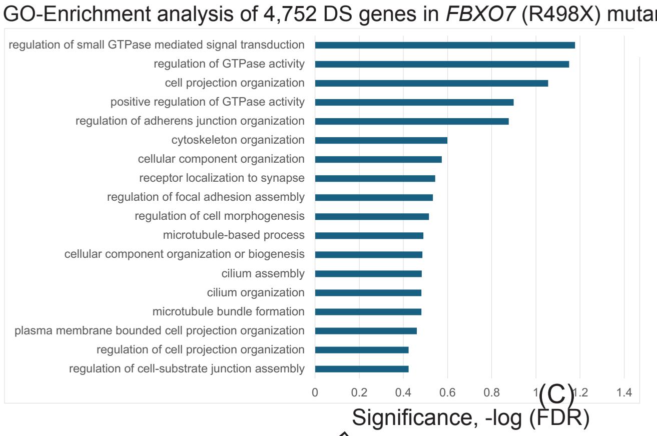

**D**

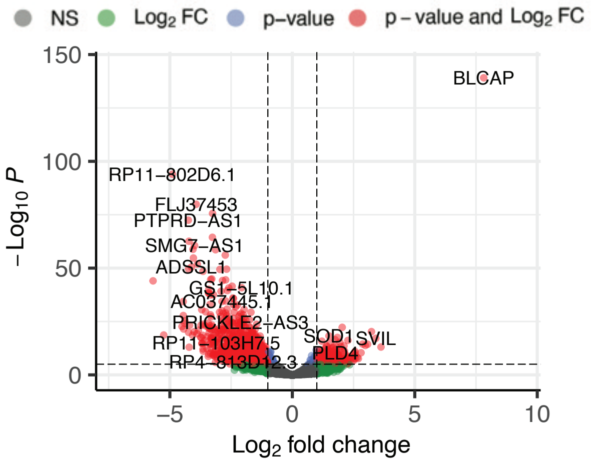

**C**

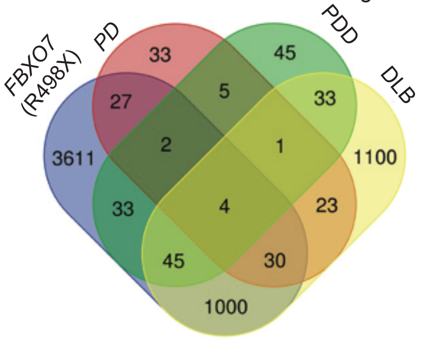

| | total | Genes |
|---|---|---|
| DLB, *FBXO7* (R498X), PD, PDD | 4 | *SEC14L1, ADHFE1, SRRM2, EPB41L2* |
| *FBXO7* (R498X), PD, PDD | 2 | *MEF2BNB-MEF2B, ZGPAT* |
| DLB, *FBXO7* (R498X), PD | 30 | *SNX19, UPF3A, MAST4, EML4, WDR27, GRK4, ZNF789, CALD1, UBR4, C1orf162, NRXN2, MACF1, PRUNE2, ABLIM2, EML2, B3GALNT1, CPNE5, AP3B1, FYN, UBE2Q2, KIDINS220, PTPRF, EVL, L3MBTL1, LPGAT1, SLC25A16, CD46, DENND3, MICAL2, LRP2BP* |
| DLB, *FBXO7* (R498X), PDD | 45 | *NADSYN1, ZFHX4-AS1, ACAP3, TARSL2, LINC-PINT, BBS9, MYO9A, USP33, ERBIN, SHTN1, SLTM, ARHGAP17, RBMS2, STRADA, AP4B1, ANK2, UNK, DYNC2H1, ZDHHC11, ING4, MSI2, ROM1, DLG1, FBXO42, ZNF718, VEZT, PPP1R32, FAM228B, TARBP1, CTTNBP2, TNRC6A, NPIPA8, SYTL2, COL16A1, LUC7L3, MPRIP, STX2, PDE8A, HSF4, TRIM9, NT5C2, PEX5L, NPL, FAM49B, ST18* |
| *FBXO7* (R498X), PD | 27 | *PPP6R2, IKBKAP, SLIT2, PSD3, ANKZF1, SLC12A5, ARHGAP6, RELN, RASGRF2, TENM4, BBS7, WDR59, WBSCR22, RYR1, SYT7, TXNL4A, GABRA1, PLA2G7, ZNF91, LINC00403, ETV1, ZCWPW1, MYO5A, RRP8, SCN2A, ZNF708, CYB561D1* |
| *FBXO7* (R498X), PDD | 33 | *HAUS2, PDLIM2, ZNF83, TMEM161B-AS1, MAPK7, CHD2, PLEKHH1, MKL2, TMEM218, POLR2J3, PANK2, POLG2, MAP7D2, TIGD7, PREB, XPO1, ANK1, KIAA1191, UGP2, STARD9, LOC100130950, DCAF8, UBE2F-SCLY, SPG11, TNFSF12, MITD1, DPY19L1P1, TMEM91, LINC01170, TIRAP, RNF8, DROSHA, DOT1L* |

**E**

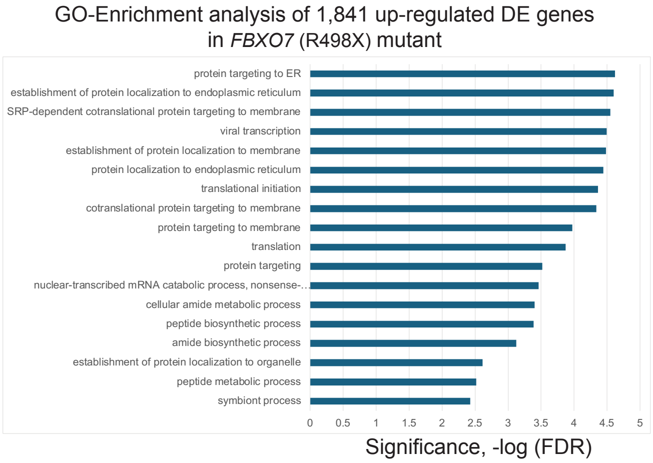

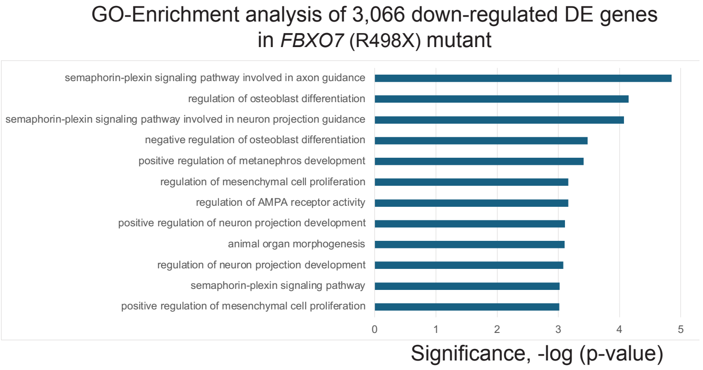

**Supplemental Figure S9. Differentially spliced and differentially expressed transcripts in mDA neuronal cells expressing the *FBXO7* (R498X) mutation.** A. Differential alternative splicing (AS) events detected in *FBXO7* (R498X) mutant differentiated mDA neurons and progenitor cells (35d). Over 4,500 splicing events were significantly altered in *FBXO7* (R498X) mutant samples of mDA neurons and progenitor cells versus the edited wild type control cells (with adjusted p-value < 0.1, ΔPSI ≥5). Data for one clonal cell line is shown. Shown on the y-axis is the number of differential splicing events and difference in percentage spliced in (or ΔPSI) shown on the x-axis. The splicing events were filtered based on the magnitude of delta PSI (ΔPSI). B. A graph of Gene Ontology (GO) term enrichment of 4,506 differentially spliced transcripts in *FBXO7* (R498X) mutant protein-expressing DA neuronal cells against the edited wild type control cells. C. Comparison of differentially spliced transcripts in the *FBXO7* (R498X) mutant cell line to differentially spliced transcripts found in diseased patient brains (Parkinson disease (PD), Parkinson disease with dementia (PDD), dementia with Lewy bodies (DLB)). D. A volcano plot of D2 (differentiation set 2) is shown as an example with some up- and down-regulated gene names shown. DESeq2 analysis of the *FBXO7* (R498X) mutant compared to the edited wild type (EWT) controls reveals 4,907 differentially expressed genes (adjusted p-values < 0.05, FC > 1.5). E. A graph of Gene Ontology (GO) term enrichment of either 1,841 up-regulated genes or 3,066 down-regulated genes in the *FBXO7* (R498X) mutant protein-expressing mDA neuronal cells compared to the edited wild type (EWT) controls.

#### Supplemental Figure S10

**A**

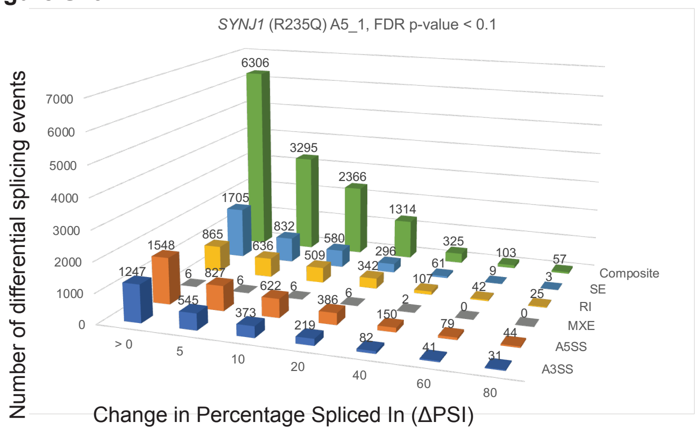

| FDR p-value < 0.1, ∆PSI ≥ 5 | | |
|---|---|---|
| *SYNJ1* (R235Q) A5_1 | | |
| PSI >5 | Events | geneID |
| A3SS | 545 | 441 |
| A5SS | 827 | 714 |
| MXE | 6 | 5 |
| RI | 636 | 522 |
| SE | 832 | 751 |
| Composite | 3295 | 2751 |
| Total | 6141 | 4481 |

**B**

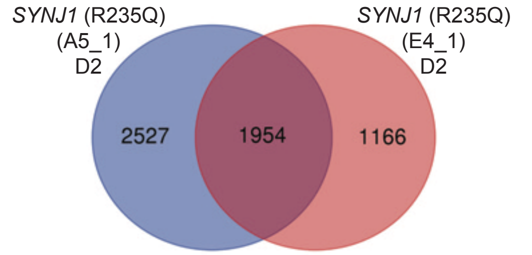

**C**

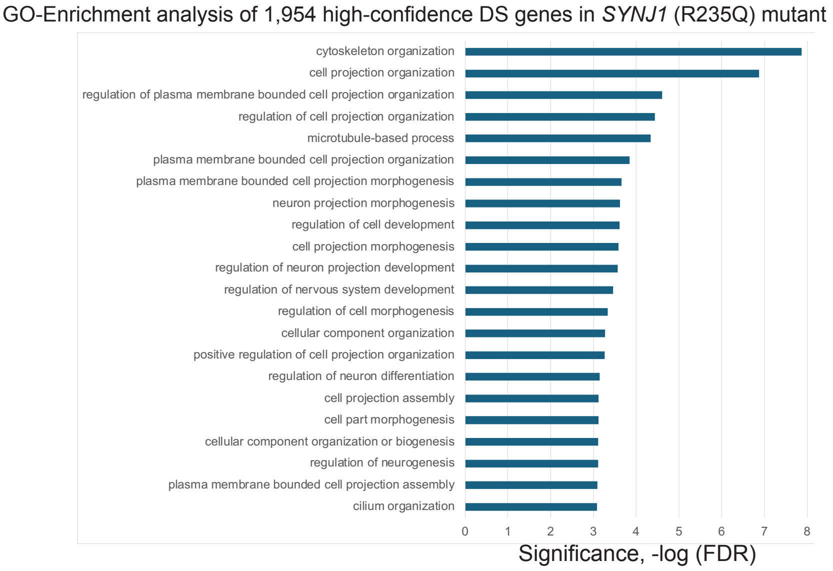

**D**

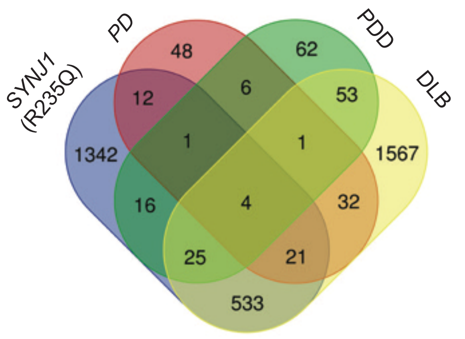

| | total | Genes |
|---|---|---|
| DLB, PD, PDD, *SYNJ1* (R235Q) | 4 | *SEC14L1, SRRM2, EPB41L2, SEC24C* |
| PD, PDD, *SYNJ1* (R235Q) | 1 | *MEF2BNB-MEF2B* |
| DLB, PD, *SYNJ1* (R235Q) | 21 | *CPNE5, AP3B1, FYN, MAST4, KIDINS220, PTPRF, CAPS2, GRK4, KIF3A, ZNF789, CALD1, EVL, MACF1, SLC25A16, CADPS, CD46, AK2, MRPL43, EML2, DENND3, MICAL2* |
| DLB, PDD, *SYNJ1* (R235Q) | 25 | *DLG1, FBXO42, ZNF718, ZFHX4-AS1, WIPI2, VEZT, CLIP2, PPP1R32, LINC-PINT, FAM228B, TARBP1, MYO9A, COL16A1, SHTN1, ARHGAP17, STX2, STRADA, AP4B1, ANK2, HSF4, TRIM9, DYNC2H1, ODF2, FAM49B, ST18* |
| PD, *SYNJ1* (R235Q) | 12 | *RBFOX1, RYR1, FHL1, FMNL1, TYK2, ZNF91, SLIT2, PSD3, ZCWPW1, RRP8, ZNF708, RASGRF2* |
| PDD, *SYNJ1* (R235Q) | 16 | *UBE2F-SCLY, ZNF83, TMEM161B-AS1, MITD1, CCDC113, MAPK7, PLEKHH1, POLR2J3, ELP2, MAP7D2, C4orf33, DOT1L, ARHGAP18, ANK1, KIAA1191, TCEA3* |

**E**

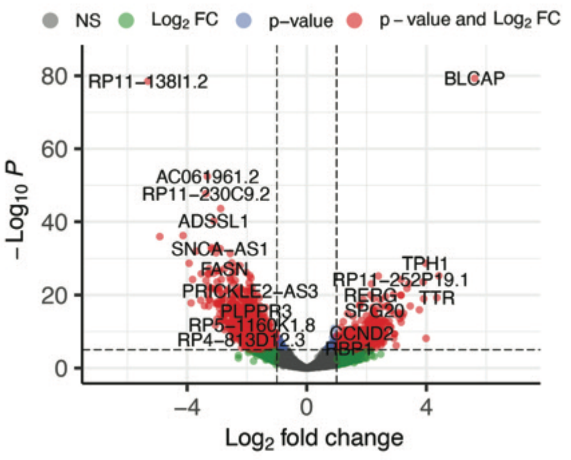

**F**

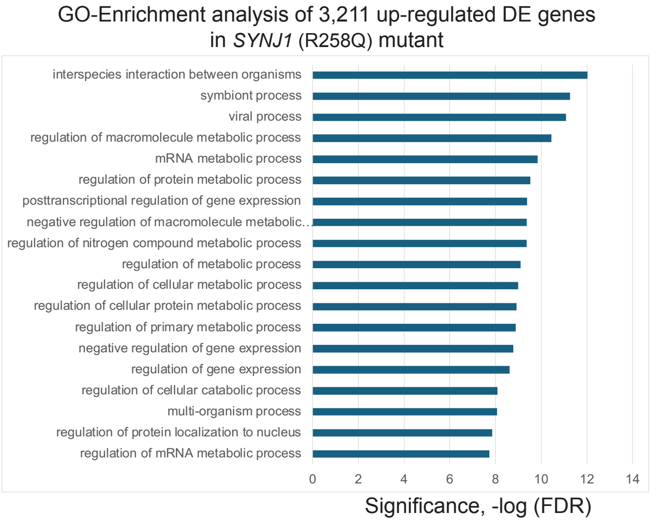

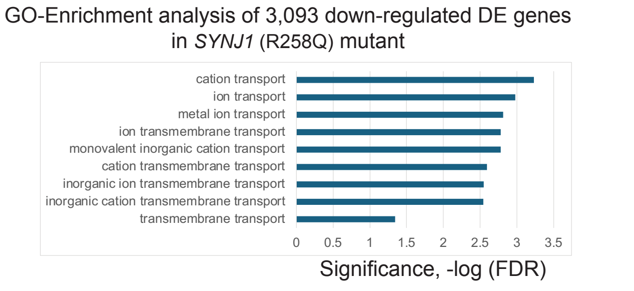

**Supplemental Figure S10. Differentially spliced and differentially expressed transcripts in mDA neuronal cells expressing the *SYNJ1* (R235Q) mutation.** A. Differential alternative splicing (AS) events detected in *SYNJ1* (R235Q) mutant differentiated mDA neurons and progenitor cells (35d). Over 1,900 splicing events were significantly altered in *SYNJ1* (R235Q) mutant samples of mDA neurons and progenitor cell lines versus the edited wild type control cells (with adjusted p-value < 0.1, ΔPSI ≥5). Data for one clonal cell line is shown as an example. Shown on the y-axis is the number of differential splicing events and difference in percentage spliced in (or ΔPSI) shown on the x-axis. The splicing events were filtered based on the magnitude of delta PSI (ΔPSI). B. Comparison of transcripts from genes that are differentially spliced in 2 different clonal lines of the *SYNJ1* (R235Q) mutant, which results in 1,954 high confidence differentially spliced gene transcripts. D2 indicates the neuronal differentiation set number. C. A graph of Gene Ontology (GO) term enrichment of 1,954 differentially spliced transcripts in *SYNJ1* (R235Q) mutant protein-expressing mDA neuronal cells compared to the edited wild type control cells. D. Comparison of differentially spliced transcripts in the *SYNJ1* (R235Q) mutant cells to differentially spliced transcripts found in diseased patient brains (Parkinson disease (PD), Parkinson disease with dementia (PDD), dementia with Lewy bodies (DLB)). E. A volcano plot of D2 (differentiation set 2) is shown as an example with some up- and down-regulated gene names shown. DESeq2 analysis of the *SYNJ1* (R258Q) mutant compared to the edited wild type (EWT) controls reveals 6304 differentially expressed genes (adjusted p-values < 0.05, FC > 1.5). F. A graph of Gene Ontology (GO) term enrichment of either 3,211 up-regulated genes or 3,093 down-regulated genes in the *SYNJ1* (R258Q) mutant protein-expressing mDA neuronal cells compared to the edited wild type (EWT) controls.

#### Supplemental Figure S11

**A**

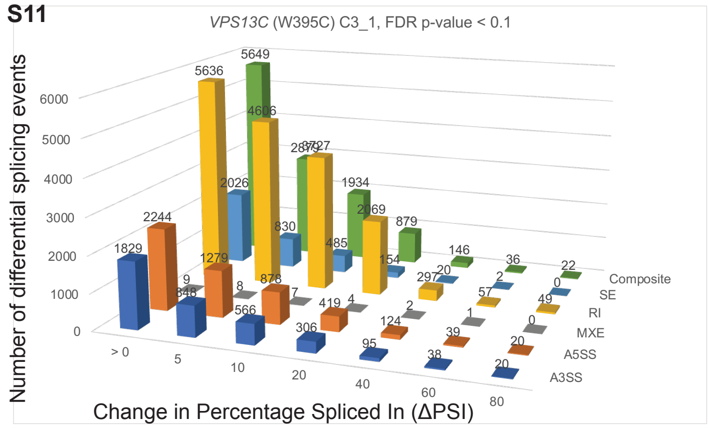

| FDR p-value < 0.1, ΔPSI ≥ 5 | | |
|---|---|---|
| *VPS13C* (W395C) C3_1 | | |
| PSI >5 | Events | geneID |
| A3SS | 848 | 715 |
| A5SS | 1279 | 1135 |
| MXE | 8 | 8 |
| RI | 4606 | 2979 |
| SE | 830 | 759 |
| Composite | 2879 | 2440 |
| Total | 10450 | 6230 |

**B**

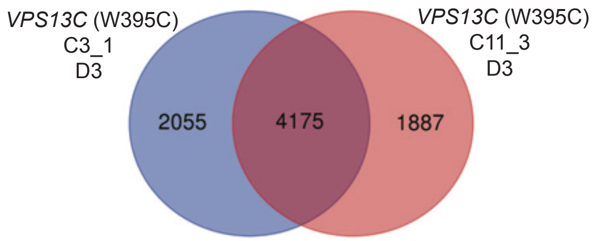

**C** GO-Enrichment analysis of 4,175 high-confidence DS genes in *VPS13C* (W395C) mutant

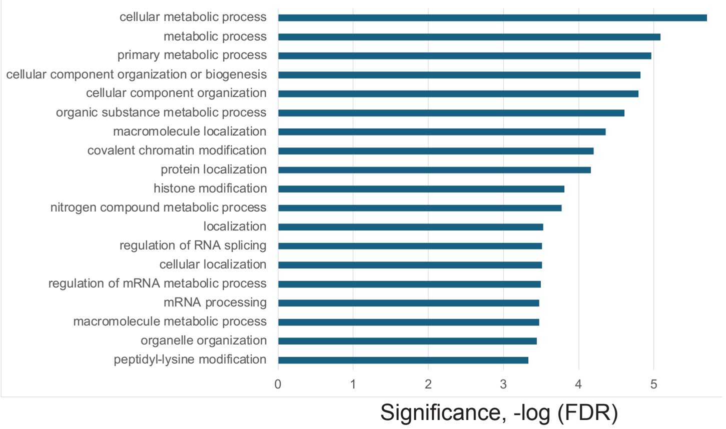

**D**

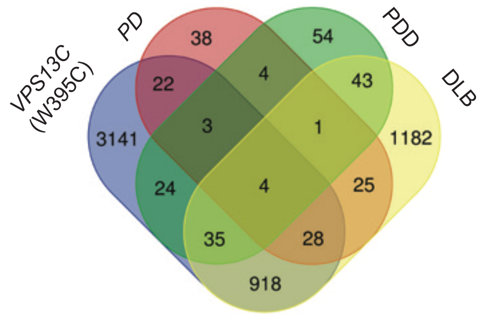

| | total | Genes |
|---|---|---|
| DLB, PD, PDD, *VPS13C* (W395C) | 4 | *SEC14L1, SEC24C, SRRM2, EPB41L2* |
| PD, PDD, *VPS13C* (W395C) | 3 | *PLXNB2, MEF2BNB-MEF2B, DDX46* |
| DLB, PD, *VPS13C* (W395C) | 28 | *SNX19, UPF3A, ABHD12, MAST4, EML4, WDR27, GRK4, ZNF789, CALD1, UBR4, NRXN2, MACF1, PRUNE2, ABLIM2, EML2, B3GALNT1, TCTN2, FYN, SPIDR, PTPRF, SCN3B, SLC25A16, CADPS, CD46, AK2, MICAL2, VCL, LRP2BP* |
| DLB, PDD, *VPS13C* (W395C) | 35 | *NADSYN1, ZFHX4-AS1, ACAP3, SLC16A4, USP33, SLTM, ARHGAP17, SGIP1, FAM171A1, AP4B1, ANK2, UNK, TPRA1, ZDHHC11, DCUN1D4, ING4, ROM1, DLG1, FBXO42, BRD8, PPP1R32, FAM228B, TARBP1, TNRC6A, IKBKB, COL16A1, TMEM14B, LUC7L3, ACACB, PDE8A, HSF4, INO80C, ODF2, OPTN, FAM49B* |
| PD, *VPS13C* (W395C) | 22 | *FHL1, LRRC16B, TCFL5, PPP6R2, IKBKAP, DUS1L, SLIT2, ANKZF1, SYNC, SLC12A5, ECD, RBFOX1, WBSCR22, TXNL4A, TYK2, ZNF91, LINC00403, DCHS2, TAMM41, MYO5A, NSUN2, RRP8* |
| PDD, *VPS13C* (W395C) | 24 | *TMEM161B-AS1, MAPK7, CHD2, MKL2, TMEM218, UBAP2, POLG2, CYP4V2, INPP1, ARHGAP18, PREB, XPO1, GLYR1, STARD9, NPC1, DCAF8, ALDH3A2, DRAM2, KIAA0586, TMEM91, DROSHA, DOT1L, GPR89A, SNX17* |

**E**

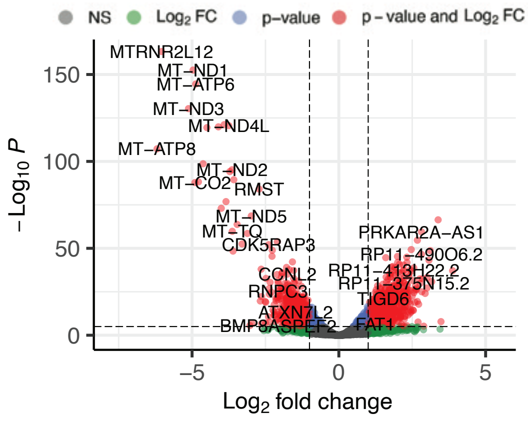

**F** GO-Enrichment analysis of 3,249 up-regulated DE genes in *VPS13C* (W395C) mutant

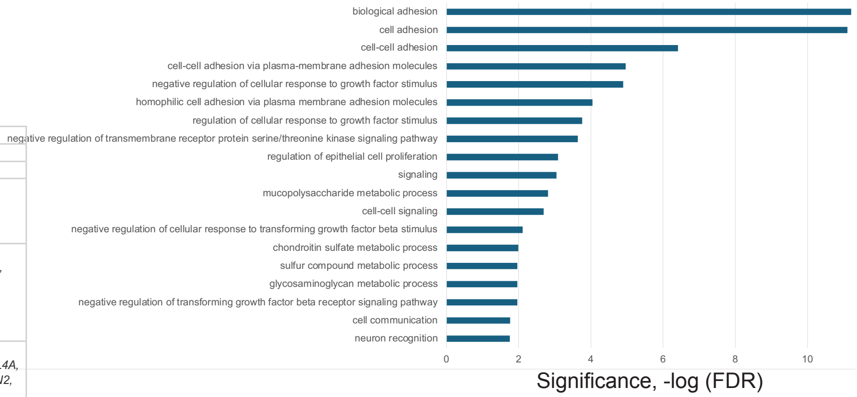

GO-Enrichment analysis of 2,830 down-regulated DE genes in *VPS13C* (W395C) mutant

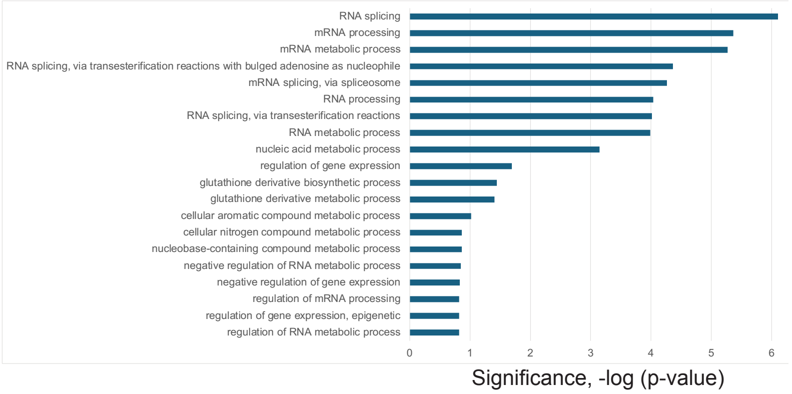

**Supplemental Figure S11. Differentially spliced and differentially expressed transcripts in mDA neuronal cells expressing the *VPS13C* (W395C) mutation.** A. Differential alternative splicing (AS) events detected in *VPS13C* (W395C) mutant differentiated mDA neurons and progenitor cells (35d). Over 4,000 splicing events were significantly altered in *VPS13C* (W395C) mutant samples of mDA neurons and progenitor cell line versus the edited wildtype control (with adjusted p-value < 0.1, ΔPSI ≥5). Data for one clonal line is shown as an example. Shown on the y-axis is the number of differential splicing events and difference in percentage spliced in (or ΔPSI) shown on the x-axis. The splicing events were filtered based on the magnitude of delta PSI (ΔPSI). B. Comparison of transcripts from genes that are differentially spliced in 2 different clonal lines of *VPS13C* (W395C) mutant, which results in 4,175 high confidence differentially spliced gene transcripts. D3 indicates neuronal differentiation set number. C. A graph of Gene Ontology (GO) term enrichment of 4,175 differentially spliced transcripts in *VPS13C* (W395C) mutant protein-expressing mDA neuronal cells compared to the edited wild type control cells. D. Comparison of differentially spliced transcripts in the *VPS13C* (W395C) mutant cells to differentially spliced transcripts found in diseased patient brains (Parkinson disease (PD), Parkinson disease with dementia (PDD), dementia with Lewy bodies (DLB)). E. A volcano plot of D3 (differentiation set 3) is shown as an example with some up- and down-regulated gene names shown. DESeq2 analysis of the *VPS13C* (W395C) mutant compared to the edited wild type (EWT) controls reveals 6,079 differentially expressed genes (adjusted p-values < 0.05, FC > 1.5). F. A graph of Gene Ontology (GO) term enrichment of either 3,249 up-regulated genes or 2,830 down-regulated genes in the *VPS13C* (W395C) mutant protein-expressing mDA neuronal cells compared to the edited wild type (EWT) controls.

#### Supplemental Figure S12

**A**

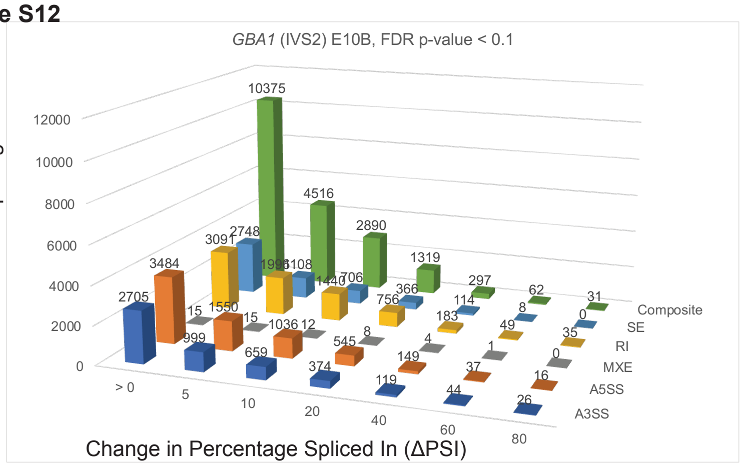

| FDR p-value < 0.1, ΔPSI ≥ 5 | | |
|---|---|---|
| *GBA1* (IVS2) E10B | | |
| PSI >5 | Events | geneID |
| A3SS | 999 | 736 |
| A5SS | 1550 | 1269 |
| MXE | 15 | 13 |
| RI | 1996 | 1415 |
| SE | 1108 | 918 |
| Composite | 4516 | 3510 |
| Total | 10184 | 6107 |

**B**

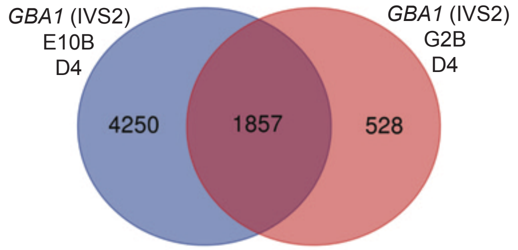

**C**

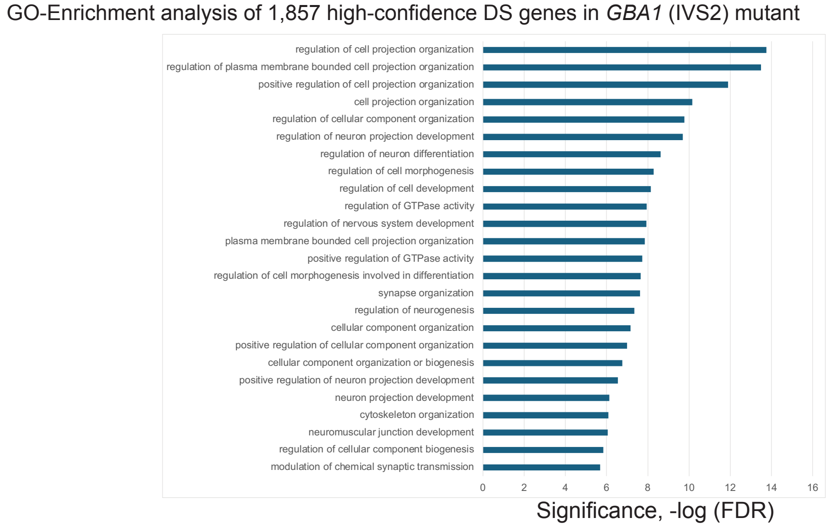

**D**

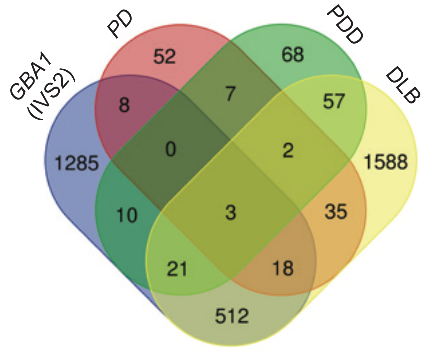

| | total | Genes |
|---|---|---|
| DLB, *GBA1 (IVS2)*, PD, PDD | 3 | *SEC14L1, SRRM2, EPB41L2* |
| DLB, *GBA1* (IVS2), PD | 18 | *MAST4, KIDINS220, PTPRF, EML4, KIF3A, ZNF789, UBR4, EVL, NRXN2, L3MBTL1, MACF1, CADPS, CD46, PRUNE2, ABLIM2, DOCK10, MICAL2, LRP2BP* |
| DLB, *GBA1* (IVS2), PDD | 21 | *DLG1, ZFHX4-AS1, TARSL2, CLIP2, TARBP1, MYO9A, TNRC6A, COL16A1, SHTN1, MPRIP, ARHGAP17, STX2, SGIP1, ANK2, HSF4, INO80C, ZDHHC11, ATP5SL, ROM1, NPL, FAM49B* |
| *GBA1* (IVS2), PD | 8 | *TENM4, RBFOX1, RYR1, FHL1, TXNL4A, DCHS2, MYO5A, SLC12A5* |
| *GBA1* (IVS2), PDD | 10 | *UBE2F-SCLY, KIAA0586, LAMA3, TMEM218, POLR2J3, UBAP2, ZFHX4, MAP7D2, XPO1, DCAF8* |

**E**

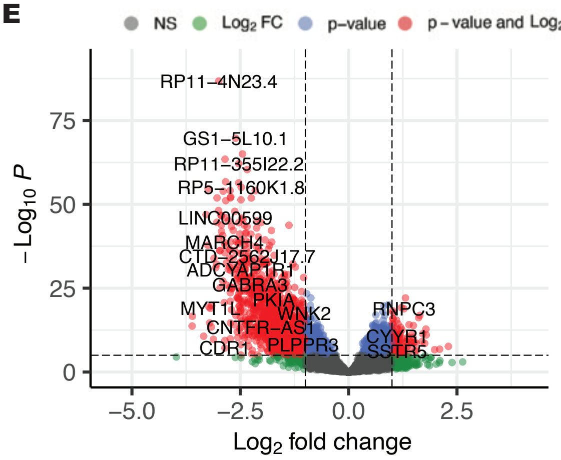

**F**

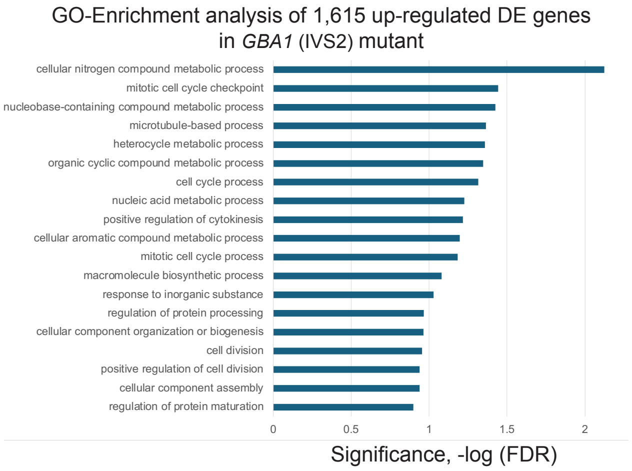

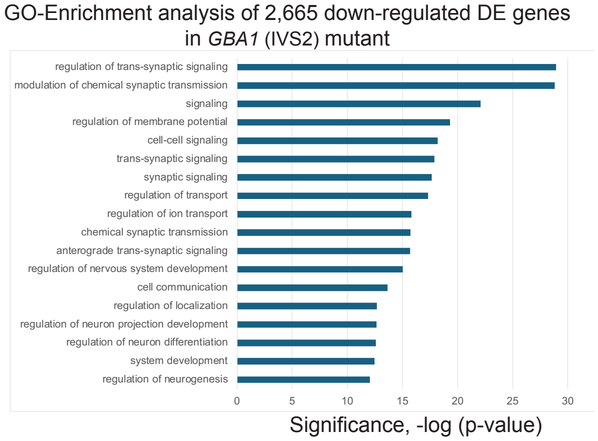

**Supplemental Figure S12. Differentially spliced and differentially expressed transcripts in mDA neuronal cells expressing the *GBA1* (IVS2) mutation.** A. Differential alternative splicing (AS) events detected in *GBA1* (IVS2) mutant differentiated mDA neurons and progenitor cells (35d). Over 1800 splicing events were significantly altered in *GBA1* (IVS2) mutant samples of mDA neurons and progenitor cell line versus the edited wild type control cells (with adjusted p-value < 0.1, ΔPSI ≥5). Data for one clonal line is shown as an example. Shown on the y-axis is the number of differential splicing events and difference in percentage spliced in (or ΔPSI) shown on the x-axis. The splicing events were filtered based on the magnitude of delta PSI (ΔPSI). B. Comparison of transcripts from genes that are differentially spliced in 2 different clonal lines of the *GBA1* (IVS2) mutant, which results in 1,857 high confidence differentially spliced gene transcripts. D4 indicates neuronal differentiation set number. C. A graph of Gene Ontology (GO) term enrichment of 1,857 differentially spliced transcripts in *GBA1* (IVS2) mutant protein-expressing mDA neuronal cells compared to the edited wild type control cells. D. Comparison of differentially spliced transcripts in the *GBA1* (IVS2) mutant cells to differentially spliced transcripts found in diseased patient brains (Parkinson disease (PD), Parkinson disease with dementia (PDD), dementia with Lewy bodies (DLB)). E. A volcano plot of D4 (differentiation set 4) is shown as an example with some up- and down-regulated gene names shown. DESeq2 analysis of the *GBA1* (IVS2) mutant compared to the edited wild type (EWT) controls reveals 4,280 differentially expressed genes (adjusted p-values < 0.05, FC > 1.5). F. A graph of Gene Ontology (GO) term enrichment of either 3,249 up-regulated genes or 2,830 down-regulated genes in the *GBA1* (IVS2) mutant protein-expressing mDA neuronal cells compared to the edited wild type (EWT) controls.

#### Supplemental Figure S13

(A)

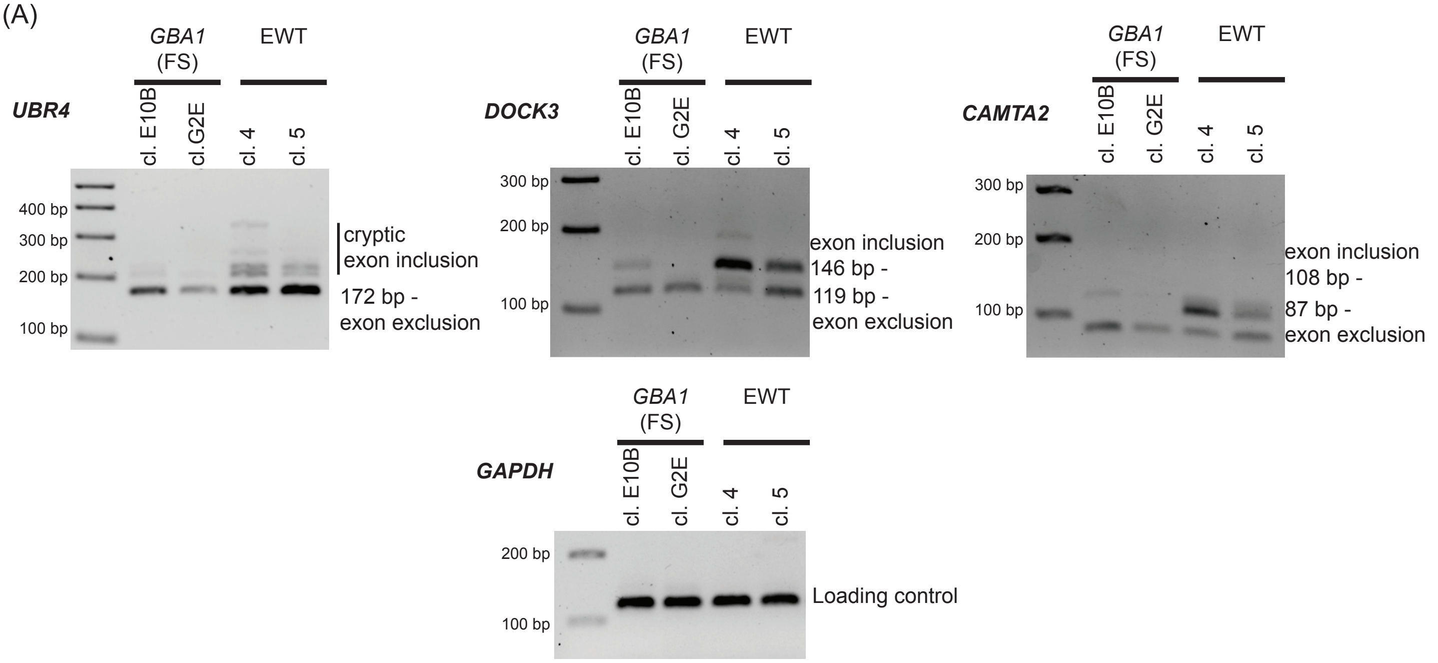

(B)

| Primers | | Genomic Coordinates (hg38): | ΔPSI (mut-EWT) |
|---|---|---|---|
| UBR4-F | ccttctgcctctgtcagcaa | chr1:19148626-19150575 | -0.35 [GBA1 (FS) E10B]<br>-0.29 [GBA1 (FS) G2E] |
| UBR4-R | tctcgatgattgggccaacg | | |
| DOCK3-F | ggaggccgagttgattgaca | chr3:51330223-51333156 | -0.59 [GBA1 (FS) E10B]<br>-0.47 [GBA1 (FS) G2E] |
| DOCK3-R | cgcgccatgtttcttgttca | | |
| CAMTA2-F | cagcgctgttaccggaagta | chr17:4969337-4969628 | -0.58 [GBA1 (FS) E10B]<br>-0.36 [GBA1 (FS) G2E] |
| CAMTA2-R | gcttcggaacttgctctgga | | |
| GAPDH-F | ctctgctcctcctgttcgac | | |
| GAPDH-R | gcgcccaatacgaccaaatc | | |

**Supplemental Figure S13. Validations of differential splicing events by RT-PCR.** (A) 0.5 μg of total RNA was reverse-transcribed according to the manufacturer's instructions (Bio-Rad, 1708891) and subjected to RT–PCR using primers listed in (B). *UBR4*, *DOCK3*, and *CAMPTA2* are shown as an example from *GBA1* (FS) mut and EWT control cells. *GAPDH* serves as a loading control. (B) Primer information used in RT-PCR validation.

**Supplemental Figure S14**

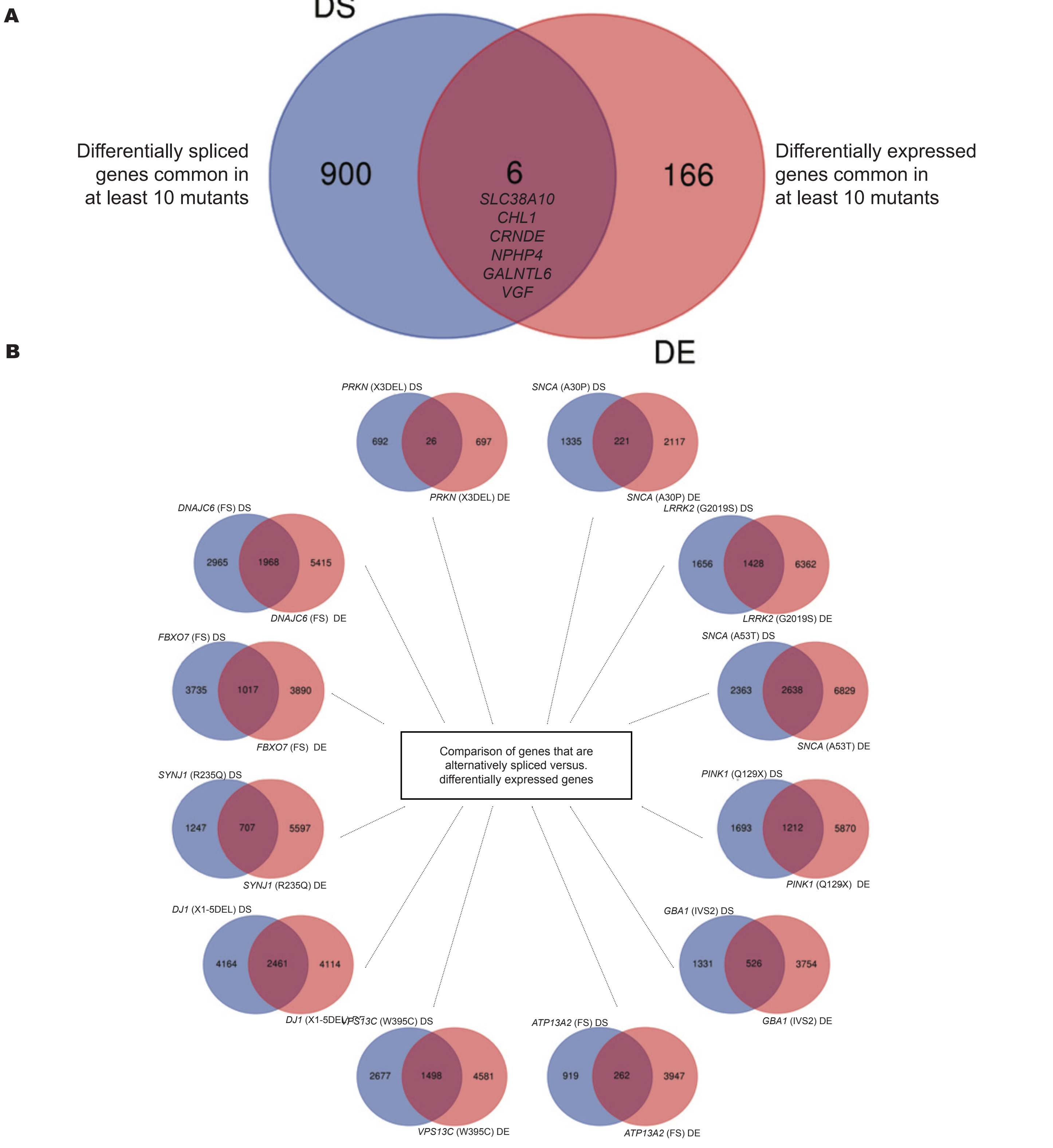

**Supplemental Figure S14. Alternative splicing and gene expression are subject to distinct and generally non-overlapping regulation.** (A) Comparison of differentially spliced (DS) versus differentially expressed (DE) gene transcripts that are common in at least 10 familial PD-associated mutants. 906 genes with significant splicing alterations were detected in 10 more distinct familial PD mutants, while 172 genes with altered expression patterns are detected in common. (B) Venn diagrams showing overlap between genes whose transcripts are differentially spliced (DS) compared to differentially expressed (DE) in familial PD-associated mutant differentiated mDA neurons. A substantial number of splicing variations are observed without corresponding changes in overall gene expression at the transcript level.

#### Supplemental Figure S15

**A**

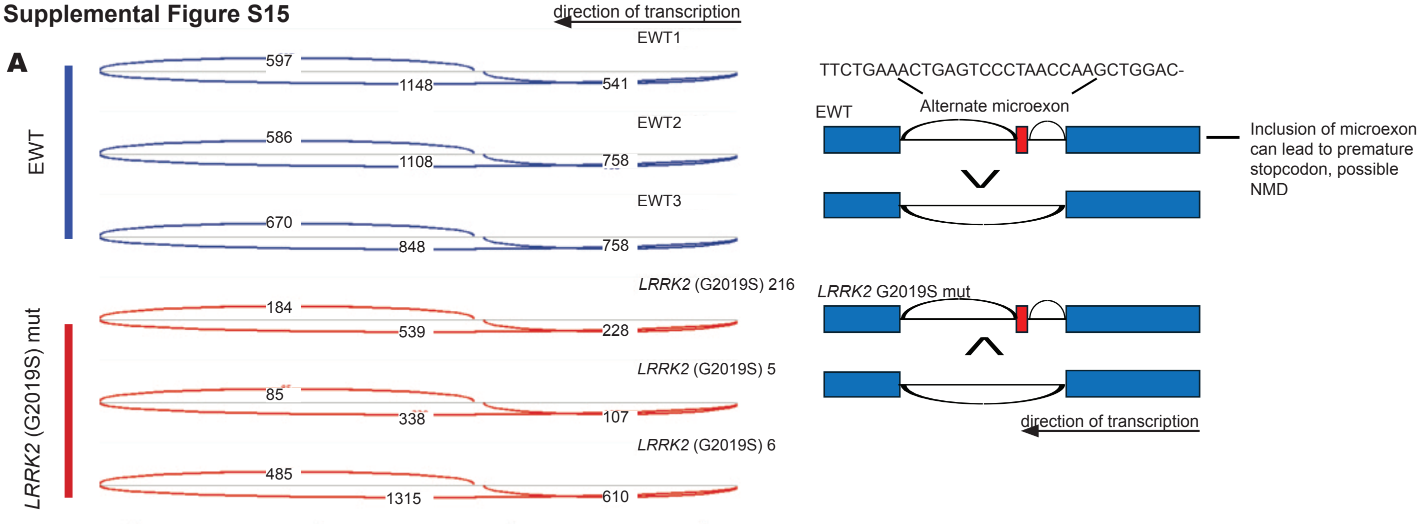

**B**

| PD mutant | Splicing mode | Genomic Coordinates | ΔPSI (EWT-PD mut) |
|---|---|---|---|
| ***DJ1*** **(X1_5DEL) 2860** | RI | chr1:19104666-19105046 | 0.18 |
| ***DJ1*** **(X1_5DEL) 2872** | RI | chr1:19110464-19110740 | 0.18 |
| ***DJ1*** **(X1_5DEL) 2876** | RI | chr1:19104666-19105046 | 0.28 |
| ***DNAJC6*** **(FS) G12_2** | SE | chr1:19148626-19150575 | 0.47 |
| ***DNAJC6*** **(FS) H10_2** | RI | chr1:19117910-19118870 | -0.20 |
| | SE | chr1:19148626-19150575 | 0.55 |
| | composite | chr1:19139214-19143978 | -0.09 |
| ***FBXO7*** **(R498X) A7_1** | RI | chr1:19185286-19186538 | -0.32 |
| | A3SS | chr1:19155075-19155439 | -0.07 |
| ***GBA1*** **(IVS2) E10B** | SE | chr1:19148626-19150575 | 0.34 |
| ***GBA1*** **(IVS2) G2E** | SE | chr1:19148626-19150575 | 0.29 |
| ***LRRK2*** **(G2019S) 5** | RI | chr1:19160970-19161014 | 0.36 |
| ***LRRK2*** **(G2019S) 6** | SE | chr1:19148626-19150575 | 0.16 |
| ***PINK1*** **(Q129X) C4_1** | SE | chr1:19148626-19150575 | 0.45 |
| ***PINK1*** **(Q129X) E7_1** | A3SS | chr1:19145930-19146824 | 0.11 |
| ***PRKN*** **(X3DEL) H2_2** | SE | chr1:19148626-19150575 | -0.19 |
| ***SNCA*** **(A30P) A2_3** | SE | chr1:19148626-19150575 | -0.15 |
| ***SNCA*** **(A30P) E1_3** | SE | chr1:19148626-19150575 | -0.23 |
| ***SNCA*** **(A53T) 2** | SE | chr1:19148626-19150575 | 0.36 |
| | composite | chr1:19139214-19143978 | -0.05 |
| ***SNCA*** **(A53T) 3** | SE | chr1:19148626-19150575 | 0.31 |
| ***SYNJ1*** **(R258Q) A5_1** | SE | chr1:19148626-19150575 | 0.32 |
| ***VPS13C*** **(W395C) C11_3** | RI | chr1:19104666-19105046 | 0.21 |
| ***VPS13C*** **(W395C) C3_1** | RI | chr1:19104666-19105046 | 0.26 |
| | RI | chr1:19110464-19110740 | 0.17 |
| **Patient Data** | **Splicing mode** | **Genomic Coordinates** | **ΔPSI (control-PD patient)** |
| **PD patients** | A3SS | chr1:19155075-19155439 | -0.10 |
| | SE | chr1:19148626-19150575 | 0.27 |
| **DLB patients** | A3SS | chr1:19101635-19104082 | -0.06 |
| | A3SS | chr1:19155075-19155439 | -0.13 |
| | RI | chr1:19093486-19093947 | -0.24 |
| | SE | chr1:19148626_-1915057 | -0.33 |

**Supplemental Figure S15. The transcripts from the *UBR4* gene are differentially spliced in midbrain DA neuronal cells expressing familial PD mutations and in PD patient brain biopsies.** (A) Sashimi plots of the differentially spliced regions from the *UBR4* (SE: chr1:19,148,626-19,150,575) gene are shown with the number of splice junction reads indicated for the *LRRK2* (G2019S) mutant and EWT control. (B) Differential splicing events of *UBR4* gene in familial PD mutants are generated in this study and PD patient postmortem brains, including genomic coordinates and delta PSI.

**1. Autosomal Dominant Parkinson's Disease (AD-PD)**

| Gene Name | Locus | Protein Function (Key Pathway) | Clinical Features |
|---|---|---|---|
| *SNCA* | *PARK1/4* | alpha-Synuclein (Synaptic function, Lewy body formation) | Rare missense mutations (e.g., A53T, E46K) and copy number variations (duplications/triplications) cause PD. Often early-onset, rapid progression, and highly associated with dementia and widespread Lewy body pathology. |
| *LRRK2* | *PARK8* | Kinase function (Vesicular/Lysosomal trafficking) | Most common AD-PD gene. Phenotype is often clinically indistinguishable from Idiopathic PD; typically late-onset and often shows incomplete penetrance. |

**2. Autosomal Recessive Parkinson's Disease (AR-PD)**

**A. Classic AR-PD Phenotype (Typical EOPD)**

| Gene Name | Locus | Protein Function (Key Pathway) | Clinical Features |
|---|---|---|---|
| *PRKN* | *PARK2* | E3 Ubiquitin Ligase (Mitophagy) | Most common cause of EOPD. Excellent Levodopa response. |
| *PINK1* | *PARK6* | Protein Kinase (Mitophagy/Mitochondrial quality control) | Very similar to PRKN-PD; Early-Onset, slow progression. |
| *DJ1* | *PARK7* | Antioxidant/Deglycase (Oxidative stress response) | Early-Onset, Levodopa-responsive, but rare. |

**B. Atypical AR-Parkinsonism Phenotype (Parkinsonism-Plus)**

| Gene Name | Locus | Protein Function (Key Pathway) | Clinical Features |
|---|---|---|---|
| *ATP13A2* | *PARK9* | Lysosomal P-type ATPase (Metal Ion/Lysosomal Transport) | Causes Kufor-Rakeb Syndrome (KRS): Juvenile onset, pyramidal signs, cognitive impairment, and supranuclear gaze palsy. |
| *VPS13C* | *PARK23* | Lipid transfer at ER-Lysosome Contact Sites | Recessive EOPD, often with atypical features. |
| *FBXO7* | *PARK15* | Ubiquitin-Proteasome System component | Juvenile-onset parkinsonism, often with pyramidal signs and dystonia. |
| *DNAJC6* | *PARK19* | Endocytic Accessory Protein (Synaptic Vesicle Cycling) | Juvenile-onset parkinsonism, often with epilepsy or intellectual disability. |
| *SYNJ1* | *PARK20* | Phosphatase (Endocytosis/Synaptic Vesicle Cycling) | Early-Onset PD with atypical features and sometimes intractable seizures. |

**3. Genetic Risk Factor (Susceptibility Gene)**

| Gene Name | Locus | Protein Function (Key Pathway) | Clinical Features |
|---|---|---|---|
| *GBA1* | *GBA1* | Glucocerebrosidase (Lysosomal Function) | Associated Disorder: Gaucher Disease (GD)<br><br>Most common genetic risk factor for PD. Associated with earlier onset, faster progression, and higher risk of dementia. |

**Supplemental Figure S16. Genetic Architecture and Clinical Phenotypes of Parkinson's Disease.** This table summarizing key PD-associated genes categorized by inheritance pattern and clinical presentation. (1) Autosomal Dominant (AD-PD) genes (*SNCA*, *LRRK2*) require a single pathogenic allele and often resemble idiopathic PD. (2) Autosomal Recessive (AR-PD) genes require biallelic mutations and typically result in early-onset PD; these are further divided into Classic AR-PD (*PRKN, PINK1, DJ1*), showing typical motor symptoms, and Atypical AR-Parkinsonism (*ATP13A2, VPS13C, FBXO7, DNAJC6, SYNJ1*), characterized by additional neurological features. (3) Genetic Risk Factors (*GBA1*) denote variants that significantly increase disease susceptibility and influence progression of PD.
